# Perceptual maps reveal rampant convergence in butterfly wing patterns across the Neotropics

**DOI:** 10.1101/2025.02.28.640901

**Authors:** Maël Doré, Eddie Pérochon, Thomas G. Aubier, Yann Le Poul, Mathieu Joron, Marianne Elias

## Abstract

In 1879, Fritz Müller formulated the first mathematical evolutionary model to explain mutualistic mimicry between coexisting defended prey. Yet, whether local mimicry drives the structure of aposematic prey signals across their entire geographic distribution remains untested because the perception of pattern similarity has never been assessed at large spatial scale. Here, we implement a citizen science survey to map, throughout the Neotropics, variation in the wing patterns of heliconiine butterflies (Nymphalidae) as perceived by vertebrate cognitive systems. We show that despite a continuum of perceived similarity in wing patterns at the continental scale, the convergence of sympatric species into discrete mimicry rings is ubiquitous among local communities. These results expand Müller’s historical predictions by quantifying the rampant convergence of prey signals across an entire continent.

## Main Text

Prey with defenses against predators, such as unpalatability, often display conspicuous traits that act as warning signals, a phenomenon called aposematism (*1*). Defended prey from the same locality often display similar aposematic signals. This resemblance, called Müllerian mimicry (hereafter, mimicry), is an iconic example of mutualistic interactions mediated by visual signals. Following Fritz Müller’s 1879 mathematical model, the oldest for evolutionary biology (*2, 3*), selection favors the convergence of aposematic signals because the risk of attack by naive predators is spread locally over all prey individuals displaying the same signal and decreases when species mimic each other (*3, 4*). Species that evolve similar appearance under selection for resemblance are called mimicry rings (*5, 6*). Mimetic convergence is determined by how predators perceive resemblance among prey. This is influenced by the visual and cognitive abilities of predators, and by their prior experience with similar prey (*7*–*9*). Selection for resemblance operated by predators hinges on how they generalize the phenotypes of distinct prey. This can vary within and among predator communities (*10, 11*) and according to the characteristics and distribution of prey phenotypes (*12*–*14*). Therefore, perception of prey resemblance by predators represents a key component of signal evolution (*15, 16*).

Our understanding of mimicry evolution and the structure of mimetic communities is limited by the difficulty in accounting for the perceptual dimension of convergence in signal (*17*). While quantitative tests for convergence in sympatric mimetic lineages exist in the literature, they rely on discrete phenotypic classifications (*18*), statistical image processing (*19*), or image analysis using machine learning (*20, 21*). Yet, over 140 years following Müller’s equations, a quantitative test that investigates how much defended species within local communities converge in appearance remains elusive, mostly because of a lack of efficient tools to quantify perceived similarity across diverse phenotypes from a predator’s perspective.

### Testing the unfolding of Müller’s historical model of mimicry in the perceptual phenotypic space with citizen science

According to Müller’s model, phenotypic convergence between species is expected to structure the phenotypic variation of defended prey into discrete mimicry rings within a locality (*3*). At a larger spatial scale, a spatial mosaic of warning signals is expected to establish because positive frequency-dependent selection acts locally and may favor different mimicry phenotypes in neighboring communities with different species composition (*22*–*24*). However, although geographic mosaics are observed in many mimetic associations showing changes in aposematic signals every few hundred kilometers (*24*–*27*), other signals are found across entire continents, sometimes with gradual variations (*5, 6, 28*–*31*). Therefore, it is unclear to what extent mimicry, operating locally, shapes the structure of perceptual similarity in phenotype comparably across both local and continental scales (*32, 33*). In this study, we aim to elucidate how variation in aposematic signals is structured, from a perceptual standpoint, within and among communities across the entire range of diverse species groups.

We focused on the emblematic example of Müllerian mimicry in heliconiine butterflies (Nymphalidae: Heliconiini), known for their mimetic interactions and their adaptive radiation through the Neotropics (*34*). This group stands as a model for the study of natural selection, coevolution, speciation, and the genetics of adaptation (*35*–*42*), and were instrumental in the crafting of Müller’s historical mimicry model (*3*). We explored variation in wing pattern across the entire geographic range and taxonomic diversity of all 77 heliconiine species, including 432 out of 457 subspecies (*35, 43*).

To quantify similarity in wing patterns, we implemented a new method that directly integrates perception by vertebrate receivers. We designed an online citizen science survey (http://memometic.cleverapps.io/) in which participants were shown triplets of butterflies and asked to rank their similarity. We used those rankings to build multidimensional perceptual spaces in which the distance between species reflects their similarity in aposematic signals as perceived by the receivers. This new approach named perceptual phenotype mapping takes directly perception into account (*44*) while also including the multidimensional nature of aposematic signals, quantifying similarity in shapes, patterns, and colors altogether. Colors, more than patterns and shapes, tend to be primarily used by bird predators and humans to assess overall similarity (*13, 45*). By design, perceptual spaces account for the hierarchical importance of the different signal modalities by letting the structure emerge directly from the data.

To investigate variation in aposematic signals as perceived by vertebrate cognitive systems and test for signal convergence at multiple spatial scales, we built a perceptual space quantifying perceived similarity across all heliconiine butterflies. We tested whether, when all pooled together, aposematic signals converged into distinct phenotypic groups of wing color patterns in the perceptual space or were instead distributed along a continuum of phenotypes. We proof-tested the use of this global perceptual space to identify mimicry rings at the local scale within five well-documented communities. Next, we used a comprehensive collection of species distribution maps (*46*) to estimate the composition of heliconiine communities within each 30 km × 30 km grid cell across the American continent and produce a local perceptual space of species wing patterns for each community. This continental coverage allowed us to test whether the convergence of warning signals into local discrete mimicry rings is supported across all grid cells, i.e. all communities. In doing so, we provide compelling evidence for the validity of Müller’s historical model of mimicry (*3*) despite the extensive diversity and spatial variations in prey signals.

### A continuum of perceived phenotypic diversity at continental scale

In the online citizen science survey (http://memometic.cleverapps.io/), participants were iteratively presented with triplets of butterfly images drawn randomly from a pool of 432 images each representing one heliconiine subspecies. For each triplet, players had to select the most resembling image pair (*47*). In total, we collected similarity rankings on 85,320 triplets of images from 1,242 distinct participants. Demographic metadata and individual response behavior for survey participants were also recorded and analyzed in detail (see **SI Appendix 4**). To build a 3D perceptual space corresponding to the image set, we employed the t-distributed Stochastic Triplet Embedding algorithm (t-STE; *37*) optimizing image coordinates to best satisfy the similarity rankings (*47*). As a result, the relative distance of images in the perceptual space represents the aggregated perception of their similarity by the pool of participants (**Fig. 1**).

**Figure 1:**
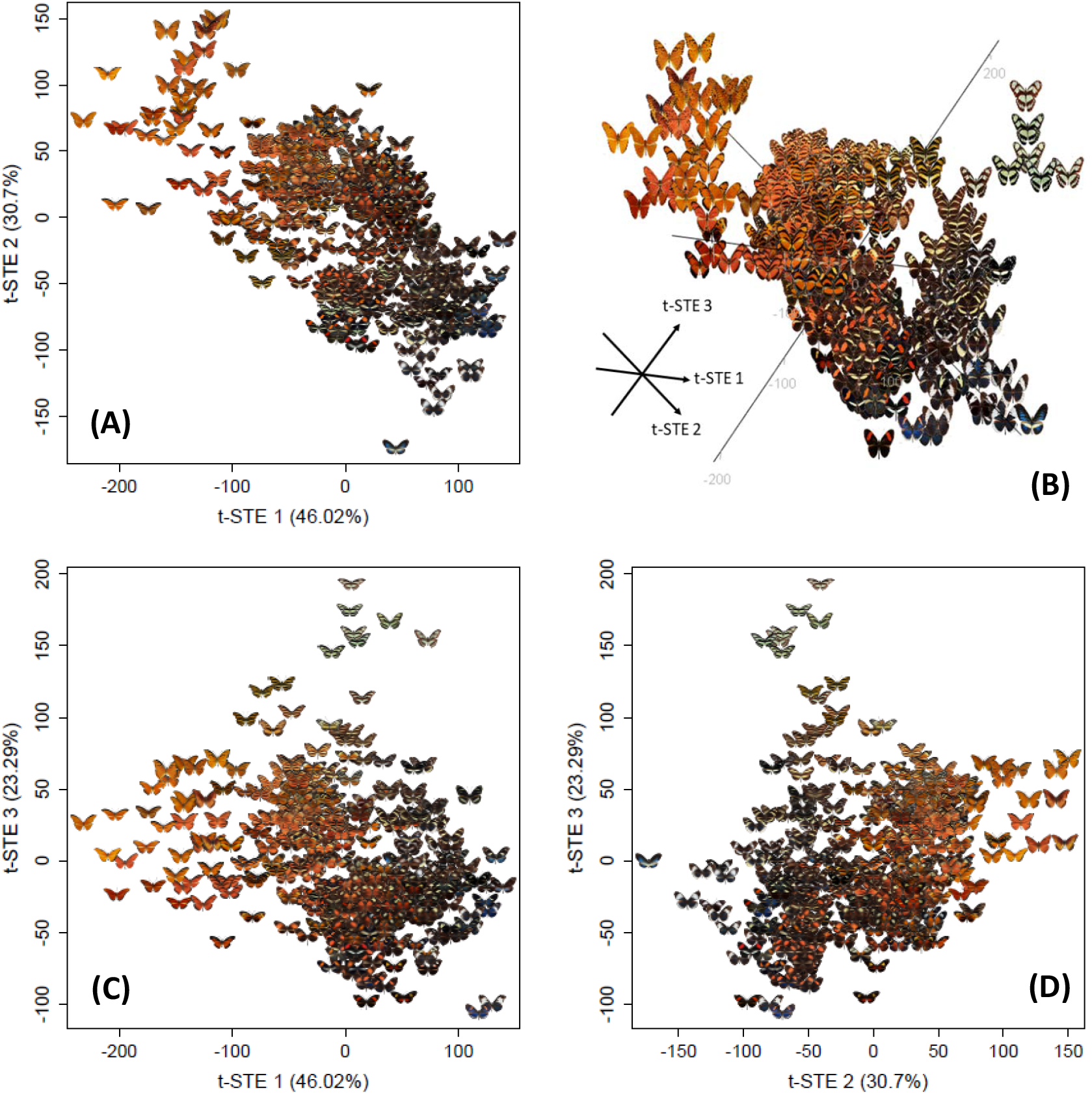
Perceptual maps of wing patterns across all heliconiine butterflies, built from similarity ranking by participants to a citizen science survey. **(A-C-D)** 2D perceptual maps. **(B)** 3D Perceptual space. All visuals display the 432 images of subspecies used during the perception survey to their t-STE coordinates in the embedded space. They encompass the whole range of heliconiine butterflies in America. Distances between patterns reflect global perception of dissimilarity. A 3D animated version of the perceptual space is available in SI for the online version.

We tested whether the distribution of wing patterns in this perceptual space, pooling all heliconiine species, complies with the expectation under a neutral model of evolution. Alternately, we examined whether it shows instead a higher degree of clustering in discrete groups of images, as expected under convergence driven by Müllerian mimicry. To this end, we computed the Perceived Phenotypic Diversity (PePD) that scores the volume occupied by phenotypes in the perceptual space based on Gaussian density kernels. The more phenotypic convergence, the less volume is occupied by the phenotypes. Comparing the observed PePD with PePD obtained from 1000 simulations of neutral evolution of wing patterns under a Brownian Motion model (*47*), we found a signal for a low level of convergence (PePD observed = 51.5; Median = 60.7; Q5% = 51.81; Z-score = −2.004; p-value = 0.045). This low level of convergence translates into an uneven continuum of perceived phenotypes (**Fig. 1**) reminiscent of the distribution of bumblebee phenotypes in the Northern American mimetic radiation (*32*). This continuum contrasts with the expectation that local mimicry rings and the associated spatial mosaic of mimetic patterns (*23*) should structure the perceptual space into clear discrete phenotypic groups. However, this apparent paradox may be explained by processes acting at multiple spatial levels.

At a local scale, species found together within a locality are expected to show significant degree of phenotypic convergence (mimicry rings) (*3*) yet they retain some degree of variation within groups (**Fig. 2**). Indeed, some predators may ignore small variations in phenotype (*49*) and slightly different patterns could provide equal protection against attacks (*50*). Ecological mechanisms leading to divergent selection such as reproductive interference may also counteract the extent of phenotypic convergence by favoring species-specific characters (*51, 52*), while developmental and genetic architecture of wing patterns may limit the possibility to evolve better resemblance (*53*). More crucially, signals may vary across regions due to genetic drift and variation in selective pressures. For instance, different suites of individual predators occurring across the landscape and exposed to different prey assemblages may select for slightly different signals (*25*). This variation in the selective optima combined with the stochastic effects of genetic drift increasing with spatial distance leads to the existence of phenotypic drift at a regional scale, even within mimicry rings (*54*).

**Figure 2:**
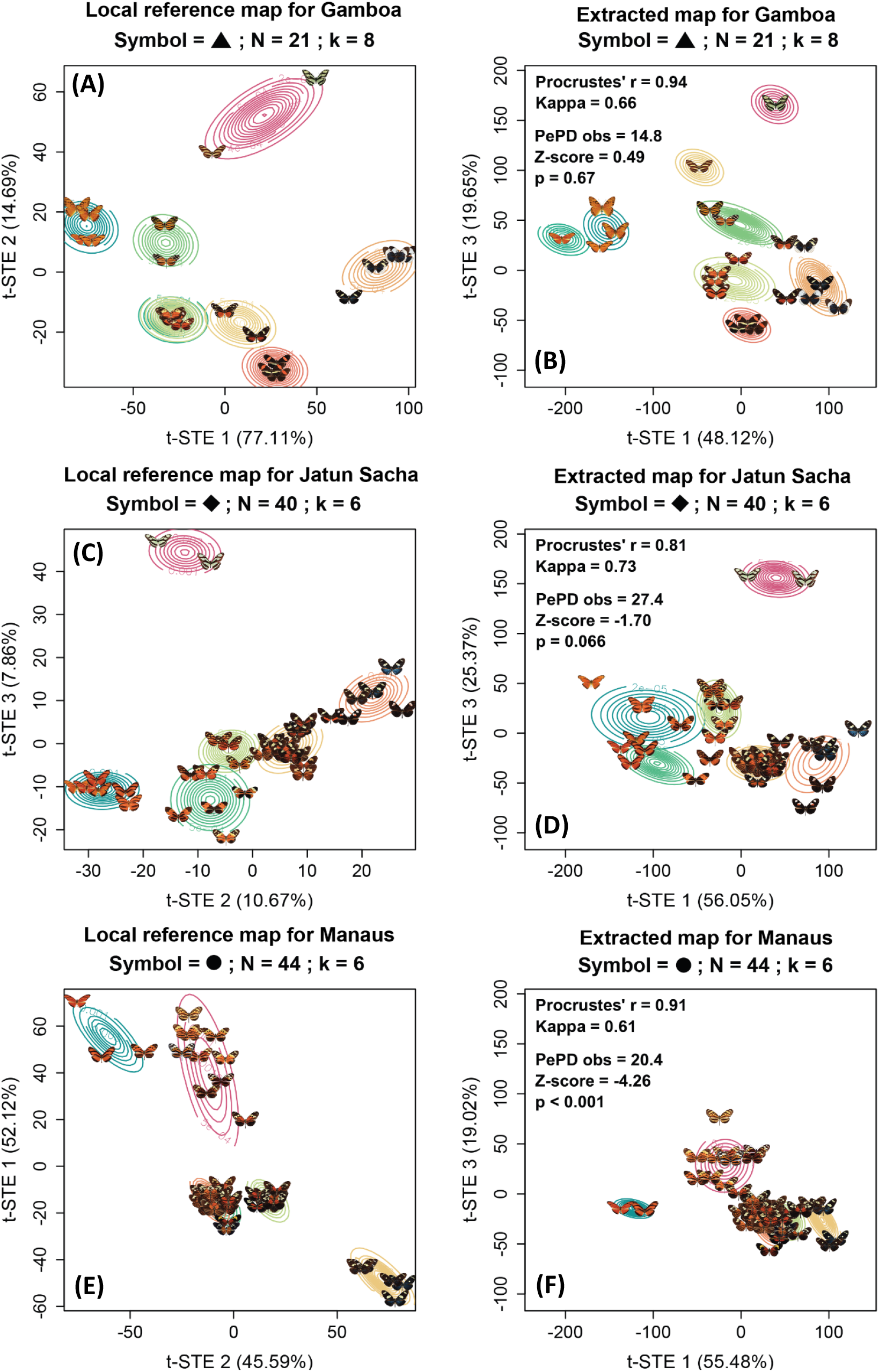
Perceptual maps of phenotypes within local communities, displayed with GMM probability densities for ‘local reference’ and ‘extracted’ maps. Contour lines represent the probability densities of GMM clusters defining local mimicry rings. Only two axes among the three are displayed. ‘Local reference’ maps **(A-C-E)** were built from similarity evaluation using triplets based only on local patterns and provided by five survey participants. ‘Extracted’ maps **(B-D-F)** were obtained from the perceptual space (**Fig. 1**) built from citizen science survey responses on triplets drawn from the entire set of Heliconiini images. Axes t-STE 1 and t-STE 3 for the ‘extracted’ maps correspond to the axes shown in **Fig. 1C**. Axes shown in ‘local reference’ maps were selected to represent similar pattern trends relative to the axes shown in ‘extracted’ maps. Procrustes correlation measures similarity in map topology between ‘extracted’ and ‘local reference’ maps. Cohen’s Kappa measures agreement in GMM classifications. Perceived Phenotypic Diversity (PePD) quantifies the coverage of local phenotypes in the perceptual space based on Gaussian density kernels, here reported as percentage of the global coverage. The Z-score compares observed PePD with PePD obtained from a model of neutral evolution of wing patterns. Negative values indicate evolutionary convergence. P-values are based on quantiles of observed values among simulated data. N = number of local patterns. k = chosen number of hypothesized mimicry rings. Maps for additional local communities of Cayenne and Santa Teresa are available in **Fig. S18** in **SI Appendix 6**. A 3D animated version of the perceptual space for ‘local reference’ of each community is available in SI for the online version.

Altogether, local imperfection in mimicry, combined with the geographic drift of selected patterns, likely contribute to forming the observed continuum of patterns in the perceptual space when all communities are pooled together at a geographic scale larger than local predator ranges (**Fig. 1**).

### Clustering of species in mimicry rings within local communities

Ecological interactions take place within local communities, where mimicry rings support mutualistic interactions associated with similarity in aposematic patterns (*55*). To explore the ability of perceptual spaces to identify local mimicry rings, we focused on five well-documented local communities and defined putative mimicry rings as clusters of wing patterns defined using Gaussian Mixture-Models (GMM; *29*) (**Fig. 2, Fig. S18** in **Appendix 6**; see **Fig. S23-S27** in **SI Appendix 10** for lists of all mimicry rings).

Real predators are only exposed to the subset of butterfly phenotypes specific to their local community. By contrast, the triplets of images proposed to our citizen science survey participants were drawn from a pool of images comprising the entire diversity of wing patterns of the continent (432 phenotypes). The perception of similarity could be influenced by this diversity (*7*). To test whether the citizen science dataset obtained with the full set of phenotypes accurately reflects perceived similarity at the local scale, we generated five local perceptual maps derived from independent evaluations to use as references. Those evaluations were carried out on wing pattern similarity solely among taxa found within each of five local communities (labeled as ‘local reference’ maps). Those maps were then compared to maps of the same five local sets of wing patterns extracted from the comprehensive perceptual space based on citizen science data (referred to as ‘extracted’ maps) (*47*).

‘Local reference’ and ‘extracted’ maps exhibited high and significant correlations in topology as quantified with Procrustes analyses (*50*; all pairwise Procrustes’ r > 0.8, all p < 0.001; **Fig. 2**; **Fig. S18** in **Appendix 6**). Furthermore, we recorded high agreement in the composition of putative mimicry rings between the two maps as assessed with Cohen’s Kappa index for classification similarity (*51*; all pairwise Cohen’s Kappa > 0.6) for all five local communities (**Fig. 2**; **Fig. S18** in **Appendix 6**). Altogether, perceptual information from the entire continental diversity can be extracted to accurately represent the perceived diversity within local communities. As such, citizen science maps (**Fig. 1)** may be coupled with species distribution data to infer community composition and describe patterns of perceived diversity across all local communities. This approach enables the comprehensive identification of local mimicry rings as described for the five case-study communities (e.g., **Fig. S23-S27** in **SI Appendix 10**). Besides, it allows for an in-depth exploration of the biogeography of this clade as it enables the mapping of perceived local diversity (**Fig. 3A**), average local phenotypes (**Fig. S19B** in **SI Appendix 7**), and phenotypic β-diversity (**Fig. S20B** in **SI Appendix 8)** across geographic space. Furthermore, it unlocks the testing of Müller’s model by investigating for local phenotype convergence while encompassing all communities at once (e.g., **Fig. 3B**; **Fig. S21 & S22** in **SI Appendix 9**).

**Figure 3:**
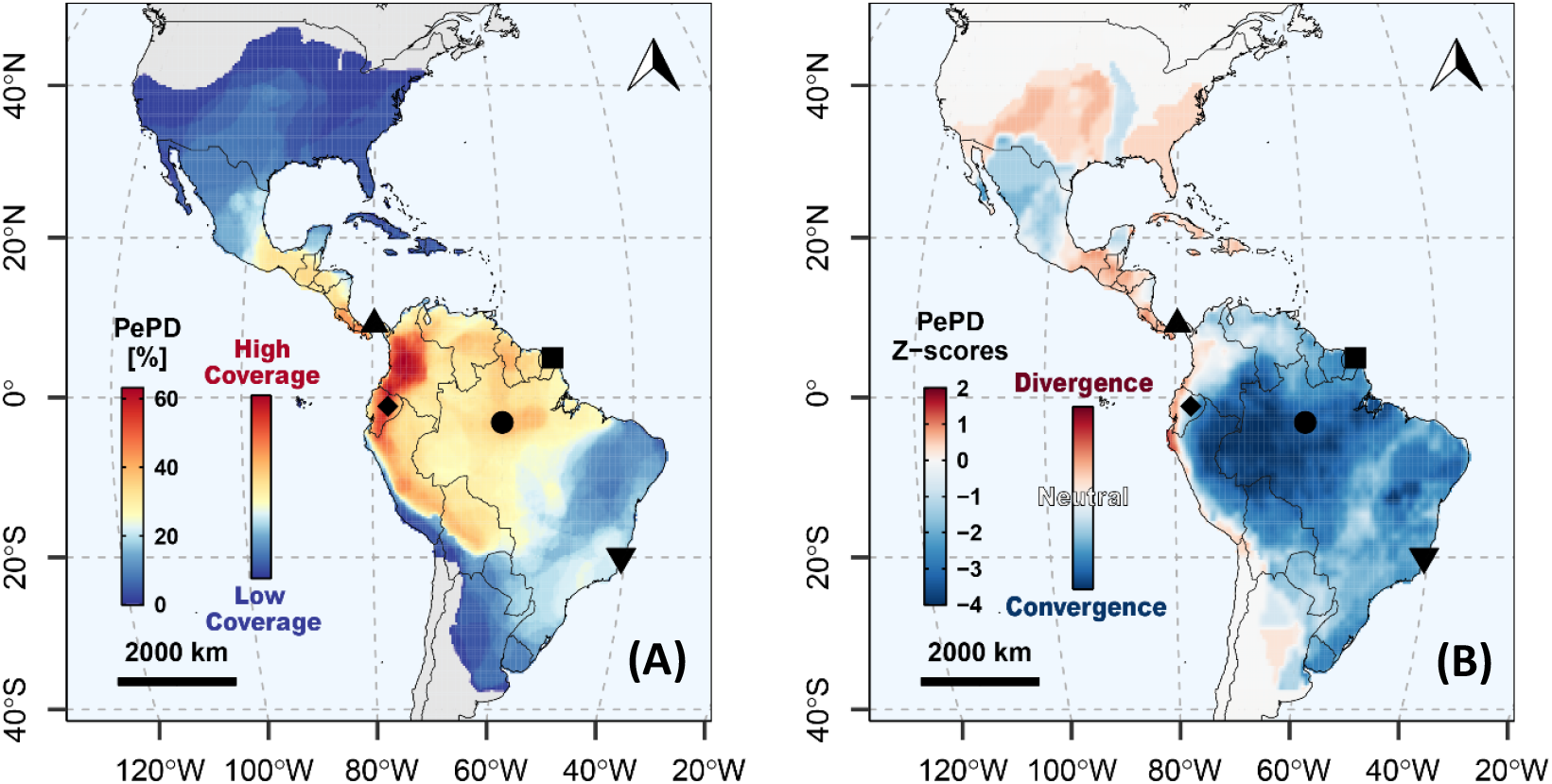
Maps of perceived phenotypic diversity (PePD) as (A) percentage of perceptual space coverage and (B) relative PePD as deviation to null model of evolution (Z-scores). PePD was computed in each grid cell (i.e., local community) as the coverage of local phenotypes in the perceptual space based on Gaussian density kernels (*47*). The more numerous and equally spread in the perceptual space are the local phenotypes, the higher the PePD index scores. (A) PePD within each local community was standardized by the global PePD computed for all communities/phenotypes pooled and represents the percentage of perceptual space occupied by local phenotypes. (B) Relative PePD was evaluated by comparing the observed PePD with PePD obtained from 1000 simulations of neutral evolution of wing patterns. An observed PePD lower than expected from neutral evolution (i.e., Z-score < −1.96; in blue) illustrates the evolutionary convergence of local phenotypes. Conversely, an observed PePD higher than expected from neutral evolution (i.e., Z-score > 1.96; in red) illustrates the evolutionary divergence among local phenotypes. Perceptual maps of the local communities symbolized with shapes are presented in **Figure 2**: square = Cayenne, up-triangle = Gamboa, diamond = Jatun Sacha, circle = Manaus, down-triangle = Santa Teresa.

### Macroecological patterns of perceived phenotypic diversity in the Neotropics

Müller’s model of mimicry predicts that sympatric species should exhibit convergence of wing patterns in the eyes of the predators. We tested this prediction quantitatively across all communities of the Neotropical area, by combining the perceptual maps from the citizen science survey (**Fig. 1)** and geographic maps of spatial distributions for heliconiine species (*59*). In each grid cell/community (ca. 30 km × 30 km), we compared the observed local perceived phenotypic diversity (PePD) to the PePD obtained from 1000 simulations of neutral evolution of wing patterns along the phylogeny of Heliconiini (*43*), thus assessing the degree of relative space coverage of local phenotypes (*47*). We provide perceptual evidence that convergence of aposematic signals is the predominant pattern in heliconiine communities across most of the distribution range of Heliconiini (**Fig. 3B**; Z-scores < −1.96). Perceived phenotypic diversity is highest in the Andes and Central America (**Fig. 3A**), yet local phenotypic clusters in those communities were compatible with a neutral model of wing pattern evolution (**Fig. 3B**; |Z-scores| < 1.96). Such pattern arises in those regions because co-mimics are often closely related species as illustrated in the Gamboa (**Fig. 2B**, upward triangle symbol; PePD observed = 14.8, Z-score = 0.49, p-value = 0.67) and Jatun Sacha communities (**Fig. 2D**, diamond symbol; PePD observed = 27.4, Z-score = −1.70, p-value = 0.066). By contrast, the lower phenotypic diversity found in the Amazon basin highlights a clustering into fewer mimicry rings with overall lower phenotypic coverage than in the Andes. In the Amazon basin, rings are composed of numerous distantly related species converging towards similar wing patterns, as illustrated by the Manaus community (**Fig. 2F**, circle symbol; PePD observed = 20.4, Z-score = −4.26, p-value < 0.001). In contrast to the steeper environmental gradients in the Andes (*60, 61*), environmental and topographical homogeneity across the Amazon basin (*62*) may limit the structuring of mimetic communities, fostering the accumulation of species in a smaller number of large and widely distributed mimicry rings across the Amazon region (*46*).

### Large-scale convergence of sympatric species in the perceptual space

While the co-occurrence of species with similar wing phenotypes is well-established when species are assigned by experts to discrete mimicry groups (*6, 18, 63*), our study shows that Müller’s prediction for local convergence in aposematic signals among species is supported when resemblance is quantified in a continuous perceptual space, without having to define a priori mimicry groups. Beyond its effect on the spatial heterogeneity of phenotypic variation (**Fig. 3B**), Müllerian mimicry predicts selection on resemblance between coexisting taxa (*3, 4*). To test this prediction in an evolutionary framework, we examined whether species with overlapping spatial distributions present a higher perceived phenotypic similarity in wing pattern than expected from their evolutionary relatedness (*47*). As predicted, perceptual distances between pairs of subspecies were significantly correlated with their geographic distances estimated through range overlaps (Multiple Regression on distance Matrices (MRM): standardized β_obs_ = 0.078, Q95% = 0.040, p = 0.003; **Figs. S21A & S21B** in **Appendix 9**), supporting the idea that co-occurring species display similar appearance. This correlation holds when accounting for phylogenetic distances across taxa, revealing convergent evolution beyond the effect of shared ancestry (MRM: standardized β_obs_ = 0.085, Q95% = 0.039, p ≤ 0.001; **Figs. S21C & S21D** in **Appendix 9**). Similar results were obtained when categorizing spatial distances between pairs of species as sympatric or allopatric (See **Fig. S22** in **SI Appendix 9**).

Altogether, amidst a wealth of phenotypic variation within and between local mimetic communities (**Fig. 2**), and despite a continuum of perceived phenotypes appearing at a large geographic scale (**Fig. 1**), our study highlights how Müllerian mimicry is a key driver of convergence in perceived wing patterns between coexisting heliconiine species across the American continent, especially in the Amazon basin (**Fig. 3B**). Therefore, our findings support the extension of the predictions from Müller’s historical model to a large spatial scale using a quantitative description of pattern variation. Crucially, the perceptual approach developed here accounts altogether for morphological and chromatic dimensions of trait variation, and also integrates cognition processes in the selective agents.

### Limits and perspectives of perceptual phenotype mapping for ecological signals

Our study aims at studying resemblance in butterfly wing patterns at the global and local scales focusing on how vertebrate cognitive systems perceive them. Neotropical butterflies face diurnal predation primarily from birds, especially those specialized in capturing insects in flight, such as jacamars (*64*). Birds and humans differ in their visual perception in some respects (*65*– *68*), yet humans are able to detect and describe butterfly mimicry, which suggests that human cognitive processing of visual signals must align sufficiently closely with those of the predators causing such mimicry (*55*). Evidence also supports a higher contrast sensibility and visual acuity of humans compared to most bird species (*69, 70*), powering human abilities in discrimination tasks with visual signals such as mimicry patterns (*71*). For instance, humans rank hoverfly mimics similarly to pigeons (*49*), while humans and great tits also show quantitatively similar abilities to learn and generalize warning signals (*72*). Therefore, quantifying mimetic similarity using human perception can provide valuable information on the rules underlying phenotypic convergence in prey signals in nature (*71*).

Perceptual phenotype mapping allows assessing variation in phenotypes taking into account morphological, chromatic, and cognitive dimensions simultaneously, which is key when studying variation in signals linked to fitness via a receiver’s perception. It also offers a practical solution to quantify complex patterns that image analysis struggles with due to simultaneous variation in shape, color, and pattern. Instead of simplifying color patterns in standardized templates (*32, 73*), decomposing complex signals into separate modalities (*74*), or relying on taxonomic classification for supervision (*20*), perceptual phenotype mapping can quantify any trait variation as long as relative perceived similarity between triplets of objects can be informed. This innovative and versatile method significantly enriches the ecologist’s and evolutionist’s toolbox to study ecological trait variation. The perceptual approach unlocked our ability to investigate wing pattern variation in heliconiine butterfly from local community to a whole continent, for the entire clade. Overall, it offers new perspectives for the investigation of phenotypic variation at multiple geographic and phylogenetic scales and may stimulate further studies on the evolution of traits whose complexity has prevented large scale comparison until now.

## Acknowledgements

We thank all citizen scientists who participated in our online survey making this study possible. We thank museum curators and expert lepidopterists who provided access to collections, agreed to share images, or helped to curate taxonomic lists including Keith Willmott (FMNH), Augusto H.B. Rosa and André V.L. Freitas (UniCamp), Rodolphe Rougerie and Jérôme Barbut (MNHN), Blanca Huertas and Robyn Crowther (NHMUK), Chris Jiggins, Pierre Boyer, Michel Cast, Gerardo Lamas, Andrew D. Warren, Daniel H. Janzen, Jim P. Brock, Kim Garwood, Luis Miguel Constantino, Christian Brévignon, Rémi Mauxion, Ombeline Sculfort, Olivier Claessens, Leila Teruko Shirai, and Kelve Franklimara de Sousa Cézar.

## Author contributions

Conceptualization: MD, EP, TA, YLP, MJ, ME

Data curation: MD, EP

Formal Analysis: MD

Software: MD, EP, TA

Visualization: MD

Writing – original draft: MD

Writing – review & editing: MD, EP, TA, YLP, MJ, ME

## Data and materials availability

All MATLAB and R scripts to carry out analyses are available on GitHub (link to be released upon publication). All 2D perceptual maps, 3D animated perceptual spaces, mimicry ring lists and subspecies images used in the online survey are available in online archives in Zenodo (https://doi.org/10.5281/zenodo.10076355). The online survey for the citizen science data collection is temporarily available on http://memometic.cleverapps.io/ with source code accessible on GitHub (link to be released upon publication). Heliconiine distribution maps were retrieved from https://doi.org/10.5281/zenodo.10903661.

## Supplementary Materials

Supplementary Materials includes:

- Supporting Information for Materials and Methods with Figure **SMM1**.
- Supporting Information for Appendices 1 to 10 with Figures **S1** to **S27 & Table S1**.
- **References 75 to 83** are only cited in the SM.
- **Movies S1-S6**: Six animated 3D versions of perceptual spaces of heliconiine butterflies, one for the global Citizen Science dataset, and one for each of the five local reference communities.

