## Supplementary Materials for "Perceptual maps reveal rampant convergence in butterfly wing patterns across the Neotropics"

**[†]Co-last authors**

***Corresponding author:** Maël Doré

**Address:**

Muséum national d'Histoire naturelle

45 rue Buffon, CP50

75005 Paris, France

**Phone number:** +33-1-40-79-37-73

**This PDF file includes:**

- Supporting Information for Materials and Methods with Figure **SMM1**.
- Supporting Information for Appendices 1 to 10 with Figures **S1** to **S27** & **Table S1**.

### Supporting Information for Materials & Methods

#### 1 Materials

##### 1.1 Study group

We focused our study on the textbook case of Müllerian mimicry in heliconiine butterflies (Nymphalidae: Heliconiini). This group is well-known for the important variation of its aposematic patterns (*35*) and for the existence of local geographic forms reflecting parallel adaptive radiations between species (*75*). Heliconiines are widely distributed across the American continent, from Canada to the North of Argentina with peaks of diversity in the Andes, in Central America, and in the Amazon basin (*36*, *46*). The study of wing patterns of heliconiines, among other groups of mimetic neotropical butterflies, has been instrumental in the emergence of the theory of mimicry in the late nineteenth century, notably by Henry Walter Bates (*76*) and Fritz Müller (*3*) who gave their names to the two major types of mimicry: Batesian mimicry and Müllerian mimicry. All species of heliconiine butterflies are engaged in Müllerian mimicry (*35*) which implies some degree of toxicity associated with conspicuous warning signals displayed by wing patterns. Some species share mimicry patterns with other groups of neotropical mimetic Lepidoptera, mostly ithomiine butterflies (*46*, *77–79*).

##### 1.2 Image collection

We retrieved standardized images of dorsal view of specimens in mounting position from museum and private collections for 432 subspecies (94.5%) of heliconiine butterflies among the 457 described subspecies. The most important contributors were the McGuire Center for Lepidoptera and Biodiversity (Gainesville, FL, USA), the Natural History Museum London (UK), and the Muséum National d'Histoire Naturelle (Paris, France). Due to the diversity of sources needed to retrieve such a collection of images, we included whenever possible a color reference chart on images. We standardize all images to achieve similarity in brightness, hue and saturation when comparing the color reference charts between images.

Polymorphism within subspecies is rare since subspecies are typically described on the basis of phenotypic variation in wing patterns. However, in the rare cases of polymorphism we encountered, we selected a specimen for which we considered the pattern to be the most representative (the closest to an "average" phenotype). Heliconiine butterflies notably lack sexual dimorphism in wing patterns (*19*), thus we selected indifferently male or female specimens according to availability. A complete list of taxa used for this study, with associated metadata is available in **Table S1** in **SI Appendix 1**.

##### 1.3 Online citizen science survey

We designed a website interface introduced as a game dedicated to the collection of information regarding the perception of similarity in wing patterns across the 432 subspecies in our image database. The website is accessible in four languages (i.e., English, French, Spanish, Brazilian/Portuguese) to encourage people from all around the world, and especially in regions where heliconiine butterflies are native, to participate. It is temporarily available at http://memometic.cleverapps.io/ with source code accessible on GitHub (link to be released upon publication)

This website is presented as a citizen science project with three goals: (1) collecting an important and diverse amount of data regarding the global perception of similarity in the wing patterns of heliconiine butterflies; (2) introducing a wide public to the basis of ecological and evolutionary concepts involved in Müllerian mimicry by offering a short introduction to those concepts; (3) involving citizens around the world into the collective building of scientific knowledge, and allowing them to access preview of new research development by providing feedback on the results acquired thanks to their contribution.

A typical game session involved 30 random sets of triplets of images drawn from the set of 432 subspecies images. For each triplet, we asked the player to select the pair of images perceived as the most similar (see **Fig. S1** in **SI Appendix 2** for an illustration of the website interface). Therefore, for each vote the player provided a set of relative distances. For instance, if the player selected the pair A-B as being the most similar images from the triplet, the following relative distances were recorded: $d_{A\text{-}B} < d_{A\text{-}C}$ and $d_{B\text{-}A} < d_{B\text{-}C}$ since distances are symmetrical ( $d_{A\text{-}B} = d_{B\text{-}A}$ ). In case of doubt, the player was allowed to skip the triplet and request a new one to evaluate, avoiding accumulation of noise in the data due to forced random choices. Each game session included two triplets drawn from a set of control trials for which the perceived relative similarity was striking (see **Fig. S2** in **SI Appendix 3** for visual illustration of the controlled trials used). Thus, we were able to filter game sessions that provided suspicious results by removing those that failed at least one control trial.

Before each game, demographic information (age, scientific background, colorblindness, experience with the game, language, and location), and game statistics (time taken per triplets, number of skips, score as a deviation to the consensus) were recorded. All this information is summarized and analyzed in the **SI Appendix 4**.

Altogether, we recorded 1,422 game sessions from 1,242 distinct players across 53 countries providing 85,320 triplets of relative distances. We kept 75,240 for downstream analyses after filtering of improper game sessions having failed controlled trials or declared colorblindness.

##### 1.4 Spatial distributions

We retrieved predictions of spatial distributions for 439 subspecies from recently made available maps (*54*; https://doi.org/10.5281/zenodo.10903661) built upon a dataset of 77,577 georeferenced occurrences and employed species distributions models to map subspecies distributions at the continental scale for a resolution of 0.25° arc (ca. 30 km). Maps of phenotypic diversity (**Fig. 3**), community mean phenotype (**Fig. S19B** in **SI Appendix 7**), and phenotypic β-diversity (**Fig. S20B** in **SI Appendix 8)** encompass the 422 (92.3%) subspecies for which we had both reference image for the survey and distribution patterns available.

##### 1.5 Phylogenetic relationships

We employed the phylogeny of the tribe Heliconiini in Kozak et al. (*43*) encompassing 67 of the 77 recognized species (87 %) to carry out all phylogenetic comparative analyses. We grafted the 412 associated subspecies (95.3% of subspecies depicted in our perceptual spaces) to their respective tip species using branches of null length as the evolutionary relationships at intra-specific levels are mostly unknown in the tribe. All tests for evolutionary convergence were carried out with this grafted subspecies-level phylogeny.

##### 1.6 References for local communities

To assess the suitability of the citizen science dataset to inform on local patterns of phenotypic diversity, and provide identification for local mimicry rings, we compared local maps extracted from the citizen science-based maps (identified as ‘extracted’ maps) with ‘local reference’ maps. The latter were based on similarity rankings provided by an independent panel of five participants who assessed pattern similarity solely among taxa found within local communities (identified as ‘local reference’ maps). We produced such reference comparison analyses for a set of five local communities selected among the 23,661 available communities defined as a unique grid-cell in the 30 km × 30 km raster maps of subspecies distributions, namely Cayenne, French Guyana (4°56'06"N, 52°19'16"E), Gamboa, Panama (9°07'12"N, 79°42'18"W), Jatun Sacha, Napo, Ecuador (1°05'16"S, 77°36'58"N), Manaus, Amazonas, Brazil (3°05'53"S, 60°03'32"W), and Santa Teresa, Espírito Santo, Brazil (19°55'55"S, 40°36'00"W). We extracted for each one a list of taxa whose presence was predicted by the spatial distributions obtained from Pérochon *et al.*, 2025 (*46*). This selection was made in order to (i) present a diversity of geographic location across the heliconiine range, (ii) display a diversity of phenotypes, (iii) display a diversity of level of richness for local species and mimicry patterns. The geographic location of each local community is featured on the maps of phenotypic diversity (**Fig. 1**), local mean phenotype (**Fig. S19** in **SI Appendix 7**), and phenotypic β-diversity (**Fig. S20** in **SI Appendix 8**). A visual list of subspecies for each local community can be found in **Figs. S13-S17** in **SI Appendix 5**.

We generated sets of 600 triplets of images of local taxa from each local community. Each participant evaluated a set of 600 triplets for each community using the same interface as the online citizen science survey thus producing sets of perceived relative distances across images. These sets were aggregated by community and used to generate the ‘local reference’ maps described in subsequent analyses. As such, compared to the ‘extracted’ maps, the ‘local reference’ maps are generated by focusing only on the phenotypes available in each local community.

#### 2 Methods

##### 2.1 t-STE

To carry out perceptual phenotype mapping and convert the lists of triplets of relative distances in maps of wing pattern (dis)similarity based on perception (i.e., perceptual maps), we adapted the t-distributed Stochastic Triplet Embedding method (t-STE; *20*). t-STE is a supervised contrastive machine learning algorithm based on similarity triplets under the form “A is more similar to B than to C” that produces an embedding of data whose coordinates in the new reduced space agree with the relative distances described in the similarity triplets. In the context of the perceptual map, triplets of perceived relative distances among images are converted into coordinates in a perceptual space where distances depict perception of (dis)similarity among the images.

The algorithm starts with a random set of coordinates for each object in a reduced space of user-defined number of dimensions (three dimensions in our case, see below). At each step, it computes the likelihood for each triplet (the data) to be satisfied given the current coordinates of the images (the model). The cost function is the log-likelihood of the current embedded space defined as the sum of the log-likelihood of each triplet computed as follows:

$$L_{abc} = \frac{\left(1+\frac{\|X_a - X_b\|^2}{\alpha}\right)^{-\frac{\alpha+1}{2}}}{\left(1+\frac{\|X_a - X_b\|^2}{\alpha}\right)^{-\frac{\alpha+1}{2}} + \left(1+\frac{\|X_a - X_c\|^2}{\alpha}\right)^{-\frac{\alpha+1}{2}}} \quad \text{(Eqn. 1)}$$

where $L_{abc}$ is the likelihood of the triplet of images A, B, and C, following X, their respective coordinates in the embedded space, and adjusted by α, the degree of freedom of the Student-t kernel equal to the requested final number of dimensions minus one (*48*). As such, the larger the Euclidean distance between A and C, compared to A and B, the higher will be the likelihood of the triplet, and conversely. Following a gradient-descent optimization, the coordinates of images in the embedded space are modified in order to maximize the cost function at each step. Therefore, the algorithm learns iteratively the best embedding to optimize the distances of images in the reduced space satisfying the initial triplets.

t-STE allows the user to define a priori the final number of dimensions of the embedding. A higher number of dimensions offers more possibilities to satisfy triplet similarities, but it also prevents easy visualization of the output, and limits performance of clustering algorithms in high dimensionality (*80*). With the objective of quantifying and visualizing the variation in wing patterns while obtaining hypotheses for phenotypic-based mimicry rings issued from clustering, we favored the use of three-dimensional perceptual spaces as our targeted number of dimensions.

##### 2.2 Gaussian Mixture Models and putative mimicry rings

To propose hypotheses for local mimicry rings, we ran Gaussian mixture-models (GMM; *29*) to cluster wing patterns within groups of phenotypically similar patterns. Any clustering method could potentially be applied on the perceptual space to define such hypotheses for mimicry rings, however Gaussian mixture-models offer a couple of advantages. Clusters represent Gaussian distributions that can have different sizes (i.e., number of objects), ellipsoidal shapes, and orientations. It allows us to detect outliers by forming singletons (i.e., group of a single object). In the context of mimicry, the underlying Gaussian distributions can be seen as selective values in an adaptive landscape describing an adaptive peak for phenotypes (i.e., learned by local predators as associated with toxicity, thus favored by selection) as a high probability density. If known, the local abundances of taxa can be used to provide weights influencing the clustering outcome. Finally, it allows us to fit a model with any given number of groups/mimicry rings and to compare them with criteria for goodness-of-fit (e.g., likelihood, BIC, AICc). This versatility allows the researcher to define a priori its final number of groups according to a required degree of generalization/refinement for the phenotypic groups, or to compare results and select the appropriate value according to a chosen criterion. In this study, we defined a specific number of local clusters for each community based on how well it depicted hypothesized mimetic relationships. For statistical analyses, we used an entire range of plausible number of clusters from 5 to 10 groups.

##### 2.3 Map variation of phenotypic patterns at large-scale

Employing the t-STE algorithm on the similarity triplets gathered from our citizen science collection of perception data, we obtained a 3D perceptual space with coordinates of images representing the aggregated perception of wing pattern similarity in heliconiine butterflies for patterns found on the entire range of the group (**Fig. 1B**). We decomposed this 3D perceptual space into 2D perceptual maps and displayed the images used during the survey on their given coordinates (**Figs. 1A, 1C, 1D**).

We combined the coordinates of (sub)species phenotypes in the perceptual space with predicted distribution data of each subspecies to map the mean local phenotype and compute two indices of mimicry diversity across all local communities (i.e., pixel in the grid-cell).

By converting each axis on a 0-255 scale, we assigned an RGB color to each image based on its coordinates in the perceptual space. We computed the mean local phenotype within each community and displayed its associated color on the RGB space decomposed in 2D maps and in the geographic space (see **Fig. S19 in SI Appendix 7**).

To describe the prevalence of phenotypic clusters, we quantified perceived phenotypic diversity (PePD) as the coverage of subspecies phenotypes in the perceptual space (**Fig. 3A**). First, we gridded the 3D perceptual space and estimated phenotype probability density in each bin using Gaussian kernel density estimates (KDE) to smooth the distribution of phenotypes. We used the Silverman's estimator (*81*) to estimate the kernel bandwidths with all subspecies included, and used the calculated bandwidths for all subsequent density estimations. We computed a KDE for each phenotype independently. Next, probability densities of all phenotypes were combined to extract the maximum probability to find any phenotype in each bin of the gridded perceptual space. As such, using the maximum densities, we built an estimator that discards the influence of phenotypes with overlapping density distributions. PePD was computed as the sum of those maximum densities across all bins of the gridded space. Since the sum of any individual phenotype KDE across all bins is one, the theoretical maximum PePD for a given set of phenotypes occurs when the individual phenotype KDEs do not overlap leading to a PePD equal to the number of phenotypes. Therefore, the more numerous and equally spread in the perceptual space are the phenotypes, the higher and closer to the species richness the PePD index will score. Conversely, if phenotypes are highly similar thus clustered, the PePD will be low, with a theoretical minimum of one when the coordinates of all phenotypes in the perceptual space are equals and all density kernels are perfectly overlapping. PePD can be computed for any set of phenotypes, thereby it can quantify local phenotypic diversity within communities as well as the global phenotypic diversity perceived when all phenotypes are pooled. We divided the observed PePD in local communities by the global PePD to compute the relative PePD representing the proportion of the global perceptual space occupied by local phenotypes found in each community (**Fig. 3A**).

Finally, we estimated phenotypic β-diversity between all pairs of local communities as the Mahalanobis distances between the coordinates of their respective local phenotypes in the global perceptual space (**Fig. 1**). To obtain a visual representation of β-diversity, we applied Non-metric MultiDimensional Scaling (NMDS) on the matrix of pairwise Mahalanobis distances to build a new 3D space where distances between communities relate to their dissimilarities in phenotypic composition as estimated from Mahalanobis distances. We converted this perceived phenotypic composition space into an RGB color space with dimensions ranging from 0 to 255 and assigned a color to each community based on its coordinates in the RGB space. We displayed communities with their associated color on the RGB space decomposed in 2D maps and in the geographic space (see **Fig. S20 in SI Appendix 8**).

#### 3 Statistical Analyses

##### 3.1 Comparing extracted citizen science-based maps *vs.* local reference maps for local communities

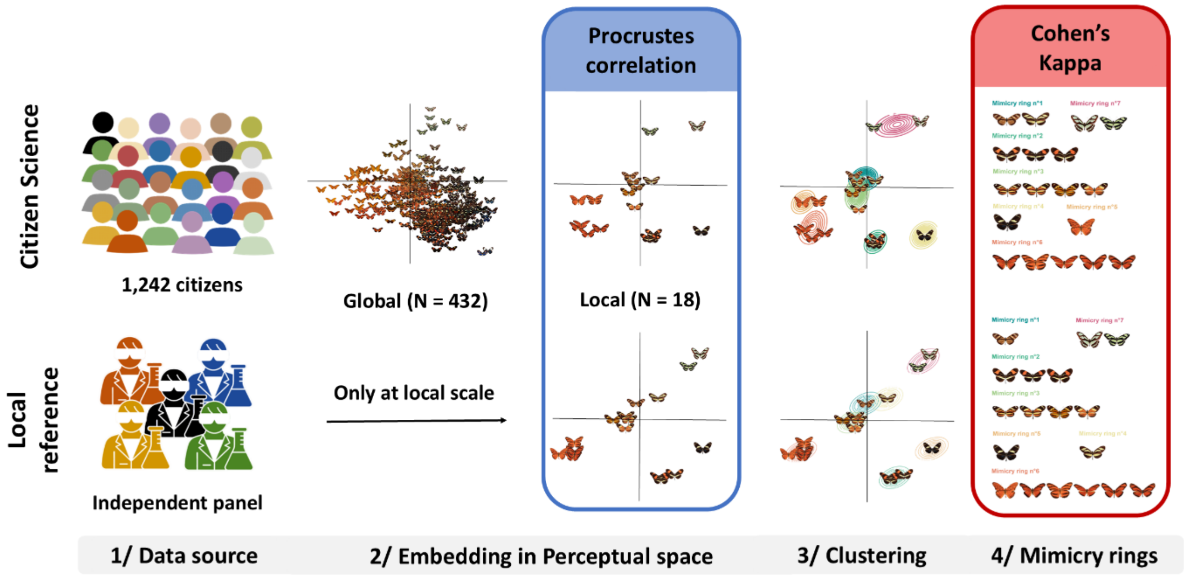


**Figure SMM1: Framework used to assess the ability of the citizen science dataset encompassing all 432 subspecies to inform on local patterns of phenotypic diversity and provide identification for local mimicry rings.** Triplets of relative perceived distances across butterfly images are gathered independently from the citizen science survey encompassing all 432 subspecies and an independent panel focusing on local phenotypes only (**Step 1**). Topology of the 3D perceptual spaces obtained from t-STE (**Step 2**) are compared with Procrustes correlation (**in blue**). Compositions of local mimicry rings obtained from clustering (**Step 3**) are compared with Cohen's Kappa (**Step 4, in red**).

To assess the suitability of the citizen science approach to inform on local patterns of phenotypic diversity, and provide suitable identification for local mimicry rings, we compared local maps extracted from the citizen science-based maps (i.e., 'extracted' maps) with reference maps based on the aggregated perception of an independent panel of participants evaluating only the patterns of local taxa (i.e., 'local reference' maps) (**Fig. SMM1**).

First, to produce the 'extracted' maps, we extracted from the macro-scale citizen science-based perceptual space the coordinates of subspecies listed in each local community to create a local perceptual space for each one. As such, these perceptual spaces have been generated with low sampling effort from thousands of players on the basis of comparison of all 432 phenotypes, not only the local ones.

Second, to produce the 'local reference' maps used as the best available standard for comparison, we ran t-STE on the aggregated sets of triplets of local patterns evaluated independently by a panel of five people (600 triplets per person per community). As such, these perceptual spaces have been generated using only local phenotypes.

As a visual comparison, we plotted the 'extracted' maps *vs.* the 'local reference' maps for each local community, displaying probability densities associated to GMM clustering applied on those perceptual maps (**Fig. 2**; **Fig. S18** in **Appendix 6**).

Additionally, we compared the topology of the 'extracted' maps and the 'local reference' maps using the Procrustes correlation statistics obtained from pairwise Procrustes Analyses (*57*). Such analyses minimize the distances across homologous points in two topologies by applying scaling, translation, and rotation actions on the initial coordinates. Similarly to a Pearson coefficient, the Procrustes correlation is computed as follows:

$$Procrustes'\, r = \sqrt{1 - \mathrm{SS}} \qquad \text{(Eqn. 3)}$$

where *SS* is the sum of squared distances between Procrustes adjusted coordinates of two maps scaled so that the maximum value is 1. We carried out tests for the significance of each pairwise Procrustes correlation based on the random permutation of coordinates across subspecies.

Furthermore, we compared the putative mimicry rings obtained from GMM clustering applied on each 'local reference' map *vs.* 'extracted' map using the Cohen's Kappa index (*58*). This index is a measurement of agreement between multinomial classifications. It allows to quantify the degree of similarity between two classifications where groups are not necessarily paired across classifications (i.e., mimicry rings do not have to be equivalent between the two classifications). In our case, it was computed from a co-membership confusion matrix summarizing the number of pairs of objects found in a similar/different group in one or the two classifications. The more the two classifications agreed on whether a pair of objects (e.g., butterfly wing patterns) should belong to the same or to a different group, the higher the Cohen's Kappa was. A Cohen's Kappa close to zero reflected a random agreement between the classifications. A Kappa higher than 0.5 reflected relatively important agreement. A Kappa close to 1 reflected an agreement close to perfection. Thus, we computed the Cohen's Kappa between the classification obtained from the citizen science-based 'extracted' maps *vs.* the 'local reference' maps, for each local community.

##### 3.2 Tests for convergence in the perceptual space

Müller's theory of mimicry (*3*) predicted the convergence of local toxic prey towards a similar aposematic pattern. Building upon our citizen science-based perceptual space (shown in **Fig. 1**), predictions for species distributions (*46*), and species-level phylogenetic tree (*43*), we tested for the convergence of local phenotypes in the perceptual space across all local communities (i.e., 30 km x 30 km grid cells). We compared the perceived phenotypic diversity (PePD) observed globally (i.e., including all phenotypes) and within each local community, with those obtained from 1000 simulations of neutral evolution of wing patterns to assess the relative degree of perceptual space coverage of local phenotypes (**Fig. 3B**; **Fig. S21** in **SI Appendix 9**).

We used the best fitted macroevolutionary model to simulate the multivariate evolution of perceived wing patterns: a BM with a Pagel's lambda = 0.891 to account for the strength of the phylogenetic signal. PePD was computed globally (i.e., including all phenotypes) and across each local community (i.e., including only local phenotypes) for each simulated trait dataset. Then, we computed Z-scores such as $Z = \frac{X_{obs} - \bar{X}}{sd(X)}$ where $X_{obs}$ is the observed diversity index, and $\bar{X}$ and *sd(X)* the mean and standard deviation across simulated datasets. An observed PePD lower than expected from neutral evolution (i.e., Z-score < 0) illustrates the evolutionary convergence of phenotypes. Conversely, an observed PePD higher than expected from neutral evolution (i.e., Z-score > 0) illustrates the evolutionary divergence of phenotypes in the perceptual space. For a significant threshold of $\alpha = 0.05$, $Z$-scores are considered significant when $|Z| > 1.96$ assuming a normal distribution of the simulated data. However, to account for potential non-normal distributions, we quantified in parallel p-values as the quantile of the observed data among simulated data.

##### 3.3 Tests for relationships between perceived phenotypic similarity and geographic distances

To evaluate if Müller's prediction for local convergence of toxic prey could influence large-scale patterns of species and phenotypic distributions, we tested if the perceptual distances between subspecies were correlated with the geographic distances between their spatial distributions. We defined pairwise perceptual distances as the Euclidean distances between pairs of subspecies in the perceptual space (shown in **Fig. 1**). We computed pairwise geographic distances as 1 - Schoener's D (*82*) in the geographic space as follows:

$$1 - D_{12} = \frac{1}{2}\sum_{ij}\left|z_{1ij} - z_{2ij}\right| \qquad \text{(Eqn. 2)}$$

where $D_{12}$ is Schoener's D between taxa 1 and taxa 2, *ij* is a cell in the geographic raster grid, $z_{1ij}$ the occupancy of taxa 1 in cell *ij*, and $z_{2ij}$ the occupancy of taxa 1 in cell *ij* with the sum of occupancy data for each entity summing to 1. As such, a distance of 0 relates to a pair of taxa with the exact same geographic distribution, a distance of 0.8 relates to a weak spatial overlap, and a distance of 1 relates to no overlap.

We used Multiple Regression on distance Matrices (MRM) to test for a relationship between pairwise perceptual distances and pairwise geographic distances. Significance of the test was based on random permutations of distances in the matrices such as the null hypothesis described a random association between perceptual and geographic distances across heliconiine subspecies (**Figs. S21A & S21B** in **Appendix 9**).

Furthermore, to test for a pattern of wing similarity among spatially co-occurring subspecies that goes beyond the expectation from evolutionary relationship across the taxa, hinting for a case of evolutionary convergence as described in Müller's model, we used MRM to test for a relationship between pairwise perceptual distances and pairwise geographic distances accounting for phylogenetic distances as an additional predictor (**Figs. S21C & S21D** in **Appendix 9**). To compute phylogenetic distances, we used pairwise patristic distances on the phylogeny of Heliconiini (*43*) with terminal branches of null length to describe the unknown relative position of subspecies in their associated species.

Because Schoener's D tend to be sensitive to a threshold effect since any distance between two subspecies with no overlap equals one independently from their spatial adjacency or farness, we also ran complementary analyses describing geographic distances as two categories: sympatric *vs.* allopatric using a threshold of 1 – Schoener's D = 0.8 to discriminate between categories (See **Fig. S22** in **SI Appendix 9**).

### 4 Data availability

All MATLAB and R scripts to carry out analyses are available on GitHub (link to be released upon publication). All 2D perceptual maps, 3D animated perceptual spaces, mimicry ring lists and subspecies images used in the online survey are available in online archives in Zenodo (https://doi.org/10.5281/zenodo.10076355). The online survey for the citizen science data collection is temporarily available on http://memometic.cleverapps.io/ with source code accessible on GitHub (link to be released upon publication). Heliconiine distribution maps were retrieved from https://doi.org/10.5281/zenodo.10903661.

### Supporting Information for Appendices

#### Appendix 1: Metadata for images of subspecies

**Table S1: Source for images used to represent the 432 heliconiine subspecies in the online survey.**

| ID | Genus | Species | Subspecies | Source | Author(s) |
|---|---|---|---|---|---|
| 1 | Agraulis | vanillae | galapagensis | BOA | Gerardo Lamas |
| 2 | Agraulis | vanillae | incarnata | Boyer_Collection | Pierre Boyer |
| 3 | Agraulis | vanillae | insularis | Boyer_Collection | Pierre Boyer |
| 4 | Agraulis | vanillae | lucina | Boyer_Collection | Pierre Boyer |
| 5 | Agraulis | vanillae | maculosa | FLMNH | Chris Jiggins |
| 6 | Agraulis | vanillae | nigrior | BOA | Andrew D Warren |
| 7 | Agraulis | vanillae | vanillae | Boyer_Collection | Pierre Boyer |
| 8 | Dione | glycera | | FLMNH | Chris Jiggins |
| 9 | Dione | juno | andicola | BOA | Gerardo Lamas |
| 10 | Dione | juno | huascuma | Boyer_Collection | Pierre Boyer |
| 11 | Dione | juno | juno | FLMNH | Chris Jiggins |
| 12 | Dione | juno | miraculosa | Boyer_Collection | Pierre Boyer |
| 13 | Dione | moneta | butleri | BOA | Gerardo Lamas |
| 14 | Dione | moneta | poeyii | FLMNH | Chris Jiggins |
| 15 | Dryadula | phaetusa | | FLMNH | Chris Jiggins |
| 16 | Dryas | iulia | alcionea | FLMNH | Chris Jiggins |
| 17 | Eueides | aliphera | aliphera | FLMNH | Chris Jiggins |
| 18 | Eueides | aliphera | cyllenella | FLMNH | Chris Jiggins |
| 19 | Eueides | aliphera | gracilis | Michel_Cast_website | Michel Cast |
| 20 | Eueides | emsleyi | emsleyi | Boyer_Collection | Pierre Boyer |
| 21 | Eueides | heliconioides | eanes | FLMNH | Chris Jiggins |
| 22 | Eueides | heliconioides | heliconioides | FLMNH | Chris Jiggins |
| 23 | Dione | moneta | moneta | FLMNH | Augusto H. B. Rosa |
| 24 | Eueides | isabella | dissoluta | FLMNH | Chris Jiggins |
| 25 | Eueides | isabella | ecuadorensis | FLMNH | Chris Jiggins |
| 26 | Eueides | isabella | hippolinus | Boyer_Collection | Pierre Boyer |
| 27 | Eueides | isabella | huebneri | FLMNH | Chris Jiggins |
| 28 | Eueides | isabella | isabella | FLMNH | Chris Jiggins |
| 29 | Eueides | isabella | melphis | Boyer_Collection | Pierre Boyer |
| 30 | Eueides | isabella | nigricornis | BOA | Andrew D Warren |
| 31 | Eueides | lampeto | acacetes | FLMNH | Chris Jiggins |
| 32 | Dryas | iulia | delila | FLMNH | Augusto H. B. Rosa |
| 33 | Dryas | iulia | dominicana | Boyer_Collection | Pierre Boyer |
| 34 | Eueides | lampeto | nigrofulva | FLMNH | Chris Jiggins |
| 35 | Eueides | libitina | libitina | MNHN_Collection | Maël Doré |
| 36 | Eueides | libitina | malleti | Michel_Cast_website | Michel Cast |
| 37 | Eueides | lineata | | FLMNH | Chris Jiggins |

| | | | | | |
|---|---|---|---|---|---|
| **38** | Eueides | lybia | lybia | FLMNH | Chris Jiggins |
| **39** | Eueides | lybia | lybioides | MNHN_Collection | Maël Doré |
| **40** | Eueides | lybia | olympia | FLMNH | Chris Jiggins |
| **41** | Eueides | isabella | dianasa | Boyer_Collection | Pierre Boyer |
| **42** | Eueides | isabella | eva | Boyer_Collection | Pierre Boyer |
| **43** | Eueides | tales | michaeli | FLMNH | Augusto H. B. Rosa |
| **44** | Eueides | pavana | | FLMNH | Chris Jiggins |
| **45** | Heliconius | demeter | subspnov | FLMNH | Keith Willmott |
| **46** | Eueides | procula | edias | FLMNH | Chris Jiggins |
| **47** | Eueides | procula | eurysaces | FLMNH | Chris Jiggins |
| **48** | Dryas | iulia | nudeola | Boyer_Collection | Pierre Boyer |
| **49** | Eueides | procula | procula | FLMNH | Chris Jiggins |
| **50** | Eueides | procula | vulgiformis | Boyer_Collection | Pierre Boyer |
| **51** | Dryas | iulia | fucatus | Boyer_Collection | Pierre Boyer |
| **52** | Eueides | tales | calathus | FLMNH | Chris Jiggins |
| **53** | Dryas | iulia | martinica | MNHN_Collection | Maël Doré |
| **54** | Eueides | tales | franciscus | Boyer_Collection | Pierre Boyer |
| **55** | Eueides | lybia | orinocensis | FLMNH | Augusto H. B. Rosa |
| **56** | Eueides | tales | pseudeanes | Boyer_Collection | Pierre Boyer |
| **57** | Eueides | tales | pythagoras | FLMNH | Chris Jiggins |
| **58** | Eueides | tales | surdus | FLMNH | Chris Jiggins |
| **59** | Eueides | procula | asidia | FLMNH | Augusto H. B. Rosa |
| **60** | Eueides | tales | tales | MNHN_Collection | Maël Doré |
| **61** | Dryas | iulia | moderata | Boyer_Collection | Pierre Boyer |
| **62** | Eueides | vibilia | unifasciatus | FLMNH | Chris Jiggins |
| **63** | Eueides | vibilia | vialis | FLMNH | Chris Jiggins |
| **64** | Eueides | vibilia | vibilia | FLMNH | Chris Jiggins |
| **65** | Eueides | vibilia | vicinalis | Boyer_Collection | Pierre Boyer |
| **66** | Heliconius | antiochus | antiochus | FLMNH | Chris Jiggins |
| **67** | Heliconius | antiochus | aranea | FLMNH | Chris Jiggins |
| **68** | Heliconius | antiochus | araneides | Boyer_Collection | Pierre Boyer |
| **69** | Heliconius | antiochus | salvinii | FLMNH | Chris Jiggins |
| **70** | Heliconius | aoede | aliciae | FLMNH | Chris Jiggins |
| **71** | Heliconius | aoede | aoede | FLMNH | Chris Jiggins |
| **72** | Heliconius | aoede | astydamia | FLMNH | Chris Jiggins |
| **73** | Heliconius | aoede | ayacuchensis | FLMNH | Chris Jiggins |
| **74** | Heliconius | aoede | bartletti | FLMNH | Chris Jiggins |
| **75** | Heliconius | aoede | centurius | FLMNH | Chris Jiggins |
| **76** | Heliconius | aoede | cupidineus | FLMNH | Chris Jiggins |
| **77** | Heliconius | aoede | emmelina | Boyer_Collection | Pierre Boyer |
| **78** | Heliconius | aoede | eurycleia | Boyer_Collection | Pierre Boyer |
| **79** | Heliconius | aoede | faleria | Heliconius.net | NA |
| **80** | Heliconius | aoede | lucretius | FLMNH | Chris Jiggins |
| **81** | Heliconius | aoede | manu | FLMNH | Chris Jiggins |
| **82** | Heliconius | aoede | philipi | FLMNH | Chris Jiggins |

| | | | | | |
|---|---|---|---|---|---|
| **83** | Heliconius | astraea | astraea | FLMNH | Chris Jiggins |
| **84** | Heliconius | astraea | rondonia | FLMNH | Chris Jiggins |
| **85** | Heliconius | atthis | | FLMNH | Chris Jiggins |
| **86** | Heliconius | besckei | | FLMNH | Chris Jiggins |
| **87** | Heliconius | burneyi | ada | Boyer_Collection | Pierre Boyer |
| **88** | Heliconius | burneyi | anjae | FLMNH | Chris Jiggins |
| **89** | Heliconius | burneyi | burneyi | FLMNH | Chris Jiggins |
| **90** | Heliconius | burneyi | catharinae | FLMNH | Chris Jiggins |
| **91** | Heliconius | burneyi | huebneri | FLMNH | Chris Jiggins |
| **92** | Heliconius | burneyi | jamesi | FLMNH | Chris Jiggins |
| **93** | Heliconius | burneyi | lindigii | MNHN_Collection | Maël Doré |
| **94** | Agraulis | vanillae | forbesi | FLMNH | Keith Willmott |
| **95** | Heliconius | burneyi | skinneri | FLMNH | Chris Jiggins |
| **96** | Heliconius | charithonia | bassleri | FLMNH | Chris Jiggins |
| **97** | Heliconius | charithonia | charithonia | FLMNH | Chris Jiggins |
| **98** | Heliconius | charithonia | churchi | FLMNH | Chris Jiggins |
| **99** | Heliconius | charithonia | ramsdeni | FLMNH | Chris Jiggins |
| **100** | Heliconius | charithonia | simulator | FLMNH | Chris Jiggins |
| **101** | Heliconius | charithonia | tuckeri | FLMNH | Chris Jiggins |
| **102** | Heliconius | charithonia | vazquezae | FLMNH | Chris Jiggins |
| **103** | Heliconius | clysonymus | clysonymus | FLMNH | Chris Jiggins |
| **104** | Heliconius | clysonymus | hygiana | FLMNH | Chris Jiggins |
| **105** | Heliconius | clysonymus | montanus | FLMNH | Chris Jiggins |
| **106** | Philaethria | dido | panamensis | Remi_Mauxion | Remi Mauxion |
| **107** | Heliconius | congener | aquilionaris | MNHN_Collection | Maël Doré |
| **108** | Heliconius | congener | congener | FLMNH | Chris Jiggins |
| **109** | Philaethria | dido | dido | FLMNH | Chris Jiggins |
| **110** | Heliconius | cydno | alithea | FLMNH | Chris Jiggins |
| **111** | Heliconius | cydno | barinasensis | FLMNH | Chris Jiggins |
| **112** | Heliconius | cydno | chioneus | FLMNH | Chris Jiggins |
| **113** | Heliconius | cydno | cordula | FLMNH | Chris Jiggins |
| **114** | Heliconius | cydno | cydnides | FLMNH | Chris Jiggins |
| **115** | Heliconius | cydno | cydno | FLMNH | Chris Jiggins |
| **116** | Heliconius | cydno | gadouae | FLMNH | Chris Jiggins |
| **117** | Heliconius | cydno | galanthus | FLMNH | Chris Jiggins |
| **118** | Heliconius | cydno | hermogenes | FLMNH | Chris Jiggins |
| **119** | Heliconius | cydno | lisethae | FLMNH | Chris Jiggins |
| **120** | Heliconius | cydno | wanningeri | FLMNH | Chris Jiggins |
| **121** | Heliconius | cydno | weymeri | FLMNH | Chris Jiggins |
| **122** | Heliconius | cydno | zelinde | FLMNH | Chris Jiggins |
| **123** | Philaethria | dido | chocoensis | Boyer_Collection | Pierre Boyer |
| **124** | Heliconius | demeter | bouqueti | FLMNH | Chris Jiggins |
| **125** | Heliconius | demeter | demeter | FLMNH | Chris Jiggins |
| **126** | Heliconius | demeter | joroni | Michel_Cast_website | Michel Cast |
| **127** | Heliconius | demeter | karinae | FLMNH | Chris Jiggins |

| | | | | | |
|---|---|---|---|---|---|
| **128** | Heliconius | demeter | neildi | FLMNH | Chris Jiggins |
| **129** | Heliconius | demeter | terrasanta | FLMNH | Chris Jiggins |
| **130** | Heliconius | demeter | titan | FLMNH | Chris Jiggins |
| **131** | Heliconius | demeter | turneri | FLMNH | Chris Jiggins |
| **132** | Philaethria | diatonica | | FLMNH | Chris Jiggins |
| **133** | Heliconius | doris | dives | Boyer_Collection | Pierre Boyer |
| **134** | Heliconius | doris | doris | FLMNH | Chris Jiggins |
| **135** | Heliconius | doris | obscurus | FLMNH | Chris Jiggins |
| **136** | Heliconius | doris | viridis | FLMNH | Chris Jiggins |
| **137** | Heliconius | egeria | egeria | FLMNH | Chris Jiggins |
| **138** | Heliconius | egeria | egerides | Boyer_Collection | Pierre Boyer |
| **139** | Heliconius | egeria | homogena | FLMNH | Chris Jiggins |
| **140** | Heliconius | egeria | hyas | FLMNH | Chris Jiggins |
| **141** | Heliconius | egeria | keithbrowni | FLMNH | Chris Jiggins |
| **142** | Heliconius | eleuchia | eleuchia | FLMNH | Chris Jiggins |
| **143** | Heliconius | eleuchia | eleusinus | FLMNH | Chris Jiggins |
| **144** | Heliconius | eleuchia | primularis | FLMNH | Chris Jiggins |
| **145** | Heliconius | elevatus | bari | Michel_Cast_website | Michel Cast |
| **146** | Heliconius | elevatus | elevatus | FLMNH | Chris Jiggins |
| **147** | Heliconius | elevatus | lapis | FLMNH | Chris Jiggins |
| **148** | Heliconius | elevatus | perchlora | FLMNH | Chris Jiggins |
| **149** | Heliconius | elevatus | pseudocupidineus | FLMNH | Chris Jiggins |
| **150** | Heliconius | elevatus | roraima | FLMNH | Chris Jiggins |
| **151** | Heliconius | elevatus | schmassmanni | FLMNH | Chris Jiggins |
| **152** | Heliconius | elevatus | taracuanus | FLMNH | Chris Jiggins |
| **153** | Heliconius | elevatus | tumatumari | FLMNH | Keith Willmott |
| **154** | Heliconius | elevatus | zoelleri | FLMNH | Chris Jiggins |
| **155** | Heliconius | erato | adana | FLMNH | Chris Jiggins |
| **156** | Heliconius | erato | amalfreda | FLMNH | Chris Jiggins |
| **157** | Heliconius | erato | amazona | FLMNH | Chris Jiggins |
| **158** | Heliconius | erato | amphitrite | FLMNH | Chris Jiggins |
| **159** | Heliconius | erato | chestertonii | FLMNH | Chris Jiggins |
| **160** | Heliconius | erato | colombina | Boyer_Collection | Pierre Boyer |
| **161** | Philaethria | constantinoi | | FLMNH | Chris Jiggins |
| **162** | Heliconius | erato | cyrbia | FLMNH | Chris Jiggins |
| **163** | Heliconius | erato | demophoon | FLMNH | Chris Jiggins |
| **164** | Heliconius | erato | dignus | FLMNH | Chris Jiggins |
| **165** | Heliconius | erato | emma | FLMNH | Chris Jiggins |
| **166** | Heliconius | erato | erato | FLMNH | Chris Jiggins |
| **167** | Heliconius | erato | estrella | Boyer_Collection | Pierre Boyer |
| **168** | Heliconius | erato | etylus | FLMNH | Chris Jiggins |
| **169** | Heliconius | erato | favorinus | FLMNH | Chris Jiggins |
| **170** | Heliconius | erato | guarica | FLMNH | Chris Jiggins |
| **171** | Heliconius | erato | hydara | FLMNH | Chris Jiggins |
| **172** | Heliconius | erato | lativitta | FLMNH | Chris Jiggins |

| | | | | | |
|---|---|---|---|---|---|
| **173** | Heliconius | erato | lichyi | FLMNH | Chris Jiggins |
| **174** | Heliconius | erato | luscombei | FLMNH | Chris Jiggins |
| **175** | Heliconius | erato | magnifica | FLMNH | Chris Jiggins |
| **176** | Heliconius | erato | microclea | FLMNH | Chris Jiggins |
| **177** | Heliconius | erato | notabilis | FLMNH | Chris Jiggins |
| **178** | Heliconius | erato | phyllis | FLMNH | Chris Jiggins |
| **179** | Heliconius | erato | reductimacula | FLMNH | Chris Jiggins |
| **180** | Heliconius | erato | tobagoensis | FLMNH | Chris Jiggins |
| **181** | Heliconius | erato | venus | FLMNH | Chris Jiggins |
| **182** | Heliconius | erato | venustus | FLMNH | Chris Jiggins |
| **183** | Philaethria | andrei | andrei | BOA | Christian Brevignon |
| **184** | Heliconius | eratosignis | tambopata | FLMNH | Keith Willmott |
| **185** | Heliconius | eratosignis | ucayalensis | FLMNH | Chris Jiggins |
| **186** | Heliconius | eratosignis | ulysses | FLMNH | Chris Jiggins |
| **187** | Heliconius | ethilla | adela | FLMNH | Chris Jiggins |
| **188** | Heliconius | ethilla | aerotome | FLMNH | Chris Jiggins |
| **189** | Heliconius | ethilla | cephallenia | Boyer_Collection | Pierre Boyer |
| **190** | Heliconius | ethilla | chapadensis | FLMNH | Chris Jiggins |
| **191** | Heliconius | ethilla | claudia | FLMNH | Chris Jiggins |
| **192** | Heliconius | ethilla | ethilla | FLMNH | Chris Jiggins |
| **193** | Heliconius | ethilla | eucoma | FLMNH | Chris Jiggins |
| **194** | Heliconius | ethilla | flavofasciatus | FLMNH | Augusto H. B. Rosa |
| **195** | Heliconius | ethilla | flavomaculatus | FLMNH | Chris Jiggins |
| **196** | Heliconius | ethilla | hyalina | FLMNH | Keith Willmott |
| **197** | Heliconius | ethilla | jaruensis | FLMNH | Chris Jiggins |
| **198** | Heliconius | ethilla | latona | FLMNH | Chris Jiggins |
| **199** | Heliconius | ethilla | mentor | Boyer_Collection | Pierre Boyer |
| **200** | Heliconius | ethilla | metalilis | FLMNH | Chris Jiggins |
| **201** | Heliconius | ethilla | michaelianius | FLMNH | Chris Jiggins |
| **202** | Heliconius | ethilla | narcaea | FLMNH | Chris Jiggins |
| **203** | Heliconius | xanthocles | zamora | FLMNH | Chris Jiggins |
| **204** | Heliconius | ethilla | numismaticus | FLMNH | Chris Jiggins |
| **205** | Heliconius | ethilla | penthesilea | FLMNH | Chris Jiggins |
| **206** | Heliconius | ethilla | polychrous | FLMNH | Chris Jiggins |
| **207** | Heliconius | ethilla | semiflavidus | FLMNH | Chris Jiggins |
| **208** | Heliconius | ethilla | thielei | FLMNH | Chris Jiggins |
| **209** | Heliconius | ethilla | tyndarus | FLMNH | Chris Jiggins |
| **210** | Heliconius | ethilla | yuruani | FLMNH | Chris Jiggins |
| **211** | Heliconius | godmani | | FLMNH | Chris Jiggins |
| **212** | Heliconius | hecale | anderida | FLMNH | Chris Jiggins |
| **213** | Heliconius | hecale | annetta | FLMNH | Chris Jiggins |
| **214** | Heliconius | hecale | australis | FLMNH | Chris Jiggins |
| **215** | Heliconius | hecale | barcanti | MNHN_Collection | Maël Doré |
| **216** | Heliconius | hecale | clearei | FLMNH | Chris Jiggins |
| **217** | Heliconius | hecale | ennius | FLMNH | Chris Jiggins |

| | | | | | |
|---|---|---|---|---|---|
| **218** | Heliconius | hecale | felix | FLMNH | Chris Jiggins |
| **219** | Heliconius | hecale | fornarina | FLMNH | Chris Jiggins |
| **220** | Heliconius | hecale | hecale | FLMNH | Chris Jiggins |
| **221** | Heliconius | hecale | holcophorus | FLMNH | Chris Jiggins |
| **222** | Heliconius | hecale | humboldti | FLMNH | Chris Jiggins |
| **223** | Heliconius | hecale | ithaca | FLMNH | Chris Jiggins |
| **224** | Heliconius | hecale | latus | FLMNH | Chris Jiggins |
| **225** | Heliconius | hecale | melicerta | FLMNH | Chris Jiggins |
| **226** | Heliconius | hecale | metellus | FLMNH | Chris Jiggins |
| **227** | Heliconius | hecale | nigrofasciatus | FLMNH | Chris Jiggins |
| **228** | Heliconius | hecale | novatus | FLMNH | Chris Jiggins |
| **229** | Heliconius | hecale | paraensis | FLMNH | Keith Willmott |
| **230** | Heliconius | hecale | paulus | FLMNH | Chris Jiggins |
| **231** | Heliconius | hecale | quitalena | FLMNH | Chris Jiggins |
| **232** | Heliconius | hecale | rosalesi | FLMNH | Keith Willmott |
| **233** | Heliconius | hecale | shanki | FLMNH | Chris Jiggins |
| **234** | Heliconius | hecale | sisyphus | FLMNH | Chris Jiggins |
| **235** | Heliconius | hecale | sulphureus | FLMNH | Chris Jiggins |
| **236** | Heliconius | hecale | vetustus | FLMNH | Chris Jiggins |
| **237** | Heliconius | hecale | zuleika | FLMNH | Chris Jiggins |
| **238** | Heliconius | hecalesia | eximius | Boyer_Collection | Pierre Boyer |
| **239** | Heliconius | hecalesia | formosus | FLMNH | Chris Jiggins |
| **240** | Heliconius | hecalesia | gynaesia | FLMNH | Chris Jiggins |
| **241** | Heliconius | hecalesia | hecalesia | FLMNH | Chris Jiggins |
| **242** | Heliconius | hecalesia | longarena | FLMNH | Chris Jiggins |
| **243** | Heliconius | hecalesia | octavia | FLMNH | Chris Jiggins |
| **244** | Heliconius | hecalesia | romeroi | FLMNH | Chris Jiggins |
| **245** | Heliconius | hecuba | cassandra | Boyer_Collection | Pierre Boyer |
| **246** | Heliconius | hecuba | choarina | FLMNH | Chris Jiggins |
| **247** | Heliconius | hecuba | creusa | FLMNH | Chris Jiggins |
| **248** | Heliconius | xanthocles | xanthocles | FLMNH | Chris Jiggins |
| **249** | Heliconius | hecuba | flava | FLMNH | Chris Jiggins |
| **250** | Heliconius | hecuba | hecuba | FLMNH | Chris Jiggins |
| **251** | Heliconius | hecuba | tolima | FLMNH | Chris Jiggins |
| **252** | Heliconius | hermathena | curua | FLMNH | Augusto H. B. Rosa |
| **253** | Heliconius | hermathena | duckei | FLMNH | Augusto H. B. Rosa |
| **254** | Heliconius | hermathena | hermathena | FLMNH | Chris Jiggins |
| **255** | Heliconius | hermathena | renatae | FLMNH | Chris Jiggins |
| **256** | Heliconius | hermathena | sabinae | FLMNH | Chris Jiggins |
| **257** | Heliconius | hermathena | sheppardi | FLMNH | Augusto H. B. Rosa |
| **258** | Heliconius | hermathena | vereatta | FLMNH | Augusto H. B. Rosa |
| **259** | Heliconius | heurippa | | FLMNH | Chris Jiggins |
| **260** | Heliconius | hewitsoni | | FLMNH | Chris Jiggins |
| **261** | Heliconius | hierax | hierax | FLMNH | Chris Jiggins |
| **262** | Heliconius | xanthocles | vala | FLMNH | Chris Jiggins |

| | | | | | |
|---|---|---|---|---|---|
| **263** | Heliconius | himera | | FLMNH | Chris Jiggins |
| **264** | Heliconius | hortense | | FLMNH | Chris Jiggins |
| **265** | Heliconius | ismenius | boulleti | FLMNH | Chris Jiggins |
| **266** | Heliconius | ismenius | clarescens | FLMNH | Chris Jiggins |
| **267** | Heliconius | ismenius | fasciatus | MNHN_Collection | Maël Doré |
| **268** | Heliconius | ismenius | ismenius | FLMNH | Chris Jiggins |
| **269** | Heliconius | ismenius | metaphorus | FLMNH | Chris Jiggins |
| **270** | Heliconius | ismenius | occidentalis | FLMNH | Chris Jiggins |
| **271** | Heliconius | ismenius | telchinia | FLMNH | Chris Jiggins |
| **272** | Heliconius | ismenius | tilletti | FLMNH | Keith Willmott |
| **273** | Heliconius | lalitae | | FLMNH | Chris Jiggins |
| **274** | Heliconius | leucadia | leucadia | FLMNH | Chris Jiggins |
| **275** | Heliconius | leucadia | pseudorhea | FLMNH | Chris Jiggins |
| **276** | Heliconius | xanthocles | similatus | FLMNH | Chris Jiggins |
| **277** | Heliconius | luciana | watunna | FLMNH | Chris Jiggins |
| **278** | Heliconius | melpomene | aglaope | FLMNH | Chris Jiggins |
| **279** | Heliconius | melpomene | amandus | FLMNH | Chris Jiggins |
| **280** | Heliconius | melpomene | amaryllis | FLMNH | Chris Jiggins |
| **281** | Heliconius | melpomene | anduzei | FLMNH | Chris Jiggins |
| **282** | Heliconius | melpomene | bellula | FLMNH | Chris Jiggins |
| **283** | Heliconius | melpomene | burchelli | FLMNH | Chris Jiggins |
| **284** | Heliconius | melpomene | cythera | FLMNH | Chris Jiggins |
| **285** | Heliconius | melpomene | ecuadorensis | FLMNH | Chris Jiggins |
| **286** | Heliconius | melpomene | euryades | FLMNH | Keith Willmott |
| **287** | Heliconius | melpomene | flagrans | FLMNH | Chris Jiggins |
| **288** | Heliconius | melpomene | intersectus | FLMNH | Keith Willmott |
| **289** | Heliconius | melpomene | madeira | Boyer_Collection | Pierre Boyer |
| **290** | Heliconius | melpomene | malleti | FLMNH | Chris Jiggins |
| **291** | Heliconius | melpomene | martinae | Michel_Cast_website | Michel Cast |
| **292** | Heliconius | melpomene | melpomene | FLMNH | Chris Jiggins |
| **293** | Heliconius | melpomene | meriana | Boyer_Collection | Pierre Boyer |
| **294** | Heliconius | melpomene | michellae | FLMNH | Chris Jiggins |
| **295** | Heliconius | melpomene | nanna | FLMNH | Chris Jiggins |
| **296** | Heliconius | melpomene | penelope | FLMNH | Chris Jiggins |
| **297** | Heliconius | melpomene | plesseni | FLMNH | Chris Jiggins |
| **298** | Heliconius | melpomene | pyrforus | FLMNH | Chris Jiggins |
| **299** | Heliconius | melpomene | rosina | FLMNH | Chris Jiggins |
| **300** | Heliconius | melpomene | schunkei | FLMNH | Chris Jiggins |
| **301** | Heliconius | melpomene | tessa | FLMNH | Chris Jiggins |
| **302** | Heliconius | melpomene | thelxiope | FLMNH | Chris Jiggins |
| **303** | Heliconius | melpomene | thelxiopeia | Boyer_Collection | Pierre Boyer |
| **304** | Heliconius | melpomene | vicina | FLMNH | Chris Jiggins |
| **305** | Heliconius | melpomene | vulcanus | FLMNH | Chris Jiggins |
| **306** | Heliconius | melpomene | xenoclea | FLMNH | Chris Jiggins |
| **307** | Heliconius | metharme | makiritare | FLMNH | Chris Jiggins |

| | | | | | |
|---|---|---|---|---|---|
| **308** | Heliconius | metharme | metharme | FLMNH | Chris Jiggins |
| **309** | Heliconius | metharme | perseis | FLMNH | Chris Jiggins |
| **310** | Philaethria | ostara | ostara | BOA | Keith Willmott |
| **311** | Heliconius | nattereri | | FLMNH | Chris Jiggins |
| **312** | Heliconius | numata | arcuella | FLMNH | Chris Jiggins |
| **313** | Heliconius | numata | aristiona | FLMNH | Chris Jiggins |
| **314** | Heliconius | numata | aulicus | FLMNH | Chris Jiggins |
| **315** | Heliconius | numata | aurora | FLMNH | Chris Jiggins |
| **316** | Heliconius | numata | bicoloratus | FLMNH | Chris Jiggins |
| **317** | Heliconius | numata | elegans | Boyer_Collection | Pierre Boyer |
| **318** | Heliconius | numata | ethra | FLMNH | Chris Jiggins |
| **319** | Heliconius | numata | euphone | FLMNH | Chris Jiggins |
| **320** | Heliconius | numata | euphrasius | Boyer_Collection | Pierre Boyer |
| **321** | Heliconius | numata | geminatus | Boyer_Collection | Pierre Boyer |
| **322** | Heliconius | numata | holzingeri | FLMNH | Chris Jiggins |
| **323** | Heliconius | numata | ignotus | FLMNH | Chris Jiggins |
| **324** | Heliconius | numata | illustris | FLMNH | Chris Jiggins |
| **325** | Heliconius | numata | isabellinus | NHMUK | B. Huertas & R. Crowther |
| **326** | Heliconius | numata | jiparanaensis | FLMNH | Chris Jiggins |
| **327** | Heliconius | numata | laura | Boyer_Collection | Pierre Boyer |
| **328** | Heliconius | numata | lenaeus | FLMNH | Chris Jiggins |
| **329** | Heliconius | numata | lyrcaeus | FLMNH | Chris Jiggins |
| **330** | Heliconius | numata | mavors | FLMNH | Chris Jiggins |
| **331** | Heliconius | numata | messene | FLMNH | Chris Jiggins |
| **332** | Heliconius | numata | mirus | FLMNH | Chris Jiggins |
| **333** | Heliconius | numata | nubifer | FLMNH | Chris Jiggins |
| **334** | Heliconius | numata | numata | FLMNH | Chris Jiggins |
| **335** | Heliconius | numata | peeblesi | Boyer_Collection | Pierre Boyer |
| **336** | Heliconius | numata | pratti | FLMNH | Chris Jiggins |
| **337** | Heliconius | numata | robigus | FLMNH | Chris Jiggins |
| **338** | Heliconius | numata | silvana | FLMNH | Chris Jiggins |
| **339** | Heliconius | numata | sourensis | FLMNH | Chris Jiggins |
| **340** | Heliconius | numata | superioris | FLMNH | Chris Jiggins |
| **341** | Heliconius | numata | talboti | FLMNH | Chris Jiggins |
| **342** | Heliconius | numata | tarapotensis | Boyer_Collection | Pierre Boyer |
| **343** | Heliconius | numata | timaeus | Boyer_Collection | Pierre Boyer |
| **344** | Heliconius | numata | zobrysi | FLMNH | Chris Jiggins |
| **345** | Heliconius | pachinus | | FLMNH | Chris Jiggins |
| **346** | Podotricha | telesiphe | telesiphe | FLMNH | Chris Jiggins |
| **347** | Heliconius | pardalinus | butleri | Boyer_Collection | Pierre Boyer |
| **348** | Heliconius | pardalinus | dilatus | Boyer_Collection | Pierre Boyer |
| **349** | Podotricha | telesiphe | tithraustes | BOA | Gerardo Lamas |
| **350** | Heliconius | pardalinus | lucescens | MNHN_Collection | Maël Doré |
| **351** | Heliconius | pardalinus | maeon | FLMNH | Chris Jiggins |
| **352** | Heliconius | pardalinus | orteguaza | FLMNH | Keith Willmott |

| | | | | | |
|---|---|---|---|---|---|
| **353** | Heliconius | pardalinus | pardalinus | FLMNH | Chris Jiggins |
| **354** | Heliconius | pardalinus | radiosus | FLMNH | Keith Willmott |
| **355** | Heliconius | pardalinus | sergestus | FLMNH | Chris Jiggins |
| **356** | Heliconius | pardalinus | tithoreides | FLMNH | Chris Jiggins |
| **357** | Heliconius | peruvianus | | FLMNH | Chris Jiggins |
| **358** | Heliconius | ricini | insulanus | FLMNH | Chris Jiggins |
| **359** | Heliconius | ricini | ricini | FLMNH | Chris Jiggins |
| **360** | Heliconius | sapho | candidus | FLMNH | Chris Jiggins |
| **361** | Heliconius | sapho | chocoensis | FLMNH | Chris Jiggins |
| **362** | Heliconius | sapho | leuce | FLMNH | Chris Jiggins |
| **363** | Heliconius | sapho | sapho | FLMNH | Chris Jiggins |
| **364** | Heliconius | sara | apseudes | FLMNH | Chris Jiggins |
| **365** | Heliconius | sara | brevimaculata | Boyer_Collection | Pierre Boyer |
| **366** | Heliconius | sara | elektra | FLMNH | Chris Jiggins |
| **367** | Heliconius | sara | fulgidus | FLMNH | Chris Jiggins |
| **368** | Heliconius | sara | magdalena | FLMNH | Chris Jiggins |
| **369** | Heliconius | sara | sara | FLMNH | Chris Jiggins |
| **370** | Heliconius | sara | sprucei | FLMNH | Chris Jiggins |
| **371** | Heliconius | sara | theudela | FLMNH | Chris Jiggins |
| **372** | Heliconius | sara | veraepacis | FLMNH | Chris Jiggins |
| **373** | Heliconius | sara | williami | FLMNH | Chris Jiggins |
| **374** | Heliconius | telesiphe | cretacea | FLMNH | Chris Jiggins |
| **375** | Heliconius | telesiphe | sotericus | FLMNH | Chris Jiggins |
| **376** | Heliconius | telesiphe | telesiphe | FLMNH | Chris Jiggins |
| **377** | Philaethria | wernickei | | Boyer_Collection | Pierre Boyer |
| **378** | Podotricha | judith | straminea | Boyer_Collection | Pierre Boyer |
| **379** | Heliconius | timareta | thelxinoe | Boyer_Collection | Pierre Boyer |
| **380** | Heliconius | timareta | timareta | FLMNH | Chris Jiggins |
| **381** | Podotricha | judith | caucana | Boyer_Collection | Pierre Boyer |
| **382** | Podotricha | judith | judith | Boyer_Collection | Pierre Boyer |
| **383** | Podotricha | judith | mellosa | BOA | Gerardo Lamas |
| **384** | Heliconius | wallacei | colon | Boyer_Collection | Pierre Boyer |
| **385** | Heliconius | wallacei | flavescens | FLMNH | Chris Jiggins |
| **386** | Heliconius | wallacei | kayei | FLMNH | Chris Jiggins |
| **387** | Heliconius | wallacei | mimulinus | Boyer_Collection | Pierre Boyer |
| **388** | Heliconius | wallacei | wallacei | FLMNH | Chris Jiggins |
| **389** | Heliconius | xanthocles | buechei | FLMNH | Chris Jiggins |
| **390** | Heliconius | xanthocles | cleoxanthe | FLMNH | Chris Jiggins |
| **391** | Heliconius | xanthocles | donatia | Boyer_Collection | Pierre Boyer |
| **392** | Heliconius | xanthocles | explicata | FLMNH | Chris Jiggins |
| **393** | Heliconius | xanthocles | hippocrene | FLMNH | Chris Jiggins |
| **394** | Heliconius | xanthocles | melete | FLMNH | Chris Jiggins |
| **395** | Heliconius | xanthocles | melior | FLMNH | Chris Jiggins |
| **396** | Heliconius | xanthocles | melittus | FLMNH | Chris Jiggins |
| **397** | Heliconius | xanthocles | napoensis | FLMNH | Chris Jiggins |

| | | | | | |
|---|---|---|---|---|---|
| **398** | Heliconius | xanthocles | paraplesius | FLMNH | Chris Jiggins |
| **399** | Heliconius | xanthocles | quindecim | FLMNH | Chris Jiggins |
| **400** | Heliconius | xanthocles | rindgei | FLMNH | Chris Jiggins |
| **401** | Eueides | isabella | cleobaea | MNHN_Collection | Maël Doré |
| **402** | Heliconius | hecuba | crispus | MNHN_Collection | Maël Doré |
| **403** | Heliconius | erato | petiverana | MNHN_Collection | Maël Doré |
| **404** | Dione | juno | suffumata | NHMUK | B. Huertas & R. Crowther |
| **405** | Dryas | iulia | carteri | NHMUK | B. Huertas & R. Crowther |
| **406** | Dryas | iulia | framptoni | NHMUK | B. Huertas & R. Crowther |
| **407** | Dryas | iulia | warneri | NHMUK | B. Huertas & R. Crowther |
| **408** | Dryas | iulia | lucia | NHMUK | B. Huertas & R. Crowther |
| **409** | Eueides | lampeto | apicalis | NHMUK | B. Huertas & R. Crowther |
| **410** | Eueides | lampeto | lampeto | NHMUK | B. Huertas & R. Crowther |
| **411** | Eueides | procula | kuenowii | NHMUK | B. Huertas & R. Crowther |
| **412** | Eueides | tales | barcellinus | NHMUK | B. Huertas & R. Crowther |
| **413** | Eueides | tales | cognata | NHMUK | B. Huertas & R. Crowther |
| **414** | Eueides | tales | tabernula | NHMUK | B. Huertas & R. Crowther |
| **415** | Eueides | tales | xenophanes | NHMUK | B. Huertas & R. Crowther |
| **416** | Heliconius | luciana | luciana | NHMUK | B. Huertas & R. Crowther |
| **417** | Heliconius | pardalinus | julia | NHMUK | B. Huertas & R. Crowther |
| **418** | Heliconius | wallacei | araguaia | NHMUK | B. Huertas & R. Crowther |
| **419** | Eueides | emsleyi | esmeraldensis | Michel_Cast_website | Yves Lever |
| **420** | Eueides | heliconioides | koenigi | Michel_Cast_website | Yves Lever |
| **421** | Eueides | lampeto | brownsbergensis | 10.18473/lepi.v64i3.a7 | H. Gernaat |
| **422** | Eueides | procula | browni | Michel_Cast_website | Yves Lever |
| **423** | Eueides | vibilia | louisi | Michel_Cast_website | Yves Lever |
| **424** | Heliconius | aoede | auca | Michel_Cast_website | Michel Cast |
| **425** | Heliconius | burneyi | boliviensis | Michel_Cast_website | Michel Cast |
| **426** | Heliconius | burneyi | koenigi | Michel_Cast_website | Michel Cast |
| **427** | Heliconius | burneyi | mirtarosa | Michel_Cast_website | Michel Cast |
| **428** | Heliconius | hecuba | lamasi | Michel_Cast_website | Pierre Boyer |
| **429** | Heliconius | metis | | ISSN 0723-9912 | G. Moreira & C. Mielke |
| **430** | Heliconius | timareta | florencia | 10.1186/1471-2148-8-324 | Chris Jiggins |
| **431** | Heliconius | timareta | linaresi | IIRB | Chris Jiggins |
| **432** | Heliconius | timareta | tristero | Michel_Cast_website | D. Lacomme |

#### Appendix 2: Online survey of wing pattern perception from similarity triplets

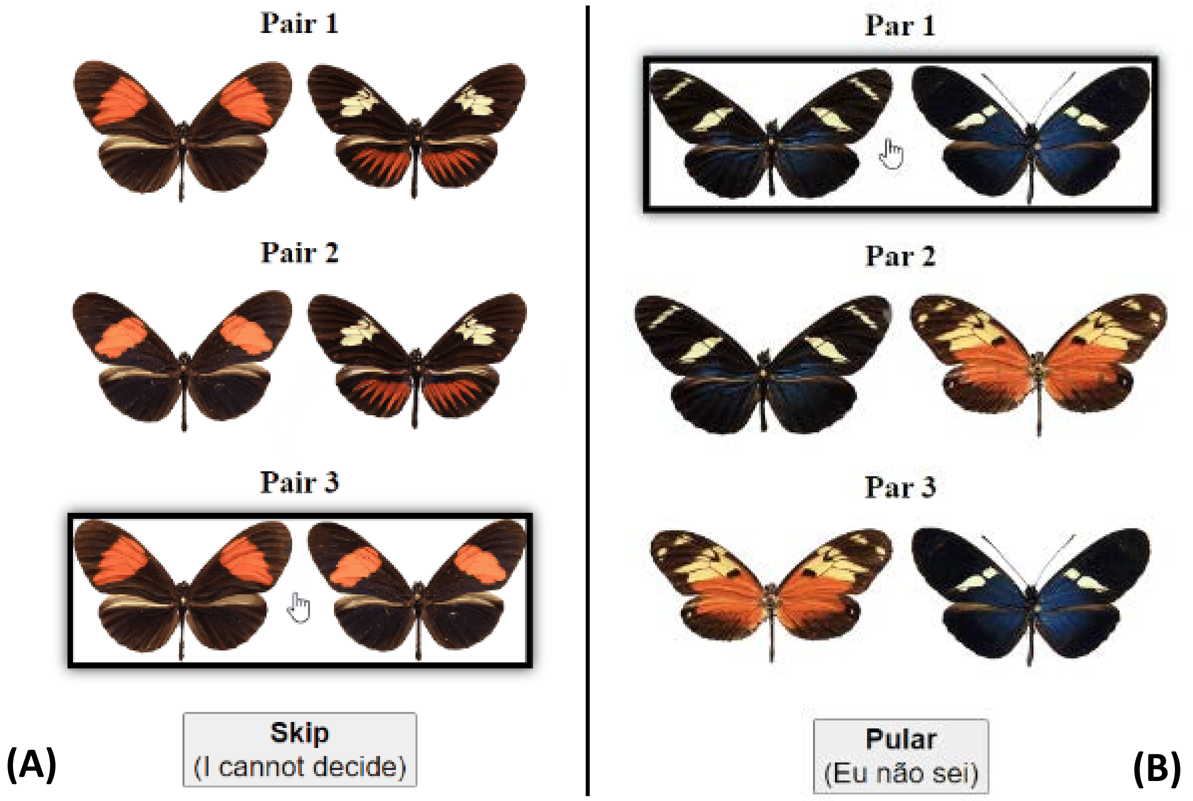


**Figure S1: Two examples of interface for triplet selection in the online citizen science survey.** The player is invited to select the pair of butterflies that displays the more similar color patterns according to its own perception. A skip button is available in case the player is undecided about a particular triplet. The website was made available in multiple languages. **(A)** English interface. **(B)** Brazilian Portuguese interface.

### Appendix 3: Control trials of similarity triplets for the online survey

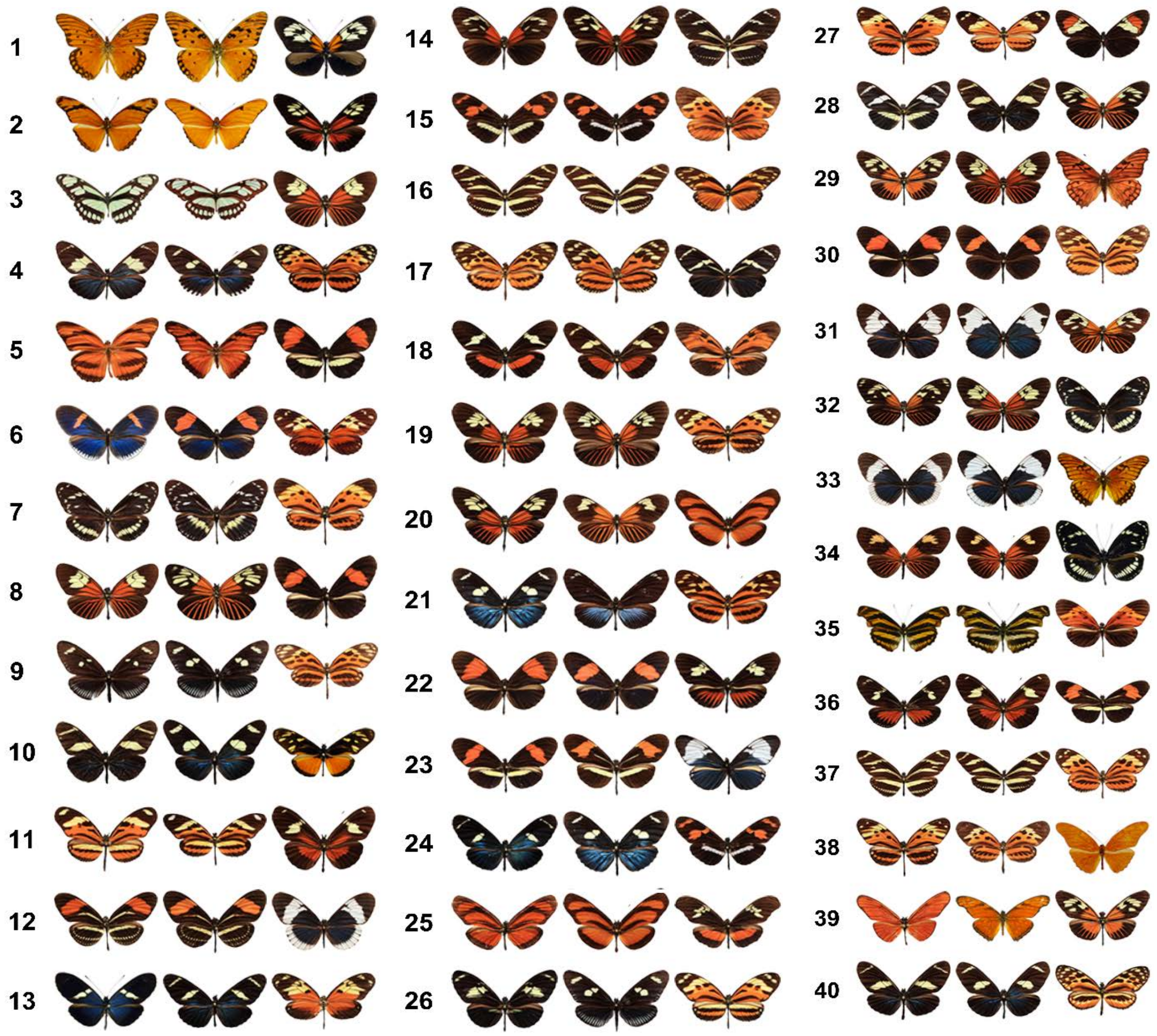


**Figure S2: Triplets of images used as control trials in the online survey.** For each game session, two random control trials were presented. The expected chosen pair for maximum similarity are the first two columns of each triplet. Images in a control trial are displayed at random to avoid positional bias. Game sessions that failed at least one controlled trial were removed from the triplet dataset.

#### Appendix 4: Online survey: demographics and game analytics

##### Website logs:

During the course of the online survey, we recorded 1,422 game sessions from 1,242 distinct players in 474 locations across 53 countries in 5 continents (**Fig. S3**), including 337 locations in 22 countries within Europe (**Fig. S4**).

Worldwide connections

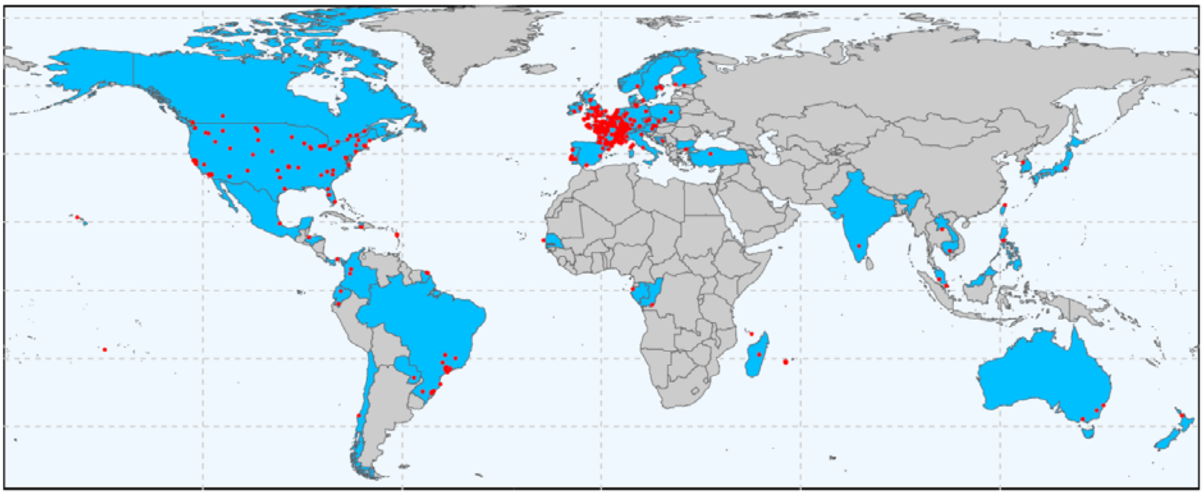

**Figure S3: Location of the 1,422 connections recorded across 474 cities in 53 countries worldwide.**

Europe connections

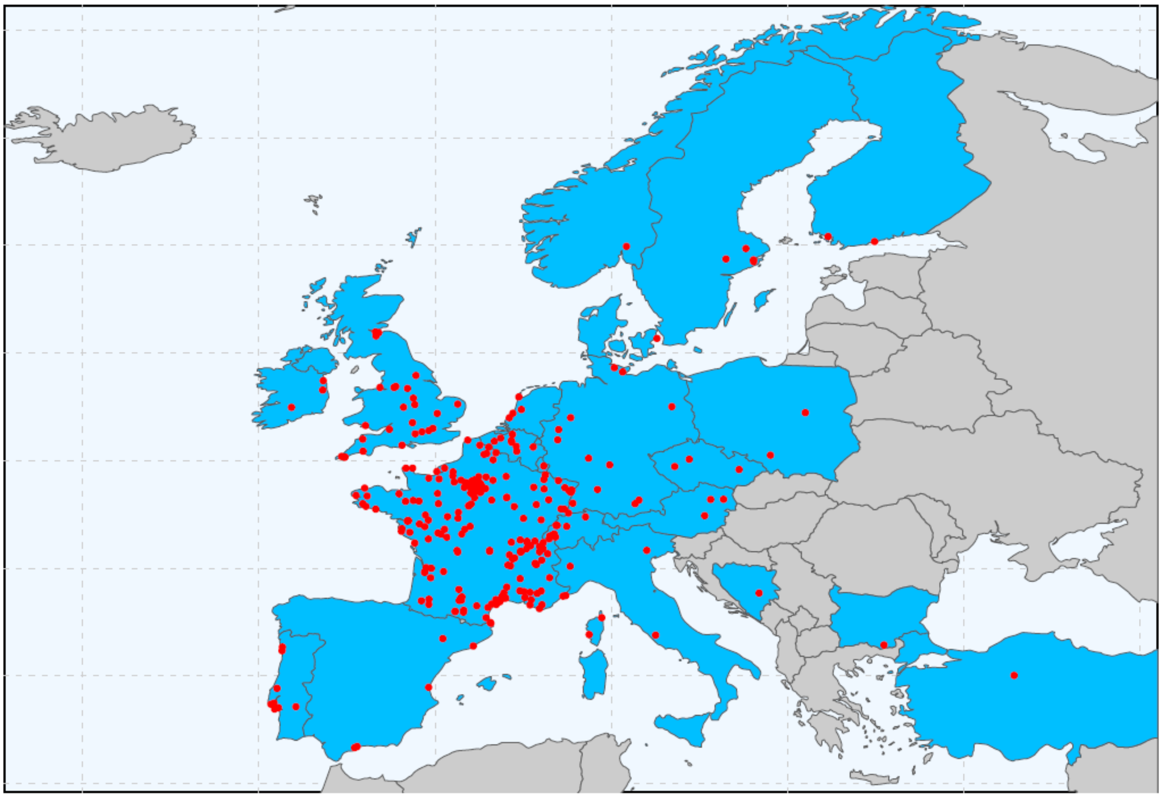

**Figure S4: Location of the 1,010 connections recorded across 337 cities in 22 countries within Europe.**

Data collection was open for six weeks from November 09th to December 18th 2022. We recorded two massive peaks of connections associated with the diffusion of the project to important French and international mailing lists of the community of evolutionary biologists (**Fig. S5**). Clearly, the effect of online publication did not last in time with a poor repeatability of play.

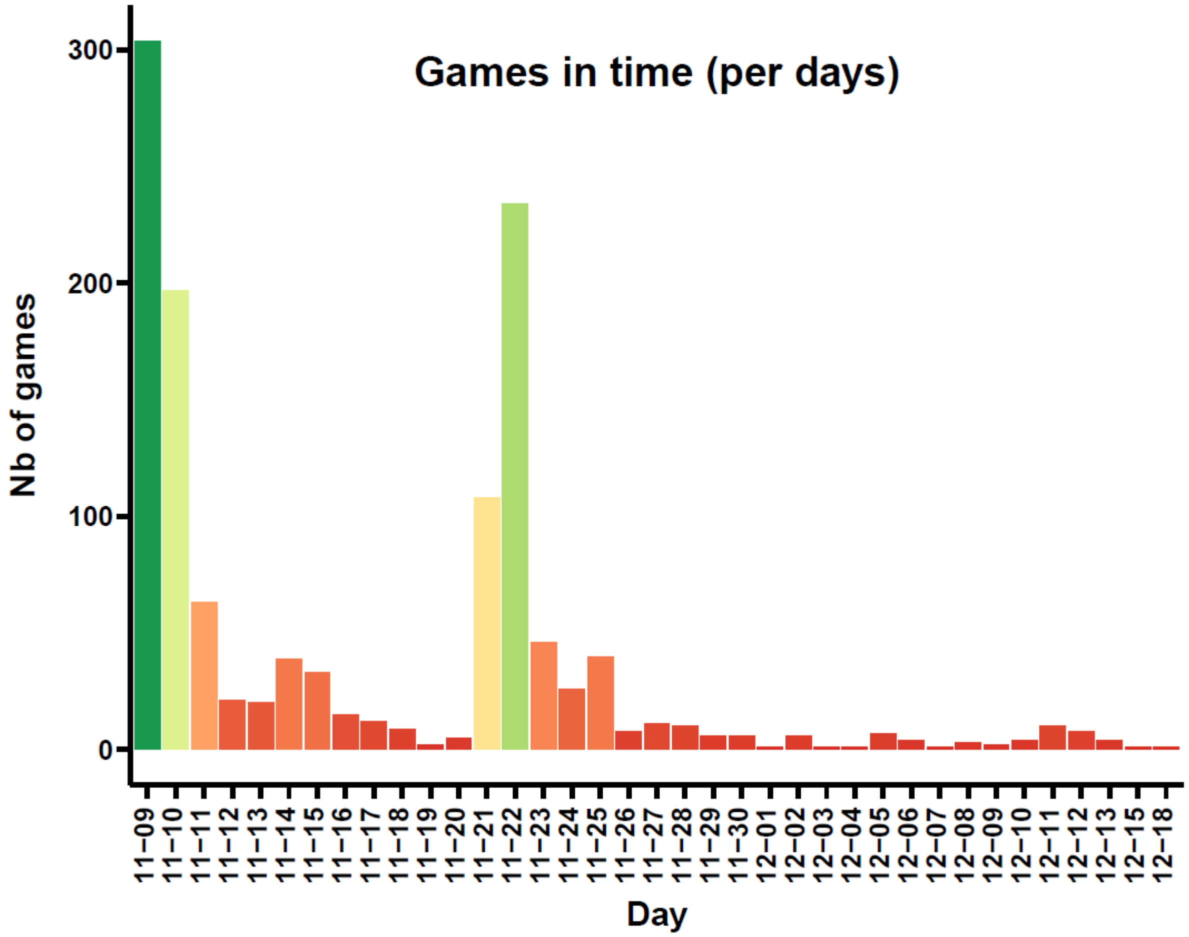


**Figure S5: Frequencies of connections to the online survey (http://memometic.cleverapps.io/).** Records show two massive peaks associated with the timing of publications advertising the project on important French and international mailing lists for evolutionary biologists.

**Playstyle:**

The repeatability of play was fairly poor with a vast majority of players playing only once despite incentive to play multiple times displayed in the online survey. The median number of games was 1.1 (**Fig. S6A**). This low fidelity is likely due to the fact we decided not to provide any direct feedback such as a score at the end of a game session because we did not want to influence the play style of people by guiding them towards more average choices rewarded with an increasing score measuring the degree of closeness to the norm. We recorded time spent per triplet showing most players took around 5 to 10 seconds to examine a triplet (**Fig. S6B**). Most people rarely use the 'Skip option' with 4.4 skips per game on average (**Fig. S6C**), with an important proportion of games without any skip (643 games = 46.1%). It seems likely players perceived the Skip option as a negative action, while it can actually help to eliminate noise in the dataset. A next version of the survey interface should emphasize the benefits from using skips as it is shown to decrease the degree of abnormal triplets recorded (see **Fig. S9A**: normality-score increases with number of skips). Finally, only 45 game sessions (3.2%) failed at least one controlled trial (**Fig. S6D**) and were subsequently removed from the triplet dataset.

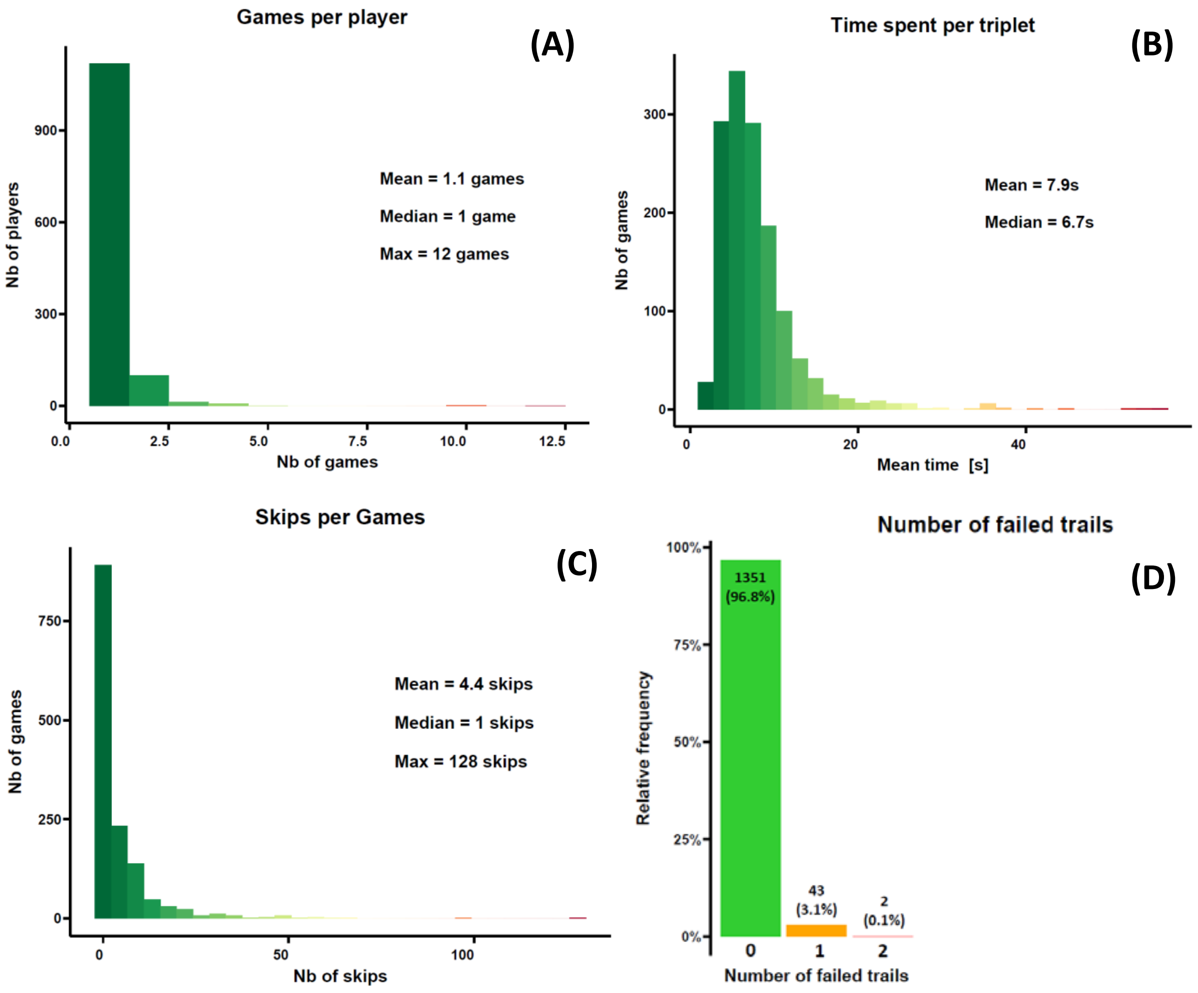


**Figure S6: Analytics for game sessions. (A)** Number of games per player. **(B)** Mean time spent examining each triplet in a game session. **(C)** Number of uses of the 'skip' option per game session. **(D)** Percentage of game sessions according to the number of failed controlled trials in the game session.

**Player diversity: age, background, geography, language**

We reached a diverse audience of human surrogate ‘predators’ with declared ages ranging from 4 years old to 92 years old. The distribution of age shows the survey reached mostly adults, likely students, researchers, and their family members (**Fig. S7A**). The four different languages of the interface were used with varying degrees of success. The Spanish and Portuguese interfaces were rarely used (< 4%; **Fig. S7B**), also because native speakers who played were researchers familiar with English who likely used the English interface. Most players had a biological scientific background (785 players = 55.2%; **Fig. S7C**) due to the use of scientific diffusion channels to advertise the game. However, having almost half of players who do not have a biological scientific background is a nice pointer to the fairly efficient diffusion of the survey outside of the scientific community despite not having been targeted during diffusion.

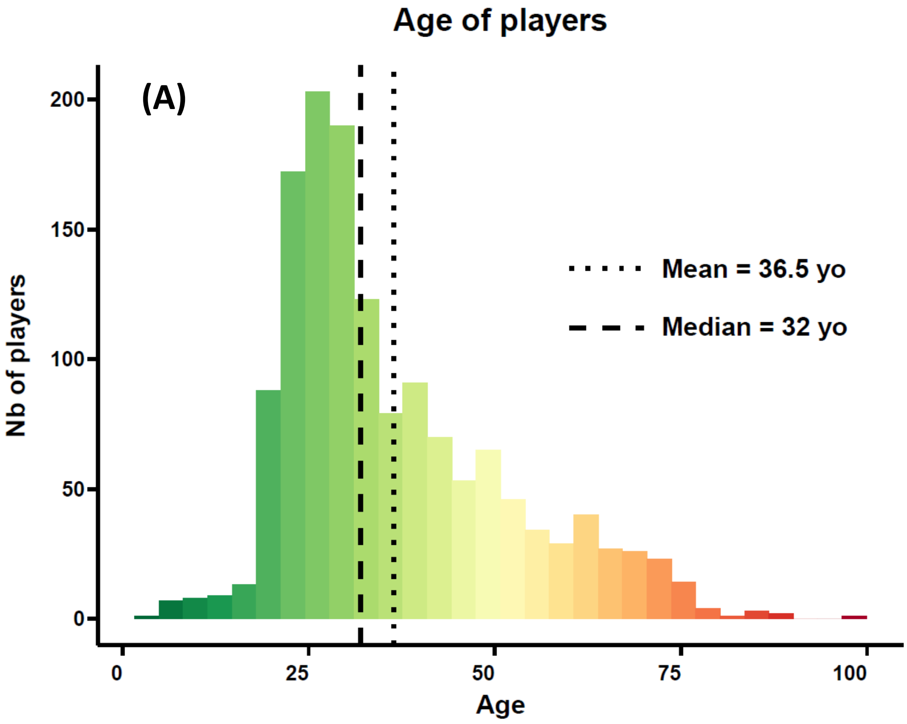

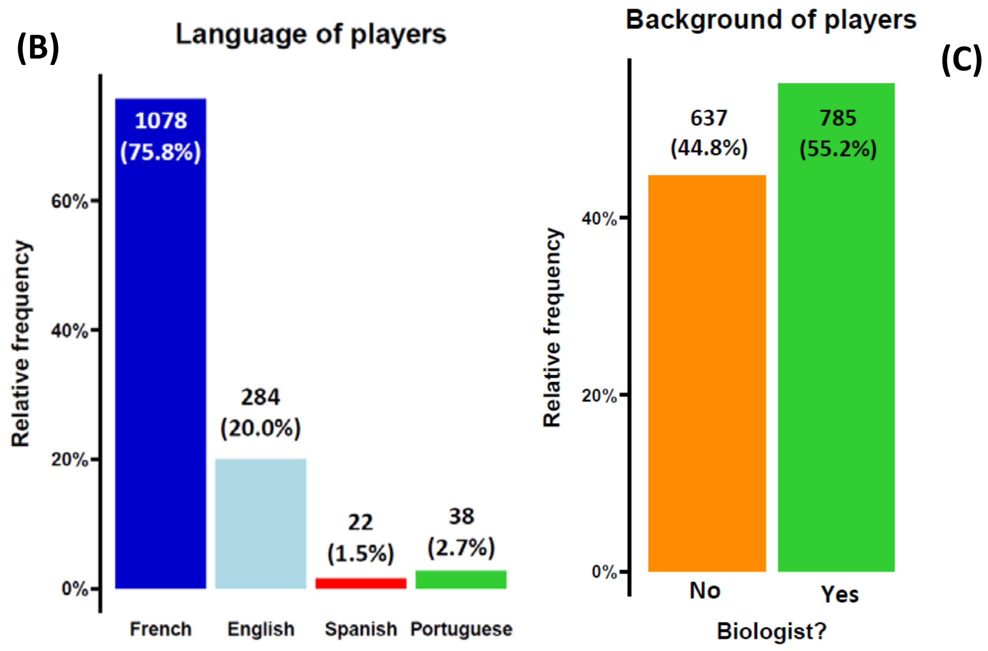


**Figure S7: Analytics for player demographics. (A)** Distribution of age of players. **(B)** Language interface used in game sessions. **(C)** Biological scientific background of players.

#### Normality scores:

In order to evaluate the dissimilarities across game sessions, we designed two normality scores reflecting the degree of agreement between the set of similarity triplets provided during a game session, and the final perceptual space built on all similarity triplets. The triplet-based score simply quantifies the percentage of triplets whose distance between images A and B, perceived as the most similar pair, is indeed inferior to the distance between images A and C in the final perceptual space (i.e., $d_{A-B} > d_{A-C}$). The distance-weighted score adjusts the triplet-based score according to relative distances of the images in each triplet as follows:

$$S_{triplet} = \min(\frac{d_{A-C}}{d_{A-B}}, 1) \quad \text{(Eqn. 4)}$$

where $d_{A-B}$ is the distance between the images perceived as the most similar and $d_{A-C}$ is the distance between the images not labeled as the most similar. As such, satisfied triplets score one and unsatisfied triplets with a high relative difference in pairwise distances score lower to account for the high abnormality of this triplet compared to the global perceptual space. The score for a game session is the sum of the score for each triplet. It ranges from 0 to the number of triplets. Final scores were converted in percentages. Importantly, they are not scores of performance but scores of similarity to the norm, thus we coined them as “normality scores”.

In average, a game session provided 86.1% of similarity triplets in agreement with the coordinates of images in the final perceptual space built on all similarity triplets (**Fig. 8A**). Once weighted for relative differences in distances, the mean score increases to 96.3% (**Fig. 8B**) which hints that the most unsatisfied triplets show a small relative distance in the final perceptual space, reflecting small variation in perception.

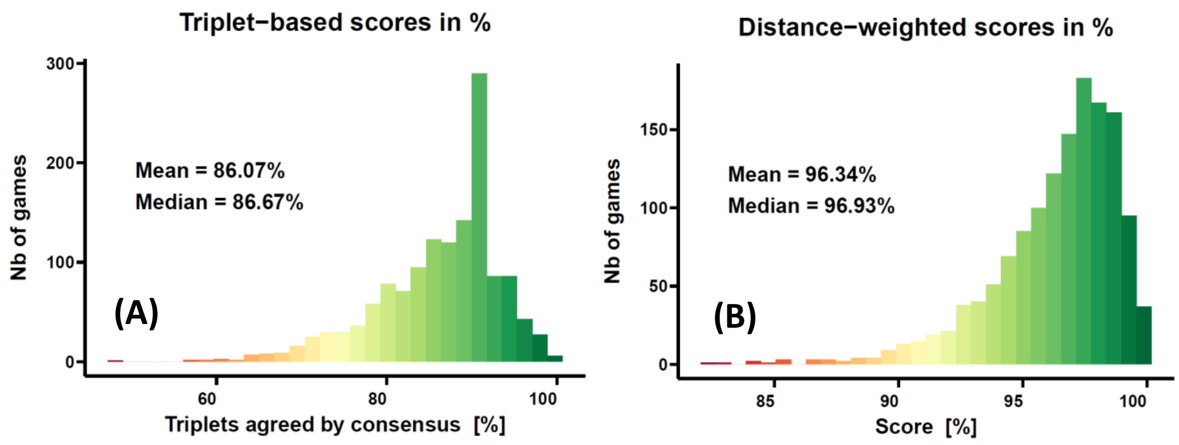


**Figure S8: Distribution of normality scores across games. (A)** Triplet-based score evaluating the percentage of triplets provided during a game session in accordance with the final perceptual space produced using all similarity triplets. **(B)** Distance-weighted score adjusting the triplet-based score according to pairwise distances of the image in each triplet. Unsatisfied triplets with a high relative difference in pairwise distances score lower to account for the high abnormality of this triplet compared to the global perceptual space.

#### Effects of skip option and control trials:

Using the distance-weighted normality scores, we explored the effect of the skip option and the number of failed control trials on the score of a game session. We showed the usefulness of the skip option by highlighting a positive correlation between the number of skips and the normality score (Spearman's $\rho = 0.292$, $p < 0.001$; **Fig. S9A**) reflecting a higher number of abnormal triplets recorded in games with low to no skip uses. Games with failed control trials appeared to score significantly lower than games without fails (Kruskal-Wallis, $\chi = 59.9$, $p < 0.001$; **Fig. S9B**) illustrating the ability of control trials to detect abnormal game sessions. Altogether, both features seem to fulfill their purpose of filtering triplets and game sessions which are strong suspicious outliers relative to the global perception of players.

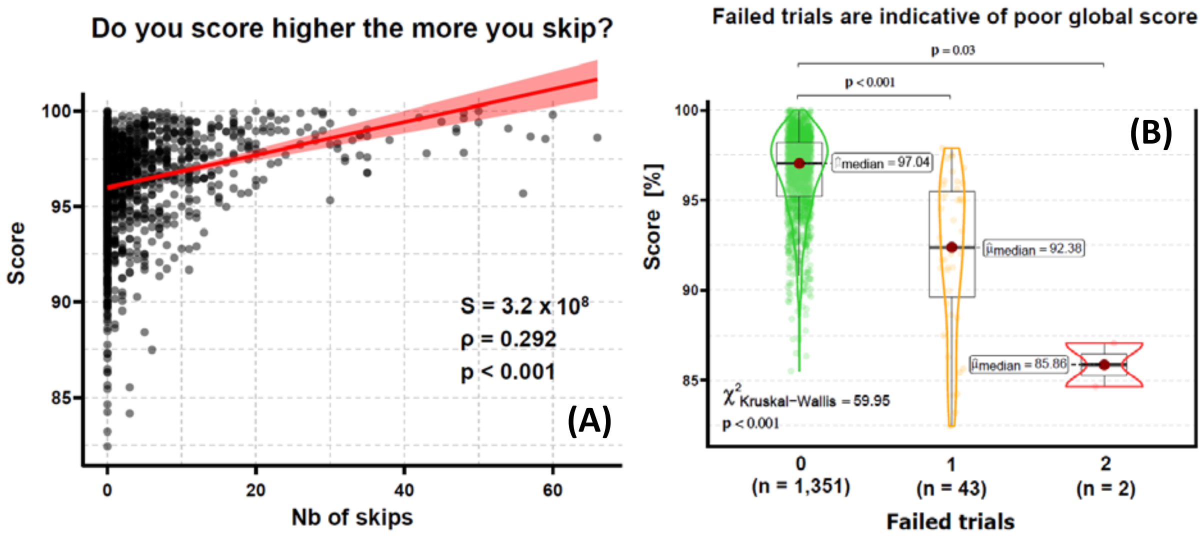


**Figure S9: Tests for the effects of the skip option and the number of failed control trials on normality scores. (A)** Positive correlation between normality scores and number of skips in a game session. **(B)** Negative relationship between normality scores and number of failed controlled trials in a game session.

#### Effects of Age and Colorblindness:

We explored the effect of age and colorblindness on game performance. Age did not have any significant effect on normality scores (Spearman's $\rho = 0.009$, $p = 0.741$; **Fig. S10A**), neither on the mean time taken to answer a triplet (Spearman's $\rho = -0.021$, $p = 0.431$; **Fig. S10B**). However, older players tended to use more the skip option than younger players (Spearman's $\rho = 0.077$, $p = 0.033$; **Fig. S10C**). This mild trend could reflect the idea that older players trust their perceptual abilities less or are more careful in their choices. A low number of colorblind people participated in the survey (N = 17). Such a low number of games did not allow to find a statistically significant difference in normality scores, however we visualized a slight tendency for the few colorblind players to score lower, reflecting their particular visual perception (Mann-Whitney, W = 9395, $p = 0.16$; **Fig. S10D**).

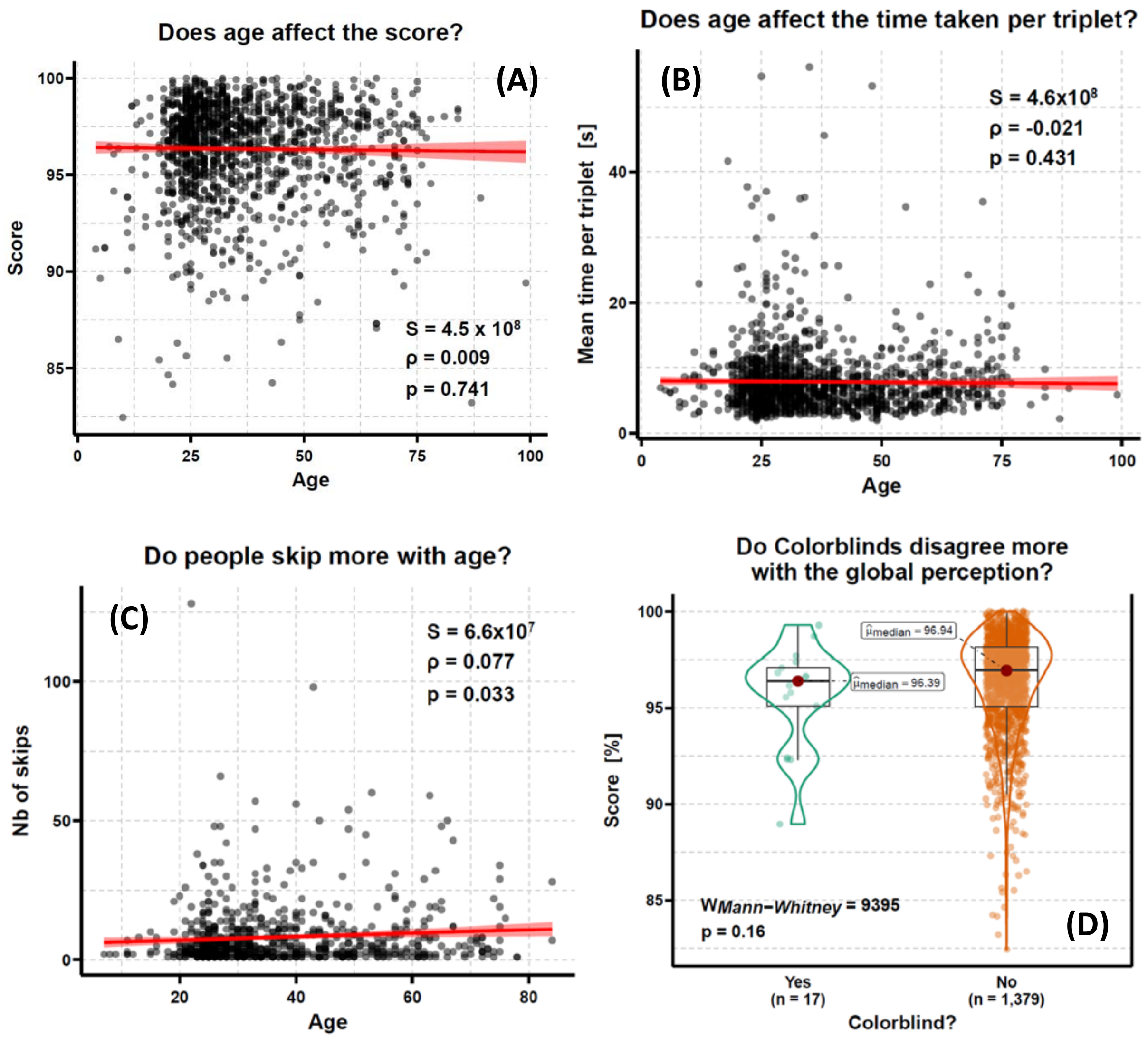


**Figure S10: Tests for the effects of age and colorblindness on game performances. (A)** No effect of age on normality scores. **(B)** No effect of age on the mean time taken to answer a triplet. **(C)** Positive correlation between age and number of skips. **(D)** No significant effect of colorblindness on normality score, yet we detect a slight tendency for the few colorblind players to score lower reflecting their particular visual perception.

#### Effects of cultural and scientific background:

We explored the effect of cultural and scientific background on game performance. Biologists appeared to play quicker (Mann-Whitney: W = 21,812, $p < 0.001$; **Fig. S11A**) and score significantly higher than non-biologists (Mann-Whitney: W = 29,743, $p < 0.001$; **Fig. S11C**), but they did not show any difference in their frequency of use of the skip option (Mann-Whitney: W = 66,464, p = 0.48; **Fig. S11B**). Since scores are normality scores and not performance scores, these results mostly tell us that experience affects the perception of similarity in wing patterns and that non-biologists tend to perceive similarity patterns that are outside the norm more frequently than trained biologists are. In parallel, Brazilian-Portuguese speakers were scoring significantly higher than other languages (Kruskal-Wallis: $\chi = 14.7$, $p < 0.001$; **Fig. S11D**). This trend is likely due again to the effect of experience: almost all respondents for this language were scientists working with neotropical butterflies, native to their country (i.e., Brazil). Altogether, these results hint for a strong effect of experience in task requiring abilities to detect differences in neotropical butterflies. However, this conclusion needs to be tempered by the significantly lower scores obtained by Spanish speaking speakers, mostly from Latin America, for which the same positive effect of experience could have been expected (**Fig. S11D**).

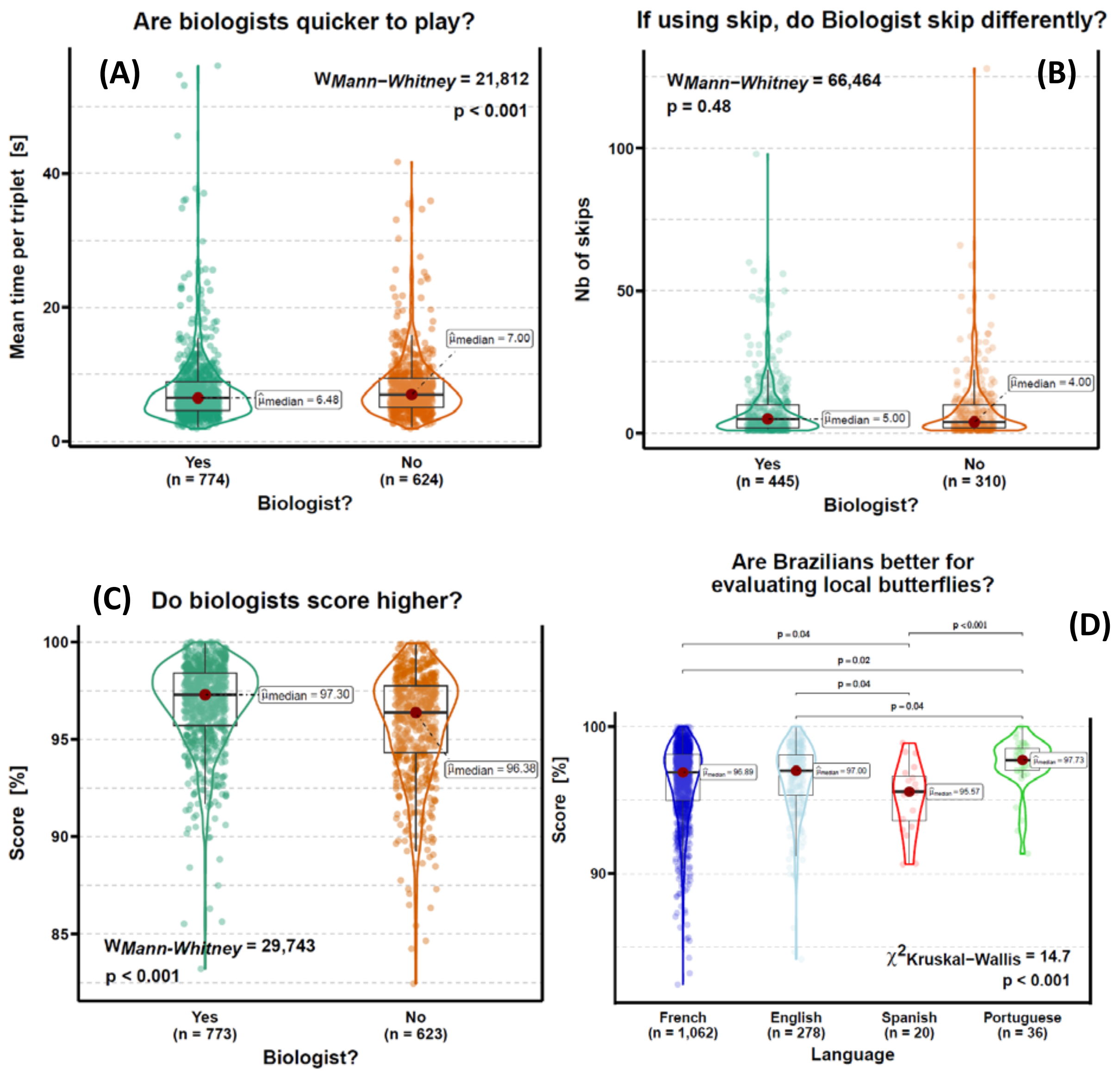


**Figure S11: Tests for the effects of cultural and scientific background on game performances. (A)** The scientific background affects the response time: biologists are significantly faster to answer. **(B)** No effect of scientific background on the use of the skip option. **(C)** Positive relationship between scientific background and normality scores: biologists score higher. **(D)** Significant effect of the language on normality scores with Brazilian-Portuguese speakers scoring significantly higher than other languages, hinting for an effect of experience in task requiring abilities to detect differences in native neotropical butterflies.

**Effect of learning:**

We explored the effect of learning through multiple game sessions on game performance. Globally, first game sessions were slower than the next game sessions (Mann-Whitney: W = 17,205, $p < 0.001$; **Fig. S12A**). This effect persisted when comparing for a given player the time taken during the first game session and the average time for the next game sessions (Wilcoxon V = 3,42848, $p < 0.001$; **Fig. S12B**). As such, players seem to learn to be more efficient (and maybe more confident) at selecting the pair they perceived as the most similar, highlighting the importance of memorization and experience in perception. Globally, first game sessions provide lower normality scores than the next game sessions (Mann-Whitney: W = 11,928, $p = 0.003$; **Fig. S12C**). However, at player-level, we detected no significant increase of the normality score between the first game session and the average of all next sessions (Wilcoxon V = 2,865, $p = 0.24$; **Fig. S12D**). Thus, players who played the most were the ones the least susceptible to showing a perception out of the ordinary, but they did not seem to provide similarity triplet closer to the global average as much as they played. Altogether, we did not show a clear effect of training experience on the evolution of perception towards the norm during game sessions, but trained players definitely played faster.

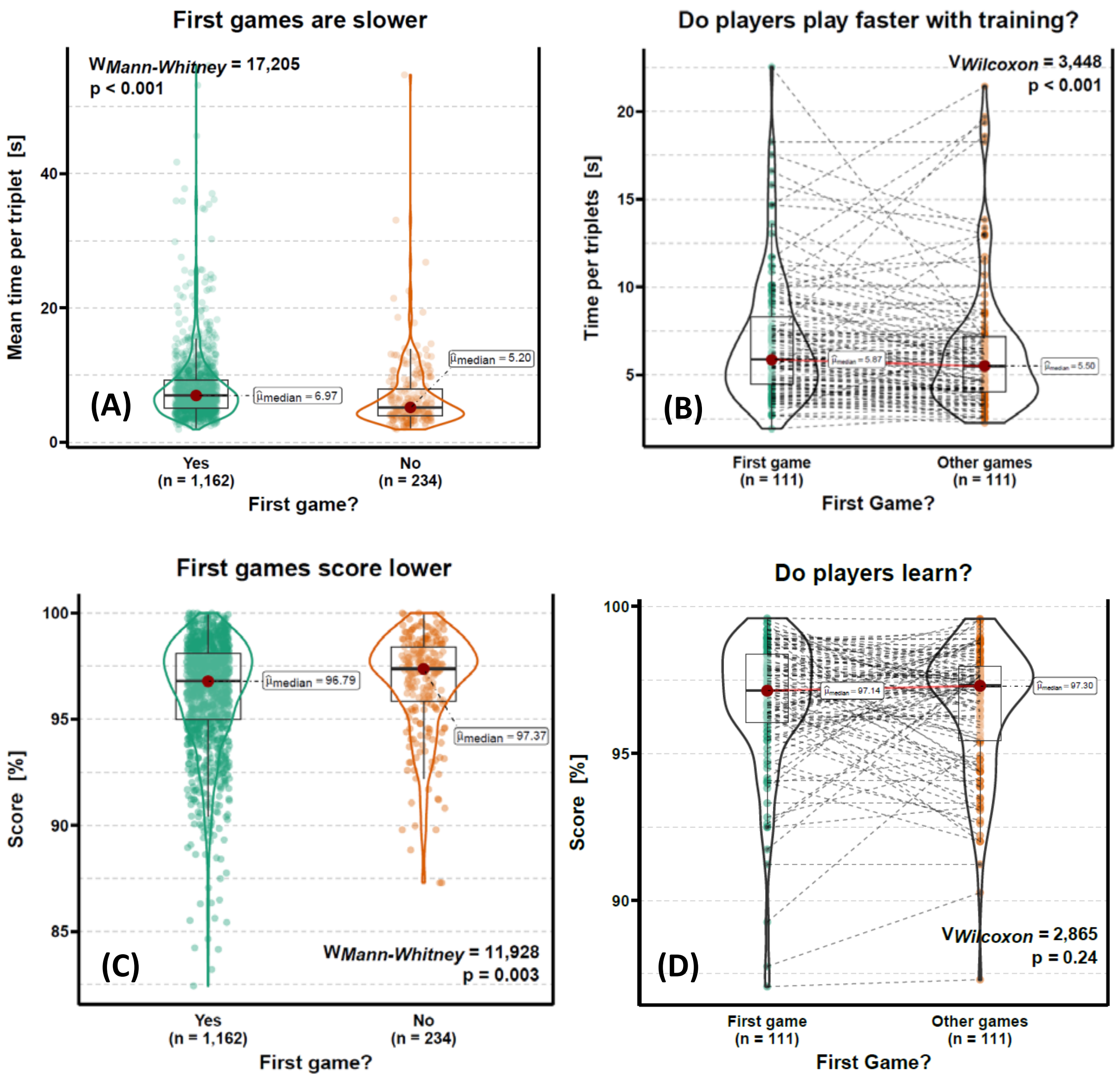


**Figure S12: Tests for the effects of learning through multiple game sessions on game performance. (A)** Globally, the first game sessions are slower than the next game sessions. **(B)** At the player level, the first game sessions are slower than the next game sessions. **(C)** Globally, the first game sessions provide lower normality scores than the next game sessions. **(D)** At player-level, there is no significant increase of the normality score between the first game session and the average of all next sessions.

### Appendix 5: Visual lists of subspecies in local communities

We produced analyses of perceived wing pattern variation in five local communities. We present here the subspecies found in each of this community: Cayenne, French Guyana (**Fig. S13**), Gamboa, Panamá (**Fig. S14**), Jatun Sacha, Napo, Ecuador (**Fig. S15**), Manaus, Amazonas, Brazil (**Fig. S16**), and Santa Teresa, Espírito Santo, Brazil (**Fig. S17**).

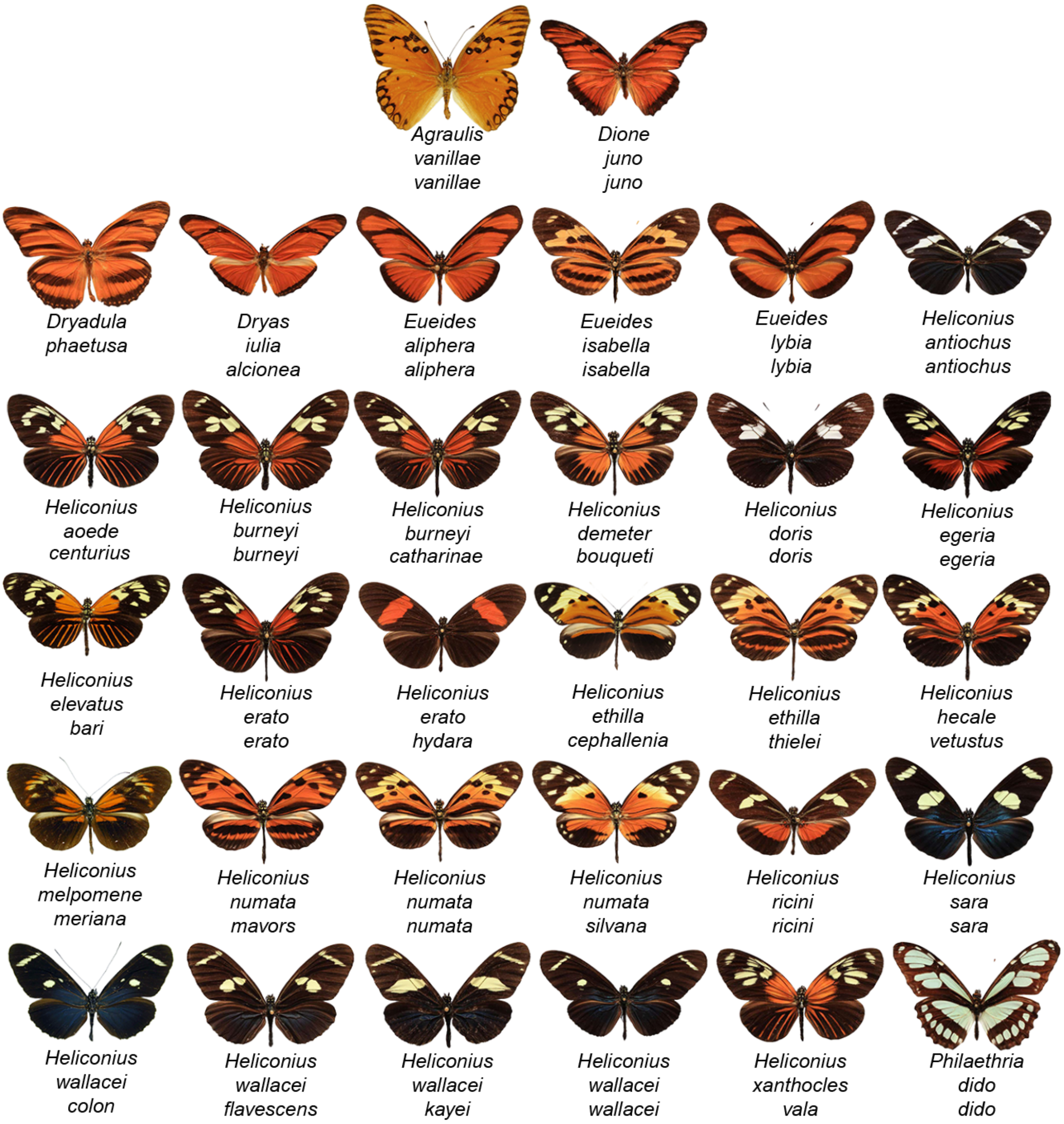


**Figure S13: Visual taxonomic list of the 32 subspecies of heliconiine butterflies found in Cayenne, French Guyana.**

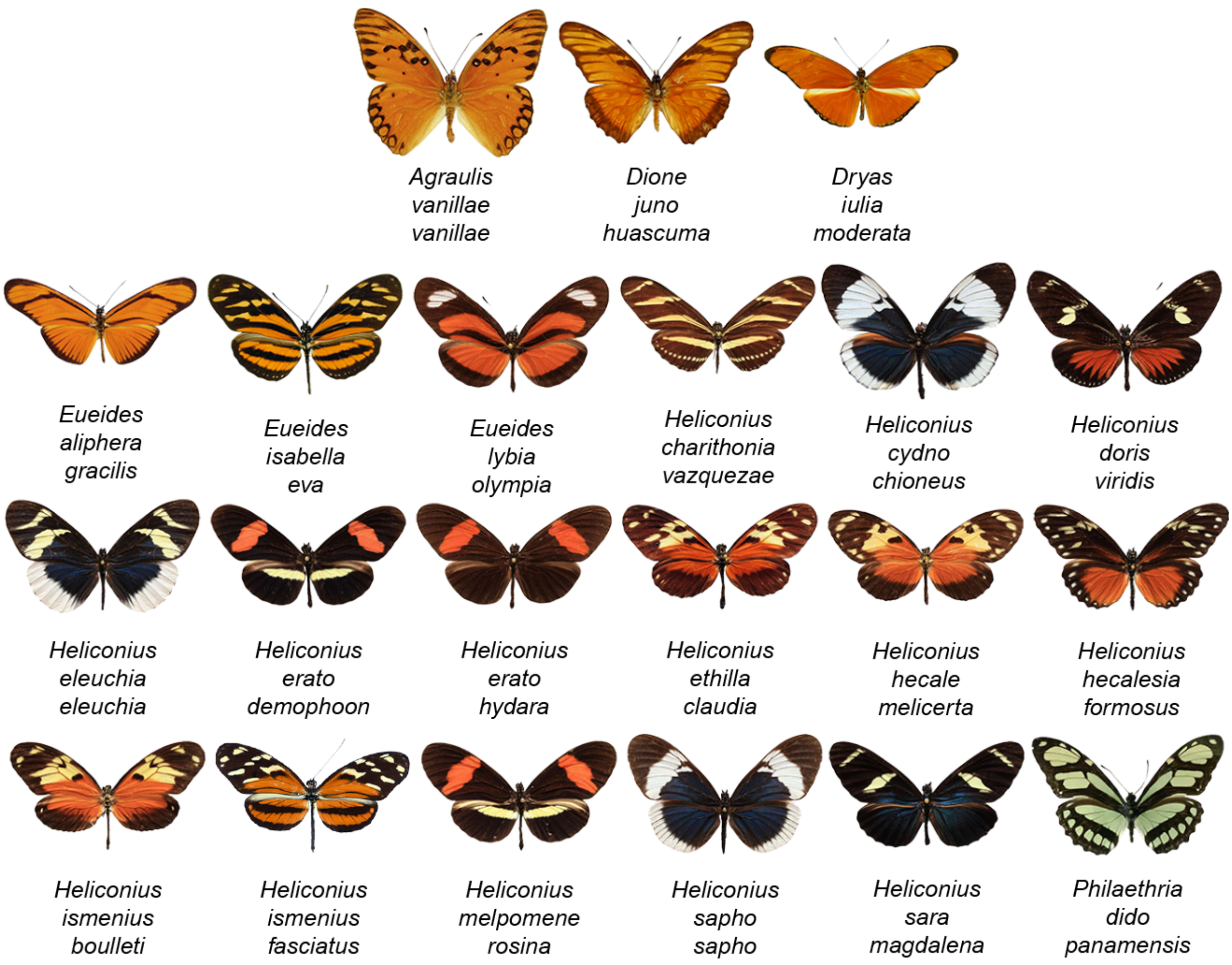


**Figure S14: Visual taxonomic list of the 21 subspecies of heliconiine butterflies found in Gamboa, Panamá.**

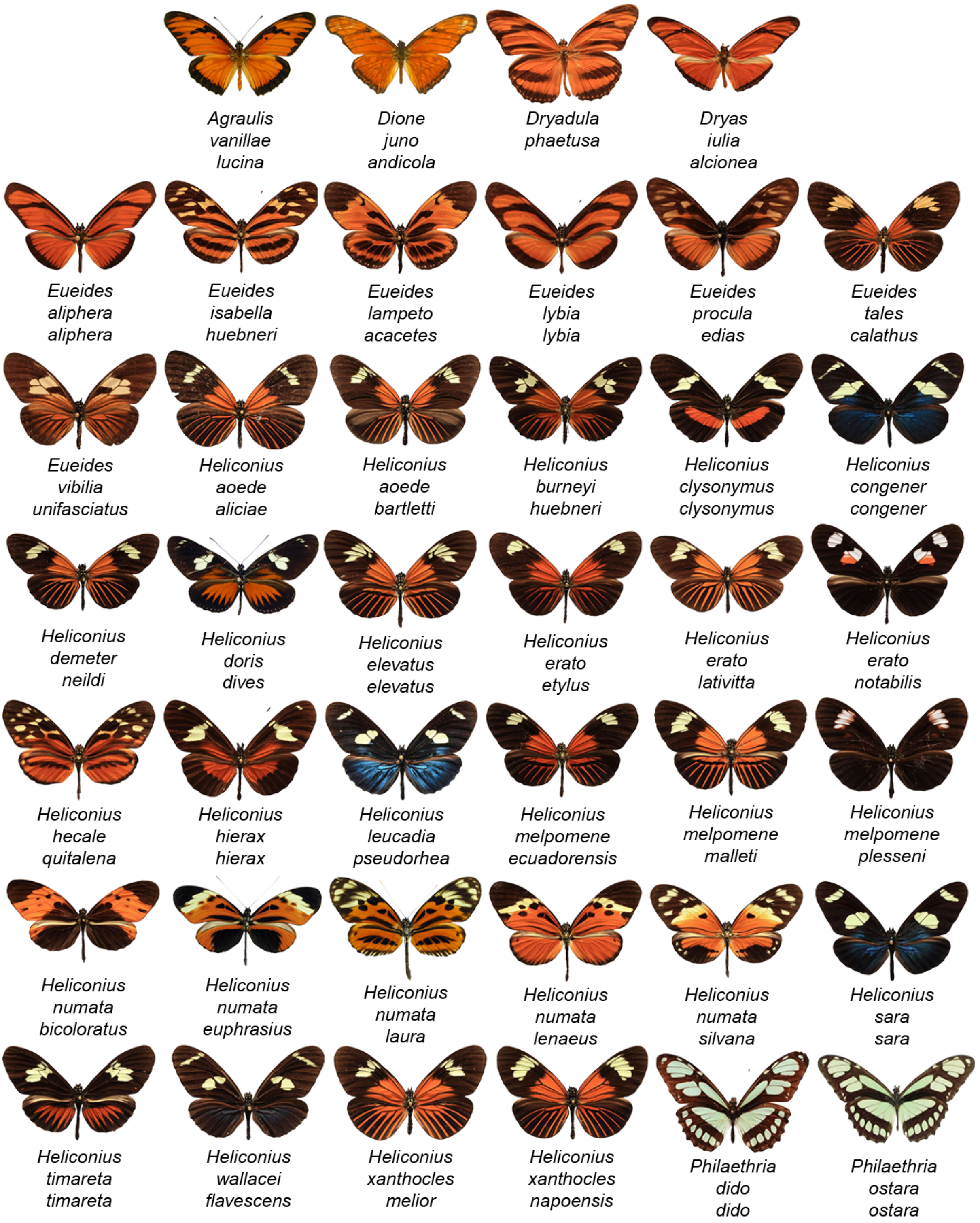


**Figure S15: Visual taxonomic list of the 40 subspecies of heliconiine butterflies found in Jatun Sacha, Napo, Ecuador.**

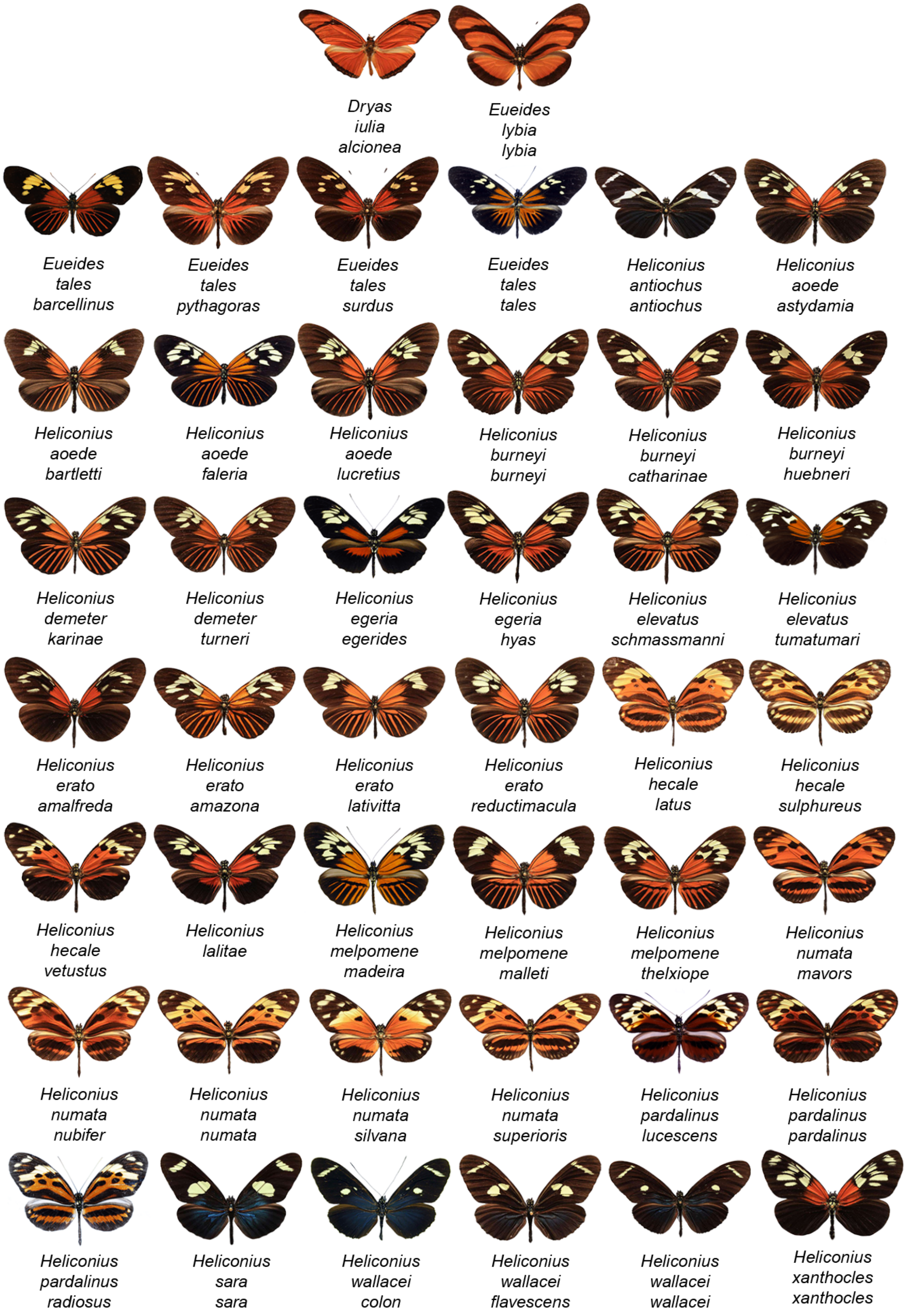


**Figure S16: Visual taxonomic list of the 44 subspecies of heliconiine butterflies found in Manaus, Amazonas, Brazil.**

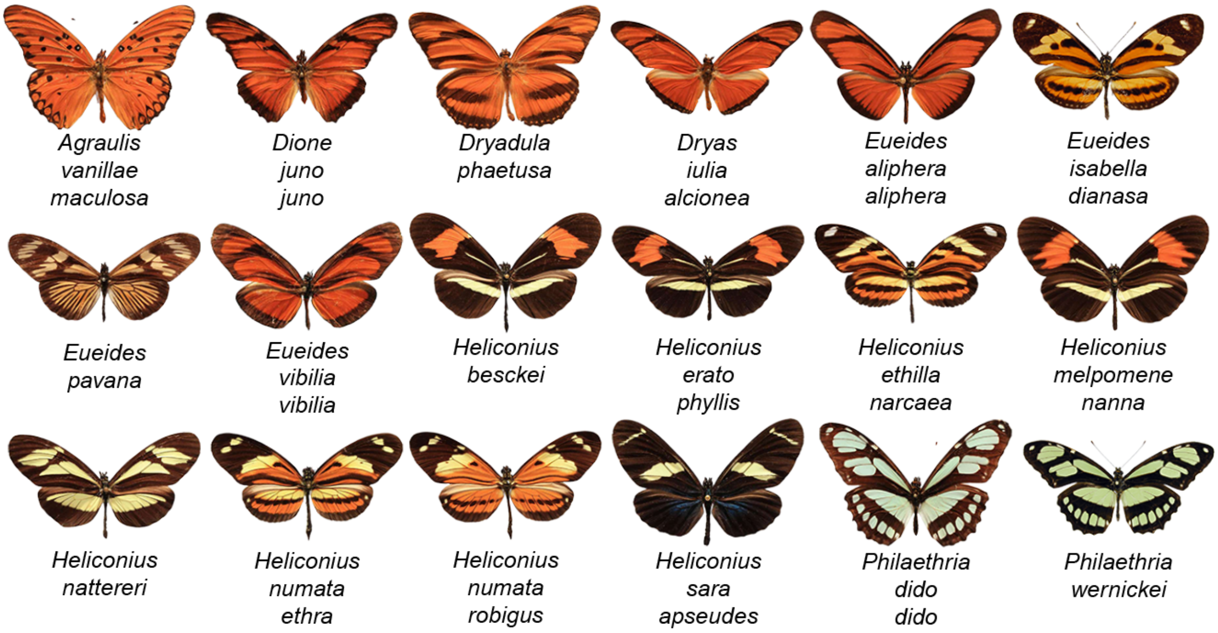


**Figure S17: Visual taxonomic list of the 18 subspecies of heliconiine butterflies found in Santa Teresa, Espírito Santo, Brazil.**

### Appendix 6: Perceptual spaces of local communities

To test whether the citizen science dataset obtained with the full set of phenotypes accurately reflects perceived similarity at the local scale, we independently generated perceptual maps derived from evaluations by a panel of participants who assessed wing pattern similarity solely among taxa found within each of five local communities (labeled as 'local reference' maps). Those maps were then compared to maps of the same five local sets of wing patterns extracted from the comprehensive perceptual space based on citizen science data (referred to as 'extracted' maps) (*47*).

'Local reference' maps and 'extracted' maps of the five local communities are displayed below (**Fig. S18**), including perceptual maps of Cayenne and Santa Teresa, omitted in the main text (**Fig. 2**).

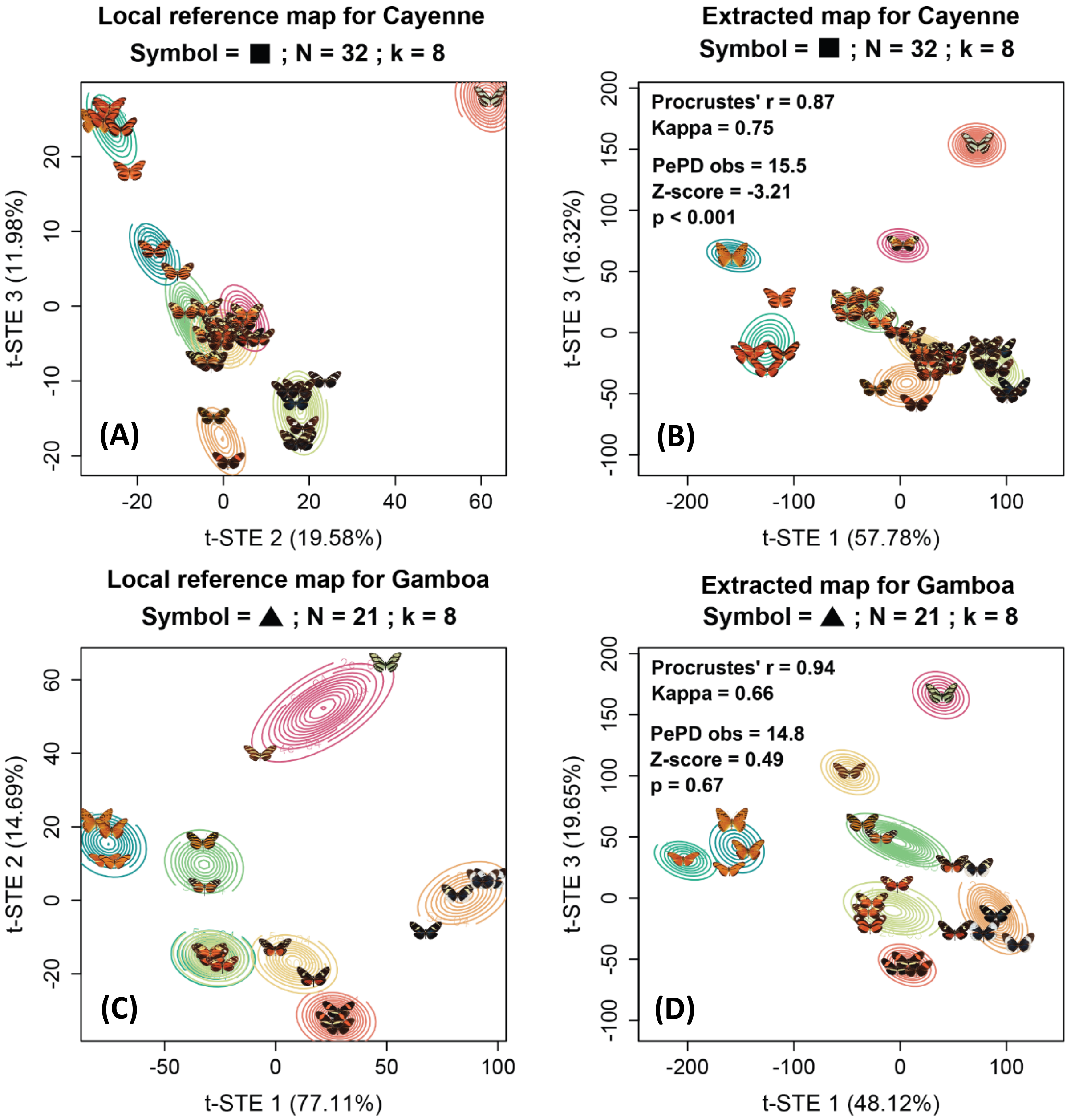

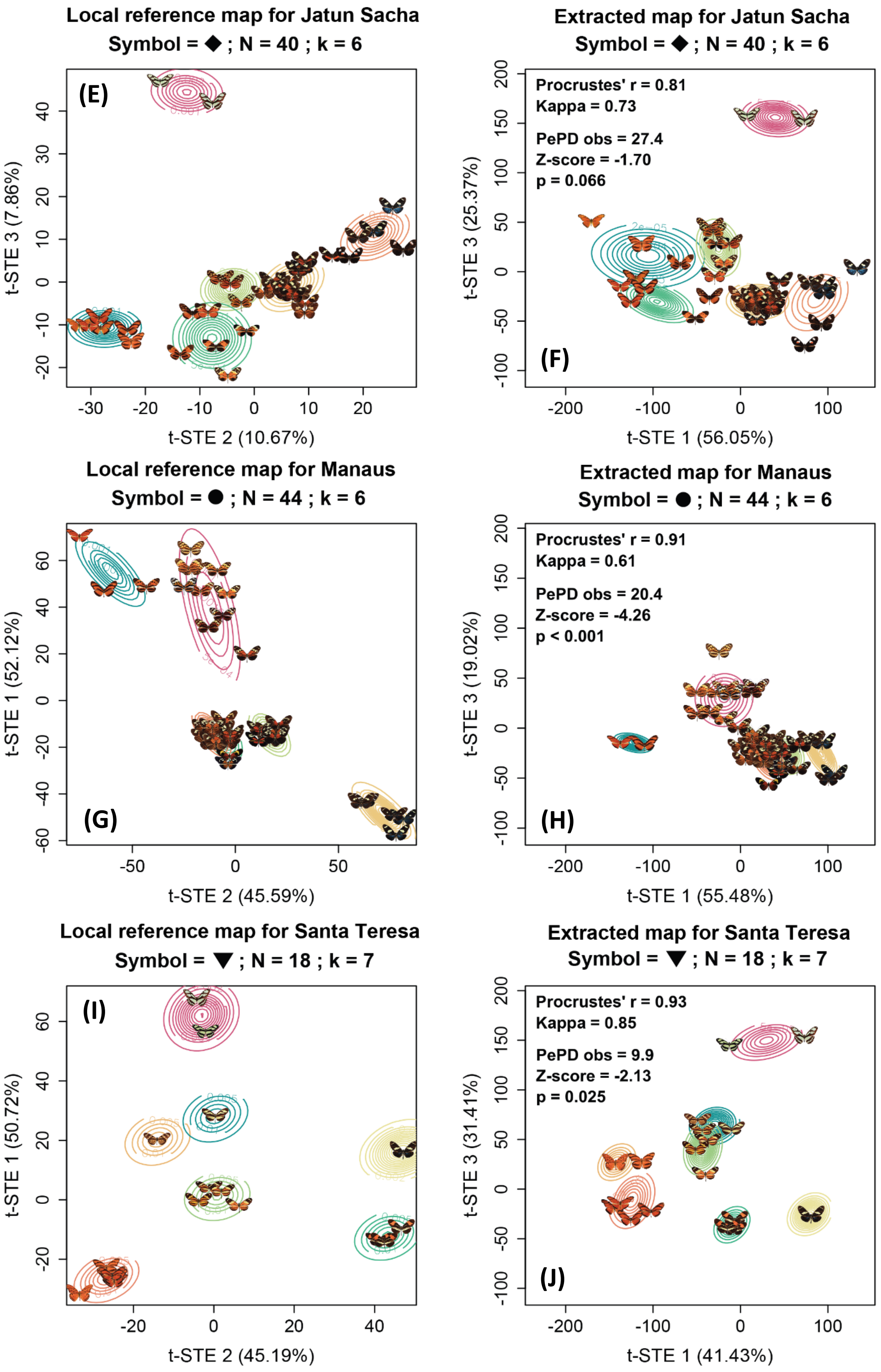
Local reference map for Jatun Sacha
Symbol = ◆ ; N = 40 ; k = 6
(E)
t-STE 3 (7.86%)
t-STE 2 (10.67%)
Extracted map for Jatun Sacha
Symbol = ◆ ; N = 40 ; k = 6
Procrustes' r = 0.81
Kappa = 0.73
PePD obs = 27.4
Z-score = -1.70
p = 0.066
(F)
t-STE 3 (25.37%)
t-STE 1 (56.05%)
Local reference map for Manaus
Symbol = ● ; N = 44 ; k = 6
(G)
t-STE 1 (52.12%)
t-STE 2 (45.59%)
Extracted map for Manaus
Symbol = ● ; N = 44 ; k = 6
Procrustes' r = 0.91
Kappa = 0.61
PePD obs = 20.4
Z-score = -4.26
p < 0.001
(H)
t-STE 3 (19.02%)
t-STE 1 (55.48%)
Local reference map for Santa Teresa
Symbol = ▼ ; N = 18 ; k = 7
(I)
t-STE 1 (50.72%)
t-STE 2 (45.19%)
Extracted map for Santa Teresa
Symbol = ▼ ; N = 18 ; k = 7
Procrustes' r = 0.93
Kappa = 0.85
PePD obs = 9.9
Z-score = -2.13
p = 0.025
(J)
t-STE 3 (31.41%)
t-STE 1 (41.43%)

**Figure S18: Perceptual maps of phenotypes within local communities, displayed with GMM probability densities for 'local reference' and 'extracted' maps.** Contour lines represent the probability densities of GMM clusters defining local mimicry rings**.** Only two axes among the three are displayed. 'Local reference' maps **(A-C-E-G-I)** were built from similarity evaluation using triplets based only on local patterns and provided by five survey participants. 'Extracted' maps **(B-D-F-H-J)** were obtained from the perceptual space (**Fig. 1**) built from citizen science survey responses on triplets drawn from the entire set of Heliconiini images. Axes t-STE 1 and t-STE 3 for the 'extracted' maps correspond to the axes shown in **Fig. 1C**. Axes shown in 'local reference' maps were selected to represent similar pattern trends relative to the axes shown in 'extracted' maps. Procrustes correlation measures similarity in map topology between 'extracted' and 'local reference' maps. Cohen's Kappa measures agreement in GMM classifications. Perceived Phenotypic Diversity (PePD) quantifies the coverage of local phenotypes in the perceptual space based on Gaussian density kernels, here reported as percentage of the global coverage. The Z-score compares observed PePD with PePD obtained from a model of neutral evolution of wing patterns. Negative values indicate evolutionary convergence. P-values are based on quantiles of observed values among simulated data. N = number of local patterns. k = chosen number of hypothesized mimicry rings. A 3D animated version of the perceptual space for 'local reference' of each community is available in SI for the online version.

#### Appendix 7: RGB maps of mean phenotype in local communities

By converting the perceptual space in an RGB color space, we assigned a color to each community based on the coordinates of its mean phenotype. The distribution of communities in the perceptual space showed that no community displayed a mean phenotype falling in the region of dark patterns with high t-STE 1 and low t-STE 2 and t-STE 3 (**Figs. S19A, S19C, S19D**). In the geographic space, the most species-rich regions of the Andes and Amazonia presented, by construction, a mean phenotype close to the centroid of the perceptual space symbolized by a brownish tone (**Fig. S19B**). The Cerrado and Brazilian Atlantic forest demonstrated a trend towards a majority of orange and tiger patterns symbolized by the reddish tone (**Fig. S19B**), illustrated by the relatively marginal position of local community of Santa Teresa in the RGB space (down-triangle symbol in **Fig. S19**). North America was clearly an outlier with its orange tone, having a mean phenotype falling among the plain orange patterns since the few heliconiine species found in this area are from the *Agraulis, Dione, Dryas* and *Dryadula* genera all displaying an orange-based pattern, with the notable exception of the stripped *Heliconius charithonia* (**Fig. S19B**).

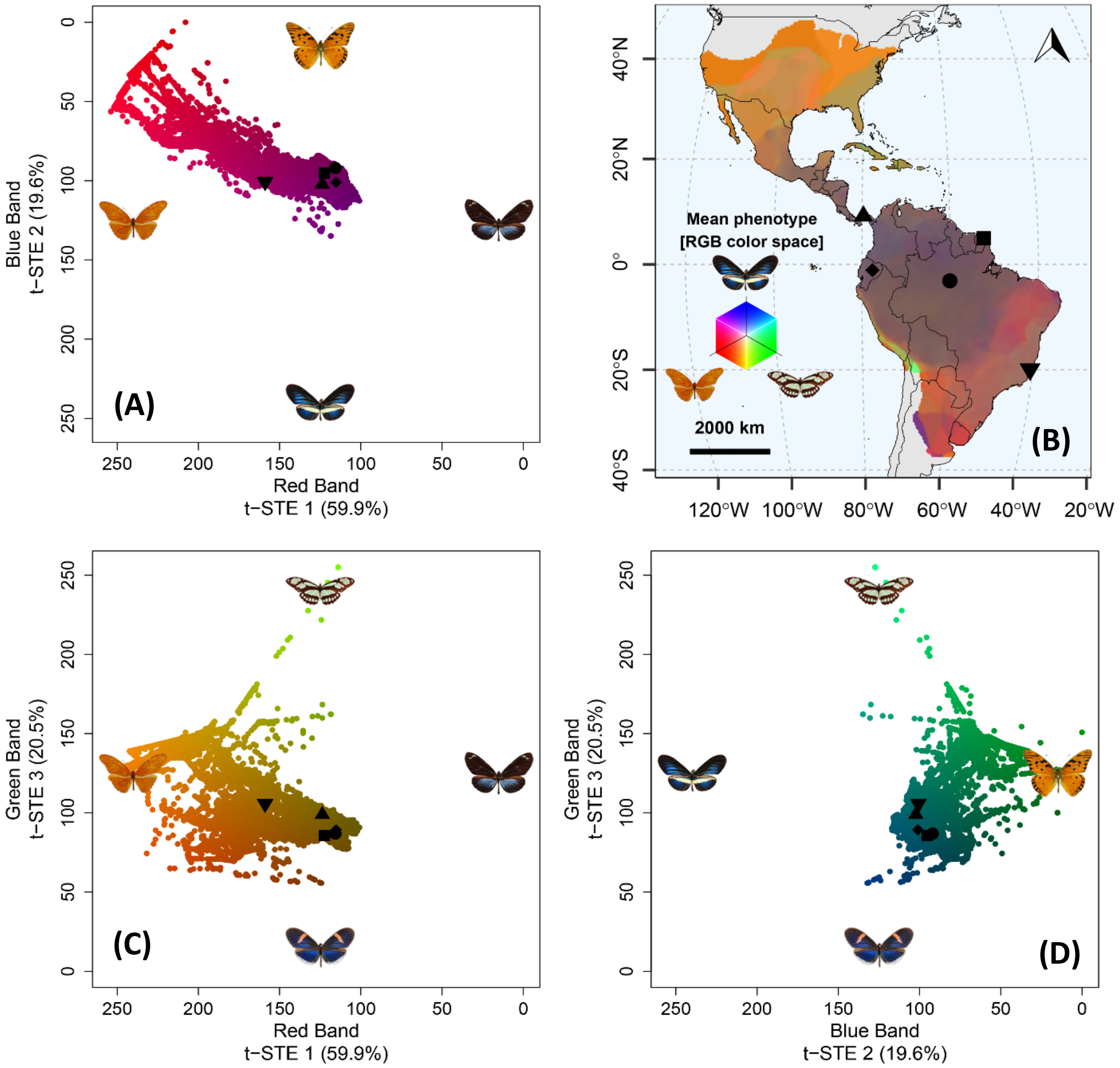


**Figure S19: RGB maps of local mean phenotype built from citizen science data. (A-C-D)** 2D RGB maps displaying the mean phenotype found in the 23,661 communities defined as 30 km x 30 km grid cells within the range of heliconiine butterflies. Axes range from 0 to 255 scaled on the range of the coordinates of the subspecies in the initial perceptual space (**Fig. 1**). Distances between communities reflect global perception of dissimilarity in the mean local phenotype. Phenotypes shown at each extremum of the perceptual space belong to the heliconiine subspecies the closest to the extremum. As such, they illustrate the changes in community mean patterns along each axis of the perceptual RGB space. **(B)** Geographic map of mean local phenotype whose color corresponds to its coordinates in the RGB space. Differences in color relate to dissimilarity in mean local phenotypes. Symbols represent local communities highlighted in **Figure 2**: square = Cayenne, up-triangle = Gamboa, diamond = Jatun Sacha, circle = Manaus, down-triangle = Santa Teresa.

#### Appendix 8: RGB maps of perceived phenotypic β-diversity across local communities

Complementarily to our map of mean local phenotypes (**Fig. S19**), we now accounted in the variance in local phenotypes when mapping perceived phenotypic β-diversity in heliconiine wing patterns across communities. As such, we computed the Mahalanobis distances between all pairs of communities based on the whole sets of coordinates of their respective local phenotypes in the global perceptual space. Appling Non-metric MultiDimensional Scaling (NMDS), we built a new 3D space where distances between communities relate to their dissimilarities in perceived phenotypic composition as estimated from Mahalanobis distances. We converted this phenotypic composition space into an RGB color space and assigned a color to each community based on its coordinates (**Fig. S20)**. This RGB space is not a perceptual phenotypic space: a point does not correspond to a unique phenotype (contrary to **Fig. 2** and **Fig. S18**), but to a set of phenotypes found in a local community.

The Amazon basin presented a trend for community dominated by species with black and red rays (**Fig. S20B**, in red tones) different from the communities in the Brazilian Atlantic Forest (**Fig. S20B**, in green tones) where striped black-and-orange "tiger" patterns and patterns with large red band on the forewings appeared more prominent, as illustrated by the relatively marginal position of local community of Santa Teresa in the RGB space (down-triangle symbol in **Figs. S19 & S20**). North America was again clearly an outlier (**Fig. S20B**, in blue tones), having most local phenotypes falling among the plain orange patterns. Overall, the geographic transposition of dissimilarities in perceived phenotypic composition of communities (i.e., perceived phenotypic β-diversity) highlight the important differences across bioclimatic regions such as the Amazon basin, the Brazilian Atlantic Forest and North America (**Fig. S20B)**.

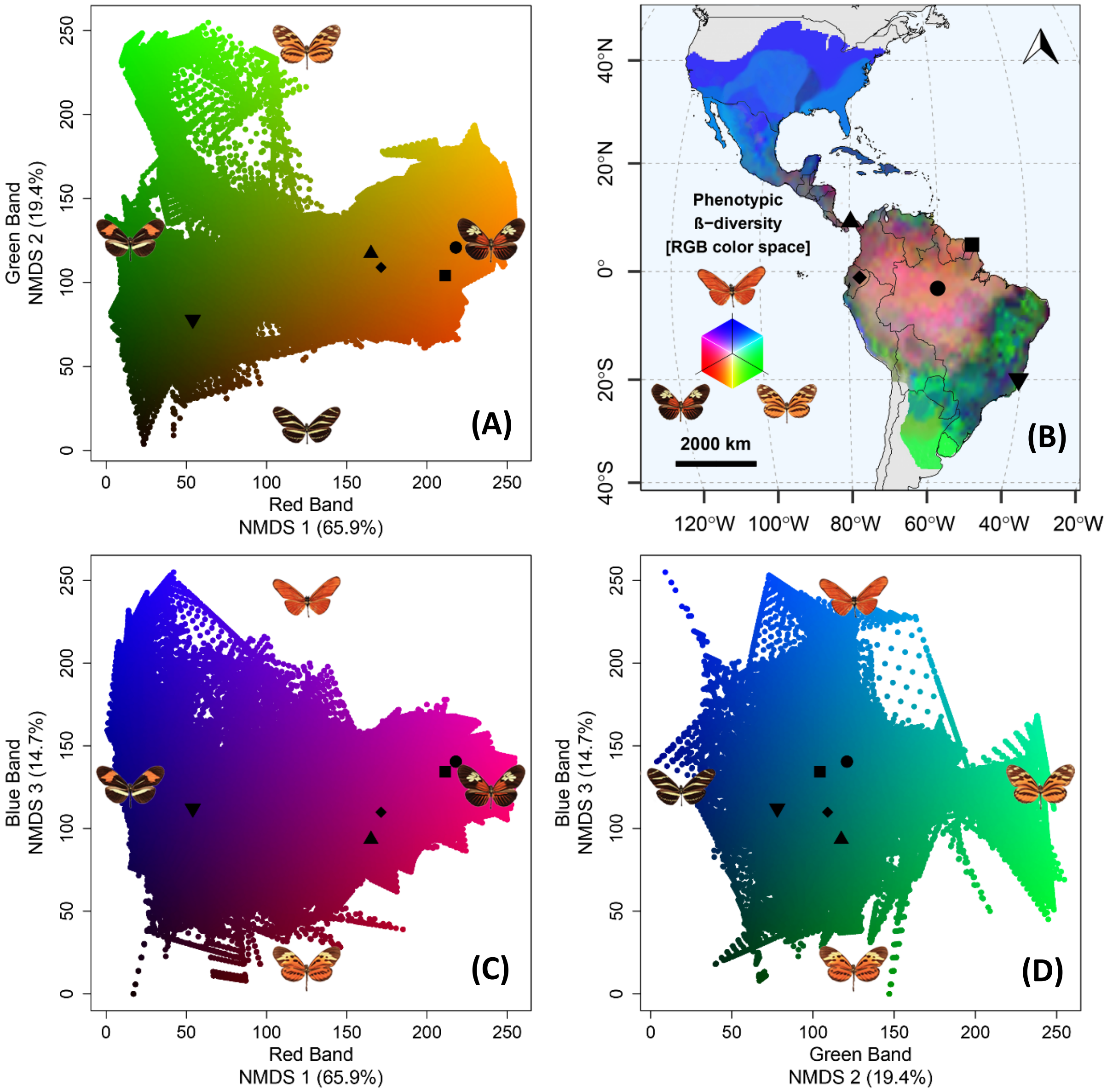


**Figure S20: RGB maps of perceived phenotypic β-diversity across local communities.** Maps were built from NMDS applied to pairwise Mahalanobis distances computed from the coordinates of local patterns of communities in the citizen science-based perceptual space. **(A-C-D)** 2D RGB maps displaying the dissimilarity in perceived phenotypic diversity across the 23,661 communities defined as 30 km x 30 km grid cells within the range of heliconiine butterflies. Axes range from 0 to 255 to allow conversion in RGB colors. Distances between communities reflect the global perception of dissimilarity between whole sets of local phenotypes. Phenotypes shown at each extremum of the perceptual space belong to the heliconiine subspecies the most representative of the set of patterns found in communities the closest to the extremum. As such, they illustrate the changes in community pattern compositions along each axis of the RGB space. This RGB space is not a phenotypic space: a point does not correspond to a unique phenotype (contrary to **Fig. 2** and **Fig. S18**), but to a set of phenotypes found in a local community. **(B)** Geographic map of perceived phenotypic β-diversity across communities whose color corresponds to their coordinates in the RGB space of perceived phenotypic composition. Differences in color relate to perceived dissimilarity in sets of local phenotypes**.** Symbols represent local communities highlighted in **Figure 2**: square = Cayenne, up-triangle = Gamboa, diamond = Jatun Sacha, circle = Manaus, down-triangle = Santa Teresa.

### Appendix 9: Tests for effect of sympatry on perceptual distances

Müller's theory of mimicry (*3*) predicts the local convergence of defended prey towards a similar aposematic pattern. To evaluate if this prediction could influence large-scale patterns of phenotypic distributions, we tested if the perceptual distances between subspecies were correlated with the geographic distances between their spatial distributions.

We used Multiple Regression on distance Matrices (MRM) to test for a relationship between pairwise perceptual distances, measured as Euclidian distances in the perceptual space (shown in **Fig. 1**), and pairwise geographic distances, quantified as 1 - Schoener's D indices (*82*). Such a similarity test allows to detect if species perceived as similar tend globally to have similar spatial distributions (**Figs. S21A & S21B**). Furthermore, to test for cases of evolutionary convergence as described in Müller's model, we accounted for the effect of evolutionary relatedness across species by using phylogenetic distances as an additional predictor in the MRM test (**Figs. S21C & S21D**).

We showed that the perceptual distances between pairs of subspecies were significantly correlated with the geographic overlap between their spatial distributions (**Figs. S21A & S21B**; MRM: standardized $\beta_{obs}$ = 0.078, Q95% = 0.040, p = 0.003), even when accounting for the effects of phylogenetic relatedness across taxa (**Figs. S21C & S21D**; MRM: standardized $\beta_{obs}$ = 0.085, Q95% = 0.039, p ≤ 0.001).

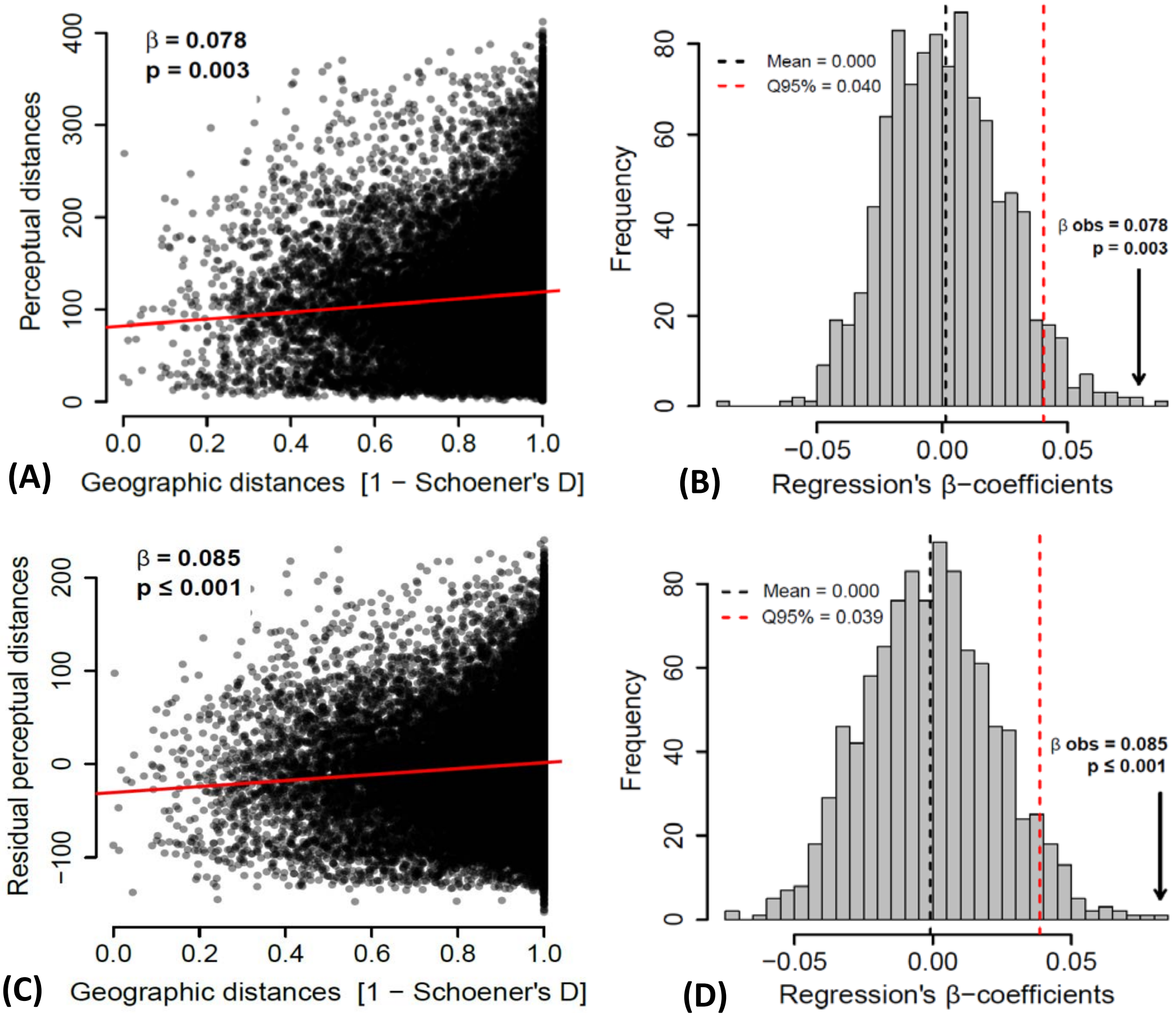

**Figure S21: Pairwise perceptual distances between heliconiine subspecies correlate with geographic distances. (A)** Scatter plot for the similarity test showing the positive correlation between perceptual distances (Euclidean distances in the perceptual space) and geographic distances (1 – Schoener's D of predicted spatial distributions). **(B)** Null distribution of the MRM statistics (β-coefficient of standardized regression) obtained through random permutations of distances across taxa showing a significant positive relationship for the similarity test. **(C)** Scatter plot for the convergence test showing the positive correlation between residual perceptual distances (once accounted for the effect of phylogenetic distances) and geographic distances (1 – Schoener's D of predicted spatial distributions). **(D)** Null distribution of the MRM statistics (β-coefficient of standardized regression) obtained through random permutations of distances across taxa showing a significant positive relationship for the convergence test.

Complementary to analyses based on Schoener's D as a continuous measurement of geographic distances, we designed analyses based on sympatric and allopatric categories for geographic status of pair of subspecies. We defined as sympatric all pairs of subspecies with a Schoener's D higher than 0.2 (thus a geographic distance lower than 0.8). All other pairs were considered allopatric.

First, we tested for differences in perceptual distances across geographic status of pair of heliconiine subspecies. To test for higher perceived similarity between sympatric species than allopatric species, we used non-parametric Wilcox tests to compare perceptual distances across geographic status. In order to account for the effect of phylogeny, we ran the same tests on the residuals of perceptual distances regressed on phylogenetic distances. We computed phylogenetic distances as the pairwise patristic distances on the phylogeny of Heliconiini (*43*) with terminal branches of null length to describe the relative position of subspecies in their associated species.

Second, we designed permutation tests aiming to test for a signal of perceived similarity and convergence higher than expected at random if patterns were distributed randomly along the phylogeny. Thus, we computed the mean perceptual distance (for similarity) and residual perceptual distances (for convergence) in sympatric and allopatric pairs. We compared the observed statistics with a null distribution obtained from random permutation of distances across pairs of subspecies. As such, an observed statistic falling in the lower tail of the null distribution relates to a significant signal of perceived similarity/convergence in the group, and conversely.

As a result, we showed that subspecies in sympatry have significantly lower perceived similarity than allopatric species (Wilcox test: $W = 210 \times 10^6$, $p < 0.001$; **Fig. S22A**), and lower than if patterns were distributed randomly (Permutation test: $D_{obs} = 104.0$, Q5% = 113.1, $p < 0.001$; **Fig. S22B**). Once accounted for evolutionary relationships across subspecies, we showed that subspecies in sympatry have significantly lower perceived residual similarity than allopatric species (Wilcox test: $W = 214 \times 10^6$, $p < 0.001$; **Fig. S22C**), and lower than if patterns were distributed randomly (Permutation test: Dobs = -11.2, Q5% = -3.33, $p < 0.001$; **Fig. S22D**). Altogether, these results comfort our conclusion from the MRM tests based on distances: spatially congruent species show significant degree of pattern similarity that goes beyond expectation from phylogenetic relatedness, hinting for the local convergence of aposematic patterns following Müller's predictions (*3*).

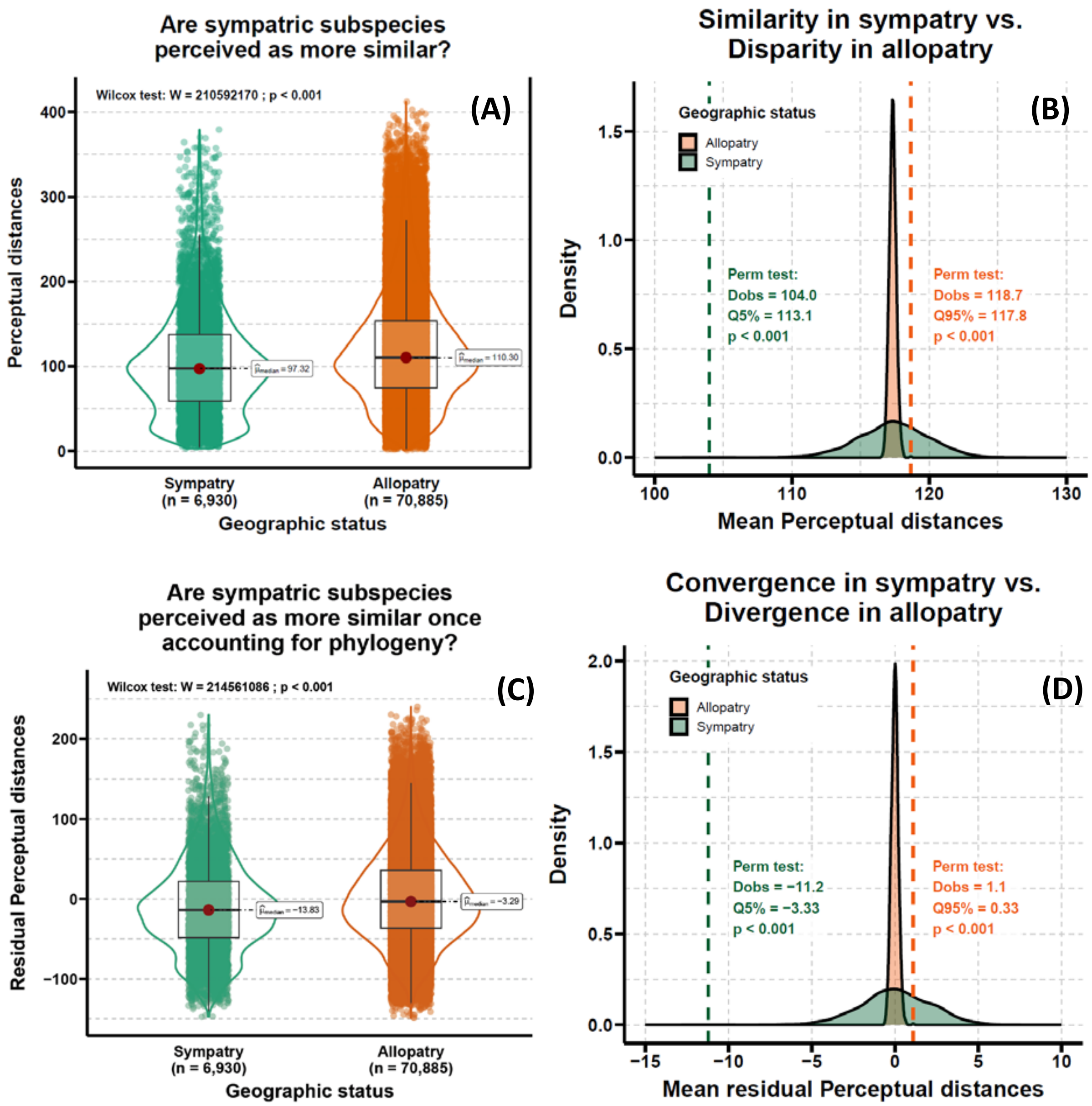


**Figure S22: Tests for similarity (A-B) and convergence (C-D) of perceived wing patterns in sympatry.** **(A)** Comparison of perceptual distances between pairs of subspecies according to their geographic status. **(B)** Null distribution of mean perceptual distances in sympatric and allopatric pairs of species obtained through random permutation of perceptual distances. **(C)** Comparison of residual perceptual distances accounting for phylogeny between pairs of subspecies according to their geographic status. **(D)** Null distribution of mean residual perceptual distances accounting for phylogeny in sympatric and allopatric pairs of species obtained through random permutation of perceptual distances.

#### Appendix 10: Lists of local putative mimicry rings

We built mimetic classifications of wing patterns in local community from GMM clustering applied on different perceptual maps. ‘Local reference’ maps were obtained from the aggregation of similarity triplets provided by an independent panel evaluating only the patterns of local taxa. ‘Extracted’ maps were obtained by extracting coordinates of those same local patterns from the macro-scale perceptual map involving all 432 subspecies patterns. Each cluster represents a putative mimicry ring of co-occurring defended species involved in mutualistic interactions though Müllerian mimicry. We present here the mimicry classifications resulting from such clustering for each local community: Cayenne, French Guyana (**Fig. S23**), Gamboa, Panamá (**Fig. S24**), Jatun Sacha, Napo, Ecuador (**Fig. S25**), Manaus, Amazonas, Brazil (**Fig. S26**), and Santa Teresa, Espírito Santo, Brazil (**Fig. S27**).

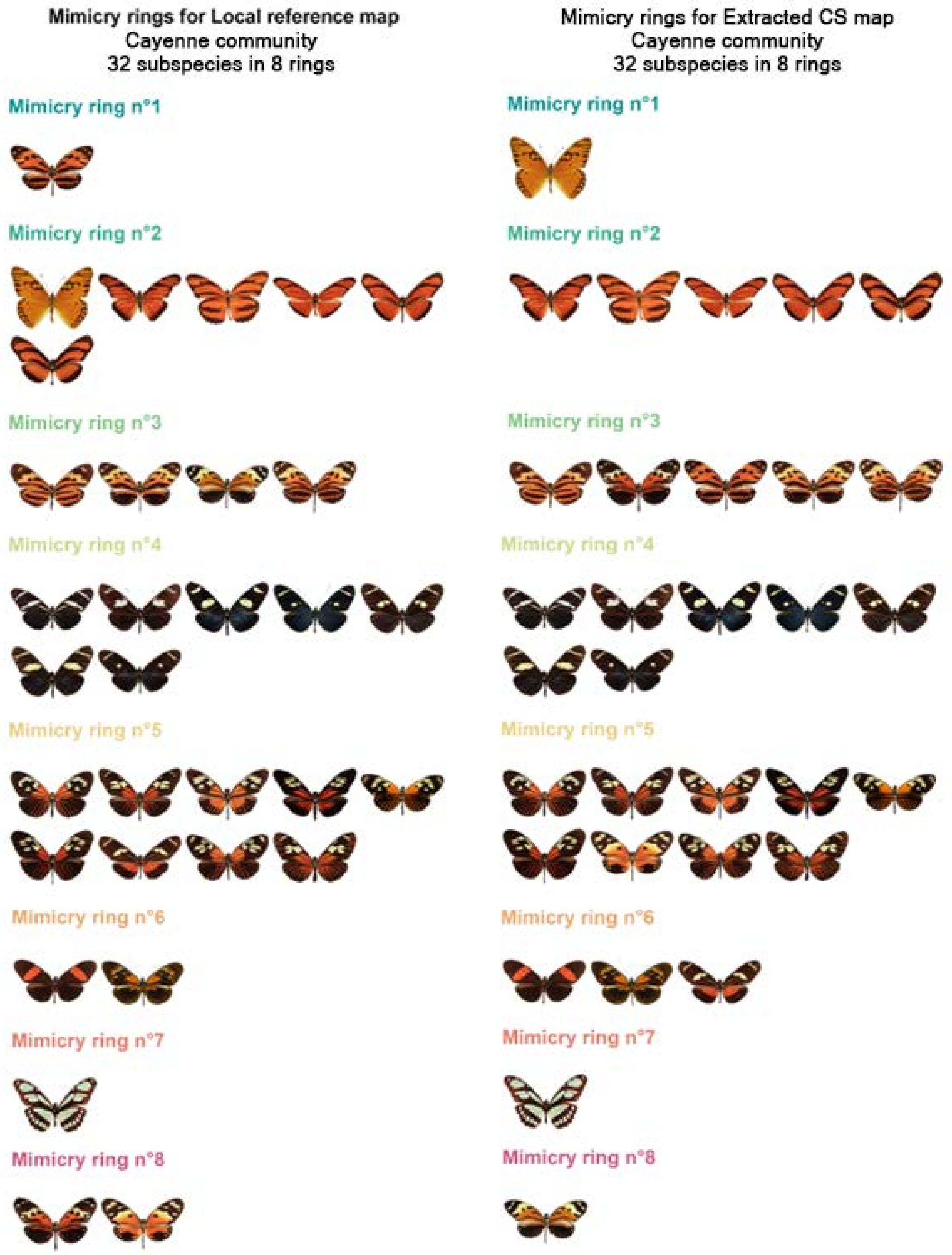


**Figure S23: Visual list of local putative mimicry rings in Cayenne, French Guyana. Left:** From the 'Local reference' perceptual space. **Right:** From the 'extracted' citizen science-based (CS) perceptual space. The eight mimicry rings are ordered by similarity, but they may not be equivalent.

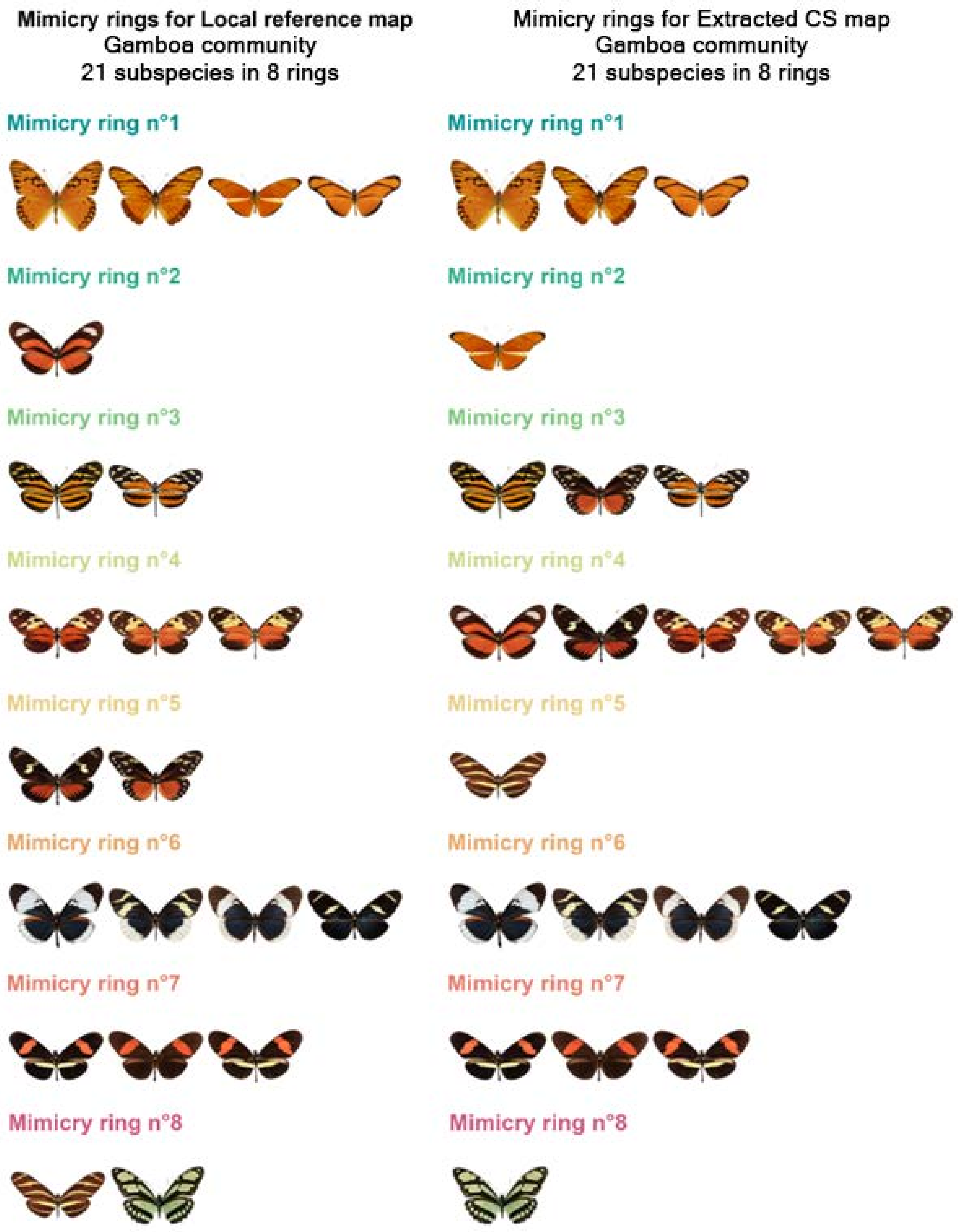


**Figure S24: Visual list of local putative mimicry rings in Gamboa, Panamá. Left:** From the 'Local reference' perceptual space. **Right:** From the 'extracted' citizen science-based (CS) perceptual space. The eight mimicry rings are ordered by similarity, but they may not be equivalent.

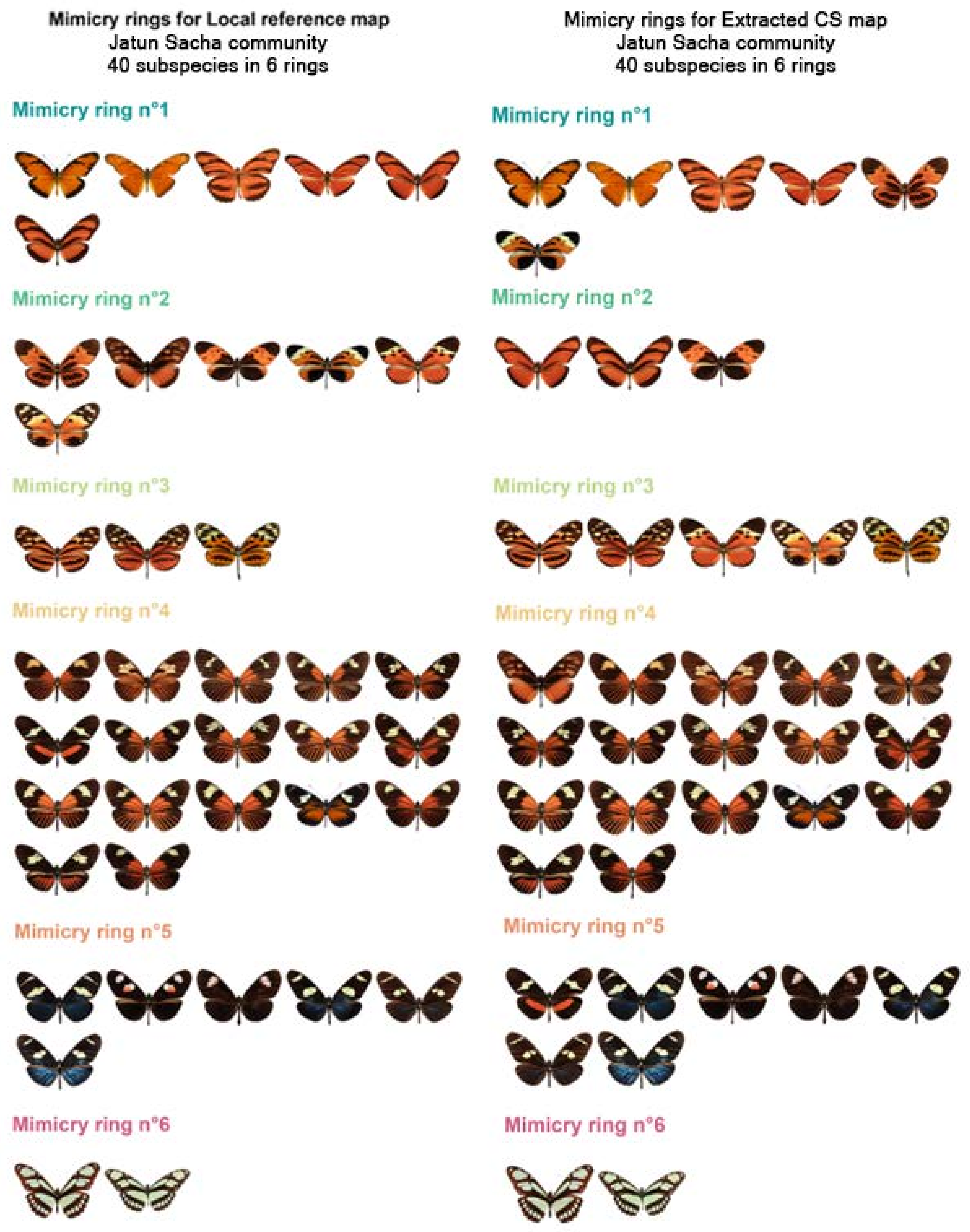


**Figure S25: Visual list of local putative mimicry rings in Jatun Sacha, Napo, Ecuador. Left:** From the 'Local reference' perceptual space. **Right:** From the 'extracted' citizen science-based (CS) perceptual space. The six mimicry rings are ordered by similarity, but they may not be equivalent.

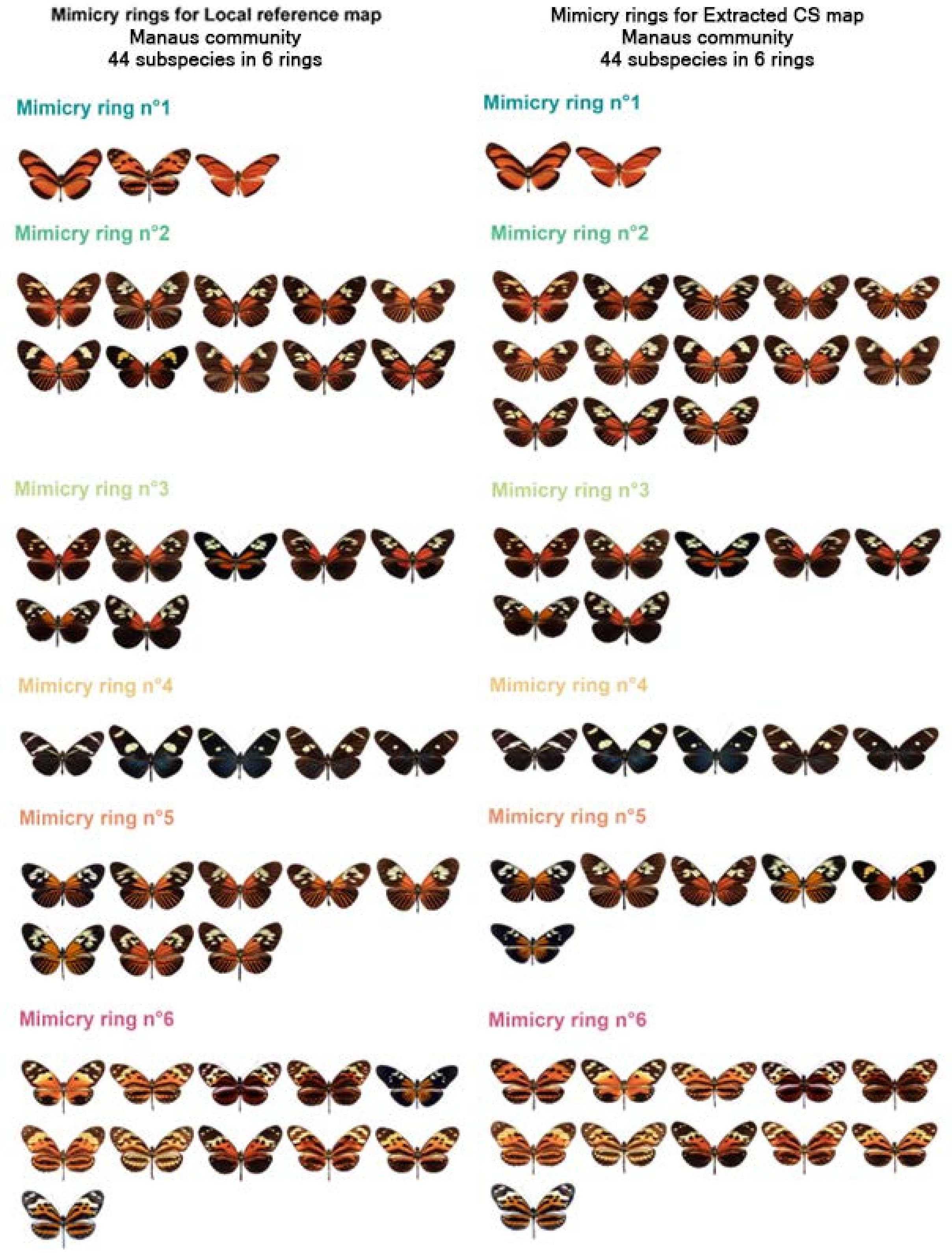


**Figure S26: Visual list of local putative mimicry rings in Manaus, Amazonas, Brazil. Left:** From the 'Local reference' perceptual space. **Right:** From the 'extracted' citizen science-based (CS) perceptual space. The six mimicry rings are ordered by similarity, but they may not be equivalent.

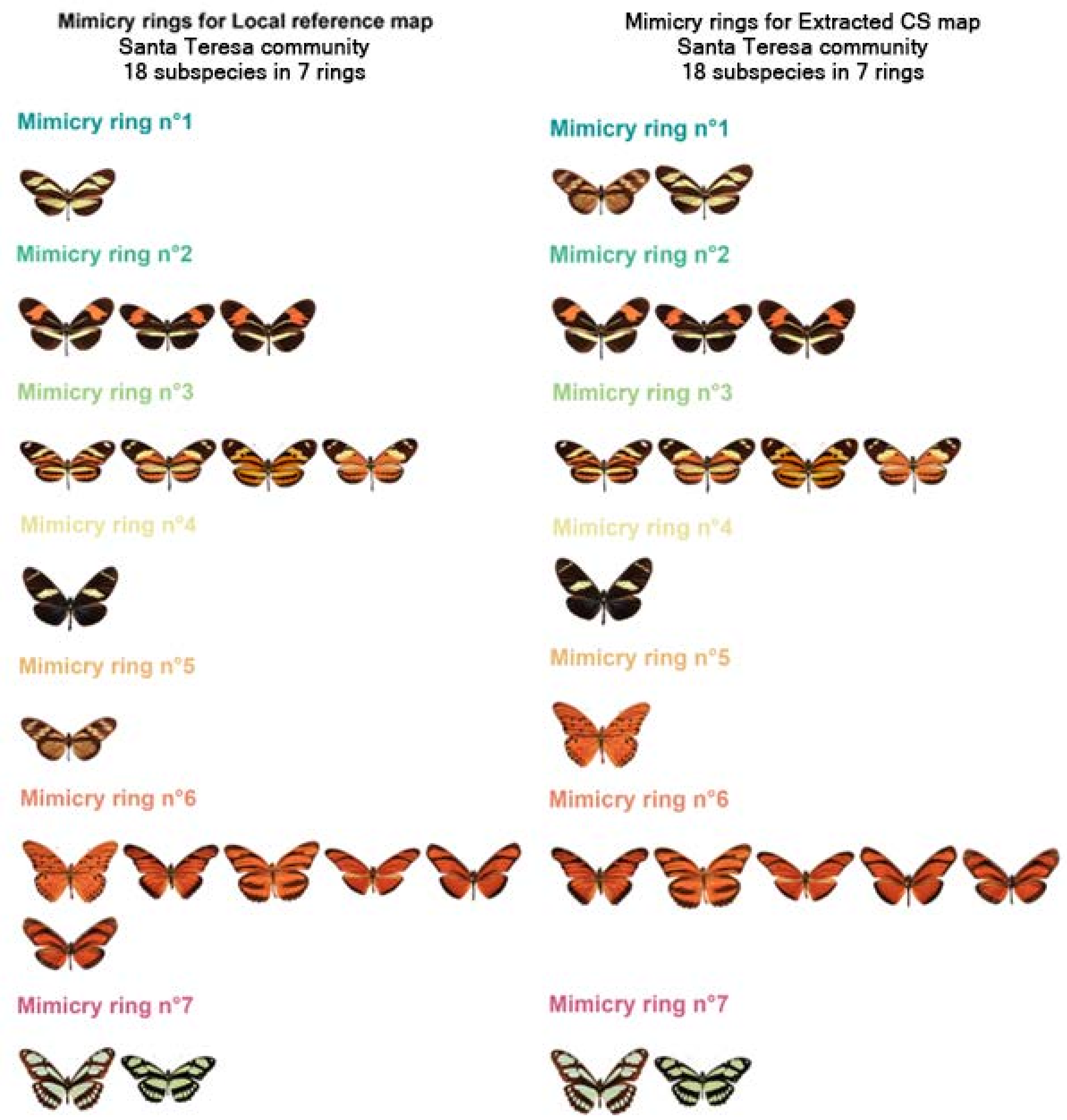


**Figure S27: Visual list of local putative mimicry rings in Santa Teresa, Espírito Santo, Brazil. Left:** From the 'Local reference' perceptual space. **Right:** From the 'extracted' citizen science-based (CS) perceptual space. The seven mimicry rings are ordered by similarity, but they may not be equivalent.
